# Gap genes are involved in inviability in hybrids between *Drosophila melanogaster and D. santomea*

**DOI:** 10.1101/2021.12.06.471493

**Authors:** Daniel R. Matute

**Affiliations:** Biology Department, University of North Carolina, Chapel Hill, North Carolina

**Keywords:** Hybridization, gap genes, *Drosophila*

## Abstract

Evolved changes within species can lead to the loss of viability in hybrids. Inviability is also a convenient phenotype to genetically map and validate functionally divergent genes and pathways differentiating closely related species. Here, I identify the *Drosophila melanogaster* form of the highly conserved essential gap gene *giant* (*gt*) as a key genetic determinant of inviability in hybrids with *D. santomea*. The presence of this allele is sufficient and necessary to cause a prevalent abdominal ablation in the developing hybrid embryo. Further genetic analysis indicates that *tailless* (*tll*), another gap gene, is also involved in the hybrid defects. *giant* and *tll* are both members of the gap gene network of transcription factors that participate in establishing anterior-posterior specification of the dipteran embryo, a highly conserved developmental process. I found no evidence for positive selection in either of the two genes. Genes whose outputs in this process are functionally conserved nevertheless evolve over short timescales to cause inviability in hybrids.

**SIGNIFICANCE STATEMENT:** Hybrid organisms—which have parents from different species—are usually less healthy than pure-species organisms. Hybrid inviability is an extreme example of this phenomenon. Defects result from incompatibilities between genes that have diverged in the parental species. In this piece, I dissect the genetic basis of hybrid inviability between two species of *Drosophila*: *D. melanogaster* and *D. santomea*. I identified two genes, *gt* and *tll* responsible for the inviability in these hybrids. The two genes belong to the gap gene network, a highly conserved pathway that is in charge of establishing embryonic polarity in insects. These results show that developmentally important genes might evolve sufficiently fast to cause hybrid inviability in species that still can interbreed.

## INTRODUCTION

The formation and persistence of species involves the buildup of barriers to gene flow as genome divergence accrues over time. These genetic barriers arise as species differentiate and involve breakdowns in a variety of cellular, developmental, and behavioral processes; eventually these barriers lead to reduced fitness of hybrids relative to pure species (1, 2). Hybrid inviability (HI), the condition in which interspecific hybrids do not achieve adulthood because of developmental defects, is one of these barriers. The question of how natural selection could allow such maladaptive and extreme phenotypes has been a subject of intense interest to evolutionary biologists and developmental geneticists alike (3–5).

Dobzhansky (6) and Muller (7) formulated a widely regarded genetic model in which hybrid defects, including HI, arise as a collateral effect of evolutionary divergence between populations that acquire incompatible changes in interacting loci, or Dobzhansky-Muller incompatibilities (DMIs; (Dobzhansky 1937; Muller 1942)). Because the divergent alleles at the DMIs loci only have fitness costs when they are forced together in hybrids, natural selection does not oppose the changes in each species. There is substantial evidence in support of the DM model (8), including nearly a dozen instances in which HI alleles have been identified to the gene level (9–19). Some of these alleles have been shown to evolve through positive selection (10–12, 14, 20) while others show no clear signature of selection (21). The variety of both the type of genes involved in HI and the processes that drive allelic divergence indicates that HI can occur in a multitude of ways (22).

Developmental processes are generally guided by interacting regulatory genes and elements, making them a rich source of potential candidates for HI. Developmental processes and their outputs are often deeply conserved phylogenetically, often displaying conserved functional attributes (reviewed in (23)). Large-effect mutations to developmental regulators are often incompatible with life, and these genes tend to be evolutionarily conserved both in sequence and phenotypic output (24–27). In some instances, alleles involved in developmental specification show evidence of rapid evolution (28). While the developmental phenotypes in which these genes are involved generally remain similar across species, the genetic underpinnings of these crucial phenotypes may evolve (29–33), and if their pace of functional evolution is sufficiently fast, could contribute to HI. The question arises, then, as to whether they evolve functionally at a sufficient pace to become incompatible in hybrids.

Two different lines of evidence elevate this possibility and suggests that developmental genes, and the processes they direct, could contribute to HI. First, several cases of embryonic hybrid lethality have been identified in *Drosophila*: female HI in hybrids between *D. montana* females and *D. texana* males (34), female lethality in hybrids of *D. melanogaster* females and *D. simulans* males (35–37); and male embryonic lethality in hybrids of *D. melanogaster* females and *D. santomea* males ((38), see below). While a multitude of processes can lead to this phenomenon, alleles involved in early development might play an important role in this dysfunction. Second, genetic dissection of conserved regulatory pathways regulating *Drosophila* development showed that, contrary to expectations, segmentation genes diverge functionally at a rapid and continuous pace in the genus, as evidenced by the non-equivalency of regulatory elements in control transgenic replacements (39).

These results pose the possibility that even for developmental phenotypes that remain similar across phylogeny, their genetic underpinnings change occasionally in substantial ways (29–33). Referred to as developmental systems drift —functional divergence of genes in developmental regulatory pathways with conserved outputs — has also been documented for nematode vulva induction (40, 41), and sex determination in frogs (42). Developmental systems drift has also been proposed to lead to hybrid defects (43). If genetic changes occur in different directions in two species, their hybrids might not have a functional pathway to produce the required developmental phenotype. This is a simple—but to date unsubstantiated— way to explain HI.

Crosses between *D. melanogaster* females and *D. santomea* males produce sterile, adult females, even though the two species diverged over 10 mya (Ks = 24%, (44–46)). The hybrid male embryos die during embryogenesis with a striking abdominal ablation phenotype (38, 47). Here, I employ fine duplication and deficiency mapping to identify the *mel* allele of the highly conserved gap gene *giant* (*gt*) as a key genetic determinant of this phenotype and consequent HI in these crosses (Figure S1). This result is surprising because *giant* is a key developmental regulator involved in the anterior-posterior specification of the dipteran embryos whose role in anterior-posterior specification is conserved over 250 million years (44, 46, 48). Out of roughly 13 *X*-linked loci involved in HI between *D. melanogaster* and *D. santomea* (47), *D. melanogaster’s giant* (*gt^mel^*) is necessary and sufficient to cause the characteristic abdominal ablation phenotype seen in *mel/san* hybrids. Additional gene mapping revealed that *tailless* (*tll*), another gap gene is also involved in HI in *mel/san* hybrids, and that the deletion of *D. melanogaster tll* improves embryonic viability but only in the absence of *gt^mel^*. The results show here indicate that even genes with crucial developmental roles that are conserved over vast evolutionary time scales can be involved in HI.

## RESULTS

In the cross between *D. melanogaster* females and *D. santomea* males*, mel/san* hybrid males inherit a *X_mel_* chromosome and a *Y* chromosome from *san* (*Y_san_*). Hybrid females inherit a *X_mel_* and *X*-chromosome from *D. santomea* (*X_san_*). Although both sexes show abdominal ablations, these phenomenon is more common in males (38, 47). I first explored whether *Drosophila* hybrids other than *mel/san* also showed embryonic HI and abdominal ablations. I examined the embryonic lethality rates and associated cuticular phenotypes from hybrid crosses between various species within the *melanogaster* supercomplex and species of the *yakuba* clade (Figure 1, Figure S1). Hybrid embryos between *D. santomea* and the other species in the *yakuba* clade — *D. teissieri* and *D. yakuba* — are mostly viable and showed no abdominal ablations in any of the six possible reciprocal crosses (Table S1; (38, 49–51)). Embryonic HI is rare in interspecific crosses between species of the *yakuba* complex (but see (51)). Hybrid inviability is also non-existent in hybrids between collections of species of the *simulans* species group —

**FIGURE 1.**
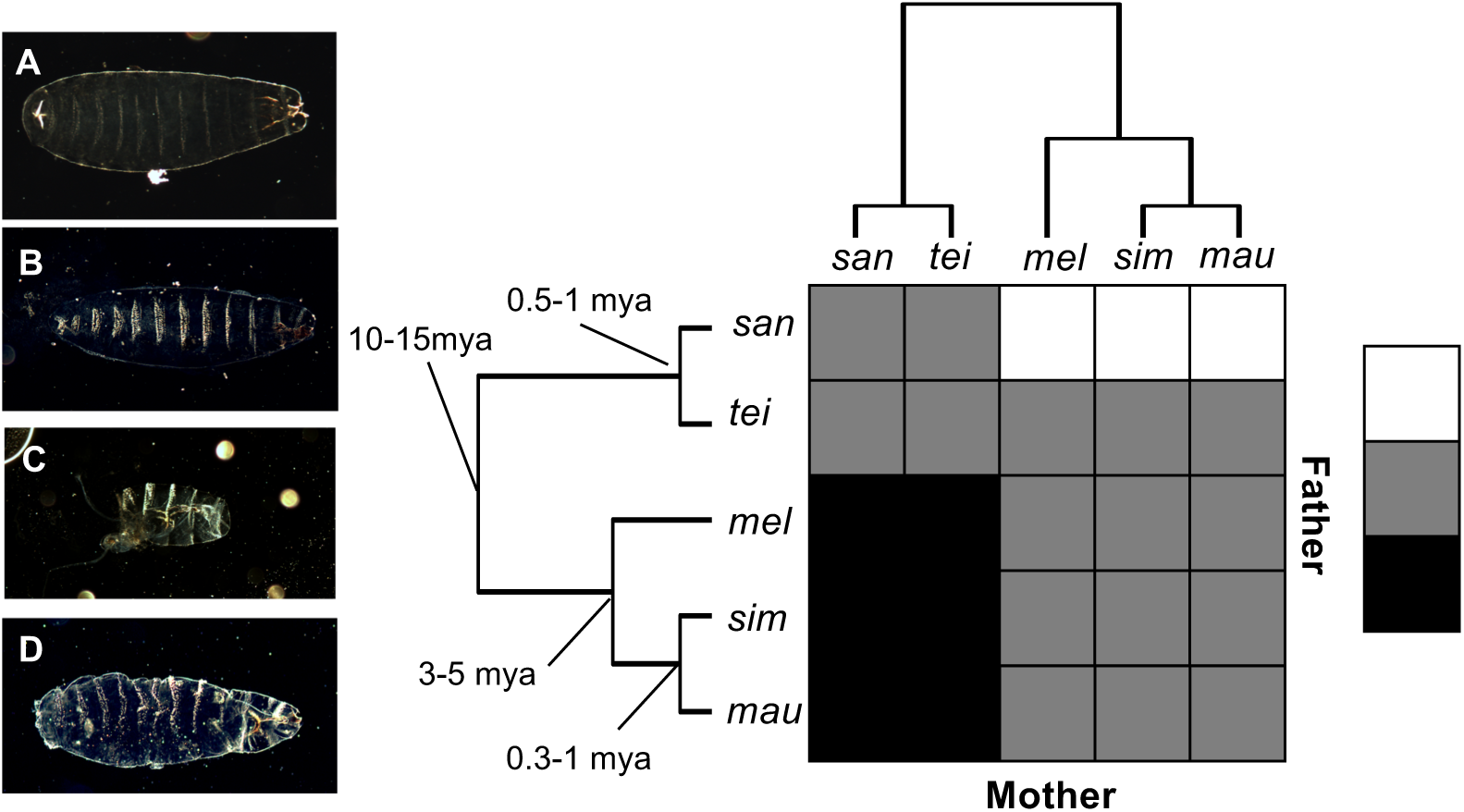
All the *X* chromosomes from the *mel* supercomplex cause inviability in *mel/san* hybrids but not in *mel/tei* hybrids. Unlike pure species (**A**. *D. santomea*; **B**. *D. melanogaster*), F1 *mel/san* hybrid males show abdominal ablations (**C**). The nature of the defect in these hybrid males is identical to that seen in *sim/san* and *mau/san* hybrid males. Females from the same cross also show such ablations but more rarely and the majority of dead embryos show a complete abdomen (Figure S12). Hybrids between females from the *melanogaster* supercomplex and *D. teissieri* males show little embryonic lethality and among the few dead embryos there are no abdominal ablations (e.g., **D.** *mel/tei* hybrid male). The viability of each genotype is shown in Figure S1. **E**. Phylogenetic relationships between the species of the *melanogaster* species subgroup. The heatmap shows the relative occurrence of abdominal ablations in hybrid males. White: common, grey: absent. Black: pairs with complete behavioral isolation.

*D. simulans* (*sim*), *D. mauritiana* (*mau*) and *D. sechellia* (*sec*) (Figure 1, (50)). The embryonic viability of male *mel/sim* and *mel/mau* hybrids is high in all cases (Table S2, (10, 37, 52, 53)). The few rare embryos that failed to develop and hatch showed no abdominal ablations (Table S2).

Crosses between females of two species of the *sim* clade (*sim* and *mau*) and *san* males showed high levels of hybrid inviability, especially of males which carry the *X*-chromosome from sim or mau (*X_sim_* and *X_mau_*, respectively; Figure S1). These dead hybrids show the characteristic abdominal ablation observed in *mel/san* hybrids. The genetic changes that ultimately lead to abdominal ablations in hybrids with *D. santomea* must have occurred before the split of the three species (*mel*, *sim* and *mau*), approximately five million years ago (44–46).

Manipulation of chromosomal inheritance using *mel* attached-*X* chromosomes (Figure S2), produces hybrid F_1_ males that inherit a *X_san_* and a *D. melanogaster Y* chromosome (*Y_mel_*), hybrid F_1_ females (*X_mel_*/*X_san_*), and a few rare metafemales (*X_mel_*/*X_mel_*/*X_san_*). The abdominal ablation is much more common in carriers of the *X_mel_* chromosome (Figure S1). Similar to crosses between *mel* attached-*X* females and *san* males, crosses between attached-*X sim* (and *mau*) to *san* males produce male hybrid progeny (*X_san_*/*Y_sim_*, and *X_san_*/*Y_mau_*) that is viable but hybrid females (*X_sim_X_sim_*/*Y_san_* and *X_sim_X_sim_X_san_* or *X_mau_X_mau_*/*Y_san_* and *X_mau_X_mau_X_san_*) that are inviable. None of the embryos resulting from crosses between attached-*X D. yakuba,* or attached-*X D. santomea* showed abdominal ablations. These genetic analyses suggest that similarly to *X_mel_*, the *Xsim* and *X_mau_* harbor a locus involved in HI in hybrids with *D. santomea*. I next identified the genetic locus that causes HI by abdominal ablation using genetic tools available in *D. melanogaster*.

### Genetic mapping shows *giant* is involved in *mel/san* HI

To identify the *X*-linked allele involved in HI, I did a genome-wide association study using a panel of inbred *D. melanogaster* lines (i.e., DGRP, (54, 55)) and studied whether any genetic variant segregating in *D. melanogaster* was associated with total strength of HI with *D. santomea*. In all crosses, the hybrid males die, but the females show differential rates of survival. I found a strong association between a 16.5 kb haplotype in the *X*-chromosome and high levels of HI (Figure 2A). This haplotype is delimited by *CG32797* and *boi*, includes *gt*, and overlaps with a segment of the *D. melanogaster X*-chromosome (*X_mel_*) previously associated with male HI (47). I also found an association between the magnitude of HI and a *X*-linked haplotype containing a single gene, *CG12496*. Six autosomal genes were associated with inviability were distributed along the *mel* genome (2L: *CG13284*, *CG31810*; 2R: *CG10936*, *CG9313*, 3R: *Glut4EF*; 3L: *CG32365*).

**FIGURE 2.**
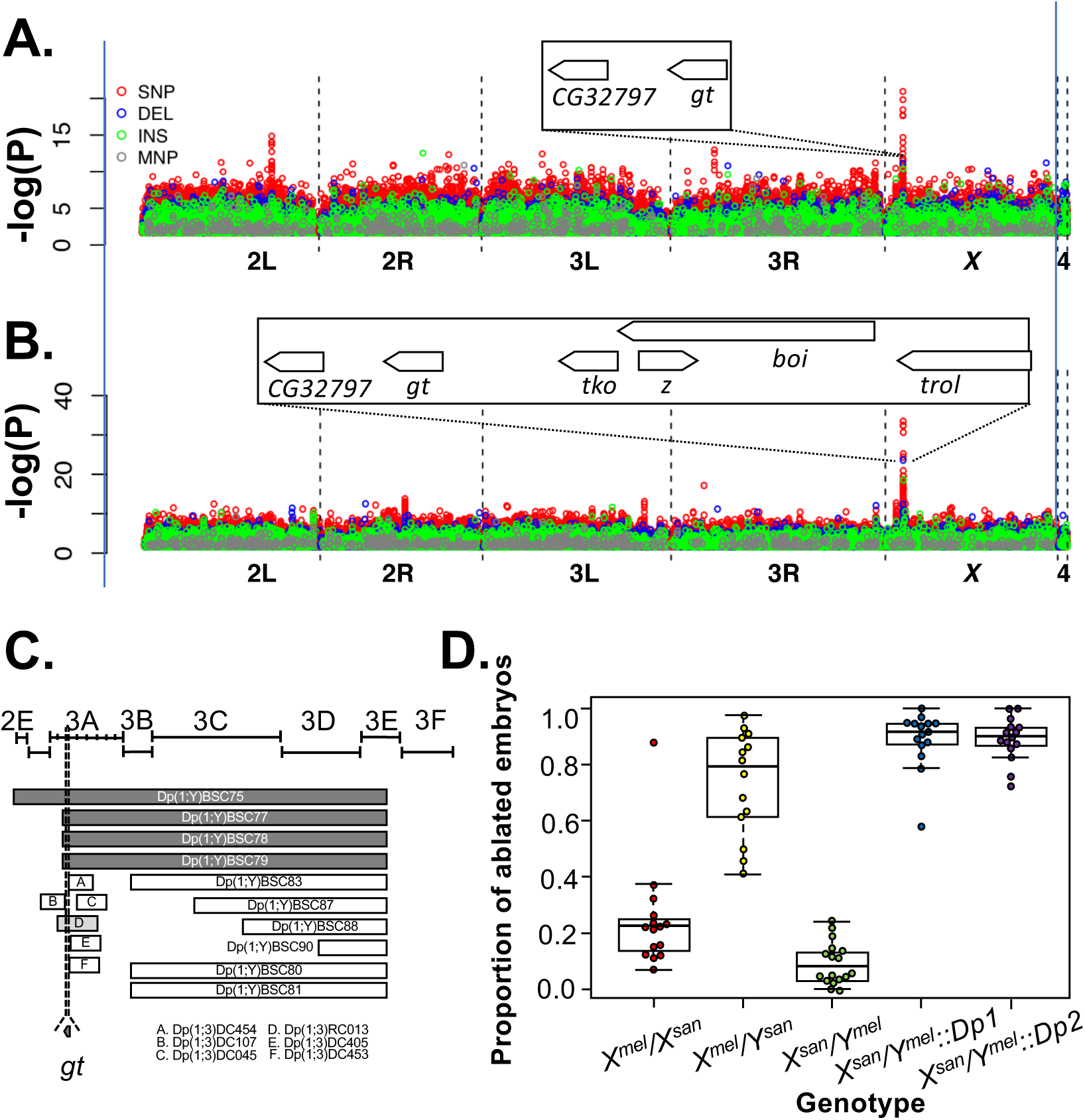
A *D. melanogaster X*-chromosome haplotype that encompasses *gt^mel^* is associated with hybrid inviability and abdominal ablations in ♀ *D. melanogaster*/♂ *D. santomea* hybrid males. **A.** Genome-wide association study of the genetic causes of hybrid inviability in *mel/san* hybrids (both sexes pooled) using segregating variation within *D. melanogaster*. A haplotype of 54kb in the tip of the *X*-chromosome is strongly associated with the presence of abdominal ablations. The haplotype harbors six genes: *CG32797, gt, tko, boi, z,* and *trol.* **B.** Genome-wide association study of the genetic causes of abdominal ablations in *mel/san* hybrids (both sexes pooled) using segregating variation within *D. melanogaster*. A haplotype of 54kb in the tip of the *X*-chromosome is strongly associated with the presence of abdominal ablations. The haplotype harbors six genes: *CG32797, gt, tko, boi, z,* and *trol.* Green: insertions, blue: deletions, red: SNPs, purple: multinucleotide polymorphisms. Green: insertions (INS), blue: deletions (DEL), red: SNPs, purple: multinucleotide polymorphisms (MNP). **C.** I introduced small *X^mel^* pieces attached to *Y_mel_* to identify *X_mel_*-linked alleles that cause hybrid inviability in in *mel/san* hybrids males. For all *Dp(1;Y)* duplications, I evaluated at least 50 embryos per cross were for viability. For *Dp(1;3)* duplications, I evaluate between 20-56 embryos as *C(1) DX, Dp(1;3)/ Dp(1;3)* are weak. I narrowed down the allele that causes HI to an interval of *X^mel^* comprising 3A3 which only contains *giant*. White bars show duplications with no abdominal ablations. The light grey bar shows a duplication with a moderate rate of abdominal ablations; dark grey show duplications with high levels of abdominal ablations. **D.** Relative frequency of abdominal defects in five different hybrid genotypes from *D. melanogaster* and *D. santomea* crosses. Each point is a replicate of at least 100 embryos. Pure species embryos show no abdominal defects and show little embryonic lethality. *mel/san* hybrid males (*X^mel^/Y^san^*) frequently show a lethal characteristic abdominal ablation (yellow points) that is also present in some *mel/san* hybrid females (red points). The reciprocal *mel/san* hybrid males (*X^san^/Y^mel^*) routinely survive and the few embryos who die do not show abdominal ablations (green points). *X^san^/Y^mel^*males carrying X-Y translocations [i.e., *Dp(1;Y)* in blue and *Dp(1;Y)* in purple] that harbor *gt^mel^* also show this lethal ablation. Dp1: *Dp(1;Y)BSC78* (stock 29802); Dps: *Dp(1;Y)BSC79* (stock 29803). A map showing the frequency of abdominal defects caused by multiple *X^mel^*translocation is shown in Figure S3.

A second GWAS for the incidence of abdominal ablations (Figure 2B) showed the frequency of abdominal ablations in *mel/san* hybrids (both sexes pooled) is associated with an *X_mel_* haplotype that contains six genes (*CG32797, gt, tko, boi, z,* and *trol*; Figure 2B). This interval also overlaps with the region associated with the total strength of HI (described immediately above). Four more *X*-linked genes are associated with abdominal ablations: *CG12496*, *inc*, *Nrg*, and *CG2685*. Eleven autosomal genes were associated with the frequency of ablations in hybrids (2L: *CG32982*, *dnt*; 2R: *CG7544*, *CG8089*, *nompA*, *Pka-R23R*; 3L: *siz*, *CG11537*, *ckd*, *Lmpt*; 3R: *CG43462*). The intersection of these two GWASs revealed that a single *X*-linked block—in the 3A cytological band of the *mel* genome—is associated with total HI and with the frequency of abdominal ablations. This chromosomal segment contains two genes: *gt* and *CG12496*. These results are in line with previous efforts that have shown that this region in the 3A region is responsible for abdominal ablation and *mel/san* male HI (47).

Next, and to refine the region of the *X_mel_* chromosome carrying the determinant of the abdominal ablation phenotype, I introduced small segments of *X_mel_* (containing *mel* alleles ranging from approximately 10 to 100 genes each; as in (47, 56, 57) into the genetic makeup of *X_san_/Y_mel_* hybrid F_1_ males (crossing scheme shown in Figure S2; fly stocks listed in Table S3). Measuring the rates of hybrid embryonic lethality in the presence of nested *Dp(1;Y)* and *Dp(1:3)* duplications of the *X_mel_* chromosome allowed me to refine the genomic interval involved in HI to the region encompassing the cytological region 3A3 (dmel6: 2,410,000-2,580,000; (58)). Male *mel/san* hybrid embryos harboring *X_san_* and *X_mel_* duplications containing the 3A3 portion fail to hatch (Figure 2C). They also show the striking abdominal ablation common in *mel/san* hybrid males carrying the full-length *X_mel_* and *Y_san_* (Figure 2D, Figure S3). Notably, the overlapping region of the duplications that cause inviability and the ablation contains only one gene: *giant* (Figure 2C, Figure S3). The ablation phenotype is not found in the presence of other lethality-inducing fragments from elsewhere on the *X_mel_* chromosome (as found in this work and (47)). This mapping rules out the potential involvement of *CG12496* in the HI of *mel/san* hybrids by abdominal ablation. *gt_mel_* caused HI with abdominal ablation in hybrids with all examined lines from *D. santomea* confirming that the interaction is not a line specific effect (Figure S4A; lines listed in Table S4). These experiments suggest that introducing a *gt_mel_* allele in a *X_san_*/*Y_mel_* male hybrid background is sufficient to cause lethality with abdominal ablation in hybrids derived from multiple *D. santomea* fathers. The comparison of *Dp(1;Y)* and *Dp(1;3)* duplications has two caveats worth noting. First, pure species *mel C(1)DX, Dp(1;3)/Dp(1;3)* females are generally weaker than *C(1)DX, Dp(1;Y)* and lay fewer eggs in conspecific and heterospecific crosses. I thus had lower power in crosses involving the former type of females (Table S5). Second, the rate of complementation of *Dp(1;3)* and *Dp(1;Y)* is not identical (Table S6) suggesting the existence of position effects.

I also did similar genetic crosses using *Dp(1;Y)* and *Dp(1;3)* duplications to assess whether the same *X*-chromosome segments (and therefore *gt_mel_*) caused HI in crosses with four additional species, two from the *simulans* species complex (*mau* and *sim*), one from the *yakuba* species complex, *D. teissieri,* and *mel* itself. In no other interspecific hybrids did *gt_mel_* have a deleterious effect in the form of reduced viability suggesting that the hybrid lethal effects *gt_mel_* are exclusive to *mel/san* hybrids (Figures S5-S6). In all these crosses, the number of dead embryos was low (i.e., fewer than 4 embryos for each cross type), but none of them showed abdominal ablations.

### The *gt_mel_* allele causes inviability in both male and female hybrids

The crosses described above address the effect of the presence of a *gt_mel_* allele on HI in *mel/san* hybrids. Next, I studied the absence of a functional a *gt_mel_* allele in both sexes using a *gt_mel_* null-allele, *gt_mel_^X11^* (59, 60). First, I tested whether the absence of *gt_mel_* had a deleterious effect on hybrid females by scoring whether any allele on *X_mel_* between the cytological positions 2F1 and 3A4 affected the viability of *mel/san* hybrid females. I used *mel* chromosomal deficiencies and scored the number of *FM7/san* hybrid females compared to their *df(i)/san* sisters ((10, 61); see Methods). Deviations from 1:1 expected ratio indicate the presence of alleles involved in HI. If *FM7/san* hybrids survive at a higher rate than *df(i)/san,* then the uncovered *san* segment contains a recessive allele involved in HI. If *FM7/san* hybrids survive at a lower rate than *df(i)/san* hybrids, then the absent *mel* segment contains a dominant (or semi-dominant) allele involved in HI. The initial screening using the line *san* Carv2015.16 showed that hybrid females with a deletion for the *X_mel_* region between 3A2 and 3A3 (which contains *gt_mel_*) are more viable than hybrid females that carry the balancer chromosome with *gt_mel_* (*df/san* to *FM7/san* ratio ≈ 2:1; Figure 3A). The minimal interval that harbors the female HI determinant contains six genes, one of them being *gt*.

**FIGURE 3.**
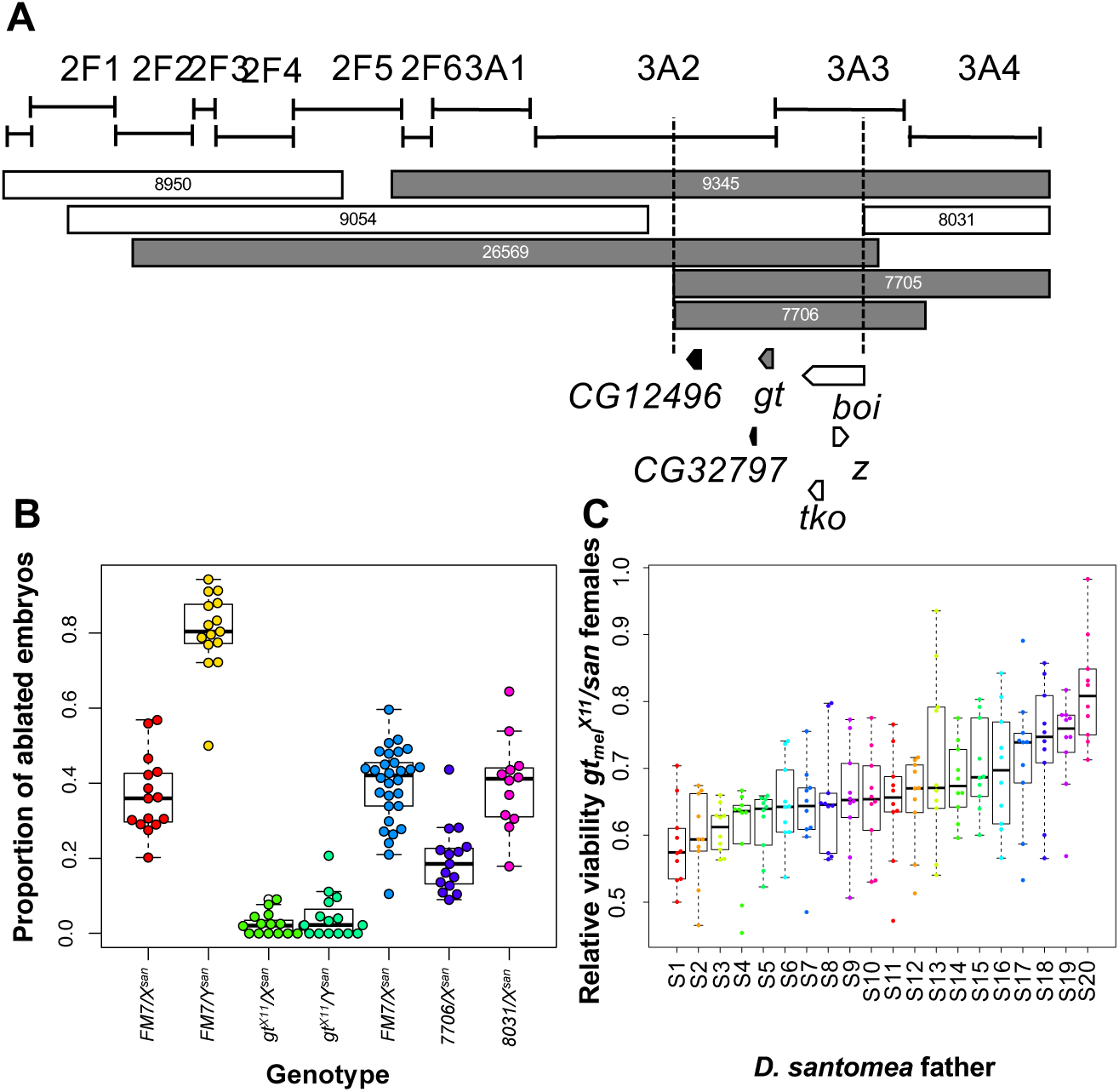
*giant^mel^*causes HI and abdominal ablations in *mel/san* females. **A**. Deficiency mapping and null-allele mapping revealed that *giant* also causes hybrid inviability in hybrid females. Grey: deficiencies that increase viability of *mel/san* F1 hybrid females (Significant linear contrasts, P < 0.05 after multiple comparison corrections). White: deficiencies that do not increase viability. **B.** When females fail to hatch, it is not uncommon for them to be abdominally ablated. The presence of *gt_mel_* increases the frequency of abdominal ablations. **C.** Relative *gt_mel_^X11^/X_san_* female viability (i.e., proportion of *gt_mel_^X11^* carriers in hybrid crosses) in twenty *D. santomea* isofemale lines. Boxes in the boxplot are ordinated from the lower median (left) to the highest (right). S1: SYN2005; S2: sanCAR1490; S3: sanCOST1250.5; S4: sanCOST1270.1; S5: sanOBAT1200; S6: sanOBAT1200.2; S7: sanRain39; S8: sanCAR1600.3; S9: Carv2015.1; S10: Carv2015.5; S11: Carv2015.11; S12: Carv2015.16; S13: Pico1680.1; S14: Pico1659.2; S15: Pico1659.3; S16: Amelia2015.1; S17: Amelia2015.6; S18: Amelia2015.12; S19: A1200.7; S20: Rain42.

I further refined the genetic analysis of this region by testing null alleles of the genes in the interval. Of the four genetically characterized genes in the mapped interval, 3A3, only mutants of *gt_mel_* lead to an increase of female hybrid viability (Table S7). Females that carry *gt_mel_* (*FM7::GFP/san*) emerged at a lesser frequency than their null allele-carrying sisters (*gt_mel_^x11^/san* Figure 3B, Table S7). In contrast, male hybrid embryos carrying a *X_mel_* chromosome and null mutations of the genes adjacent to *gt* (*boi*, *trol*, and *tko*) show abdominal ablations (Figure S7).This difference in viability holds when other *X*-chromosome balancers are used as well (Figure S8).

The abdominal ablation defect that is characteristic of *mel/san* males is also present in some proportion of *mel/san* female embryos that die (38, 47). I next tested whether *gt_mel_* causes abdominal ablations in female as it does in male hybrids. *gt_mel_-*carrying females (*FM7::GFP/san;* Figure 3B) have abdominal ablations more frequently than their *gt_mel_^x11^*/*Xsan* sisters (Figure 3B; *t*-test comparing the frequency of ablations in the two genotypes: *t* = 6.853, df = 16.147, P = 3.699 × 10^-6^). These results indicate that *gt_mel_*, the primary genetic determinant of the abdominal ablation in male *mel/san* hybrids, is sufficient to render some hybrid females inviable by inducing abdominal ablations. Two genes adjacent to *gt*, *CG32797* and *CG12496* have no available mutant stocks. The former, *CG32797*, is not expressed in embryos (62, 63) and is an unlikely candidate to cause embryonic inviability in *mel/san* hybrids. *CG12496* is expressed in the early embryos (2-14 hours after egg laying, (62, 63)), so role in HI cannot be fully excluded. However, the results for the *gt_mel_* allele explain a large proportion of the inviability and abdominal ablation phenotypes I observe with the larger deletion (Figure 3B, Figure S9). Notably, none of the deficiencies has any effect on viability in any other hybrids (i.e., *mel/sim*, *mel/mau* and *mel/tei*) or in within-species *mel* F1s (Figures S10 and S11, respectively). Taken together, the mapping efforts are consistent and reveal that the *gt^mel^* allele is necessary and sufficient to cause abdominal ablation defects in both male and female in *mel/san* F1s hybrids and not in any other hybrid genotypes.

Female HI varies across *san* lines, however, ranging from little inviability (e.g., *san* SYN2005, *df/san* to *FM7/san* ratio ≈ 1:1; Figure 3C) to almost complete inviability (e.g., *san* Rain42; *df/san* to *FM7/san* ratio ≈ 4:1; Figure 3C, Figure S4B).

Hybrid male embryos carrying *gt^X11^* were less likely to be abdominally ablated compared to other *X^mel^/Y^san^* hybrids (mean number *of FM7/Y^san^* ablated embryos: 80.33%, Figure 3B; mean proportion *of gt_mel_^X11^/Y^san^* ablated embryos: 4.41%, Figure 3B; t-test comparing the frequency of ablations in *FM7/Y^san^* males and *gt_mel_^X11^/Y_san_* males: t = 23.972, df = 21.614, P < 1 × 10^-10^), and instead show other developmental defects (Figure S12). These experiments demonstrate that *gt_mel_* is necessary for the abdominal ablation in *mel/san* hybrid males. These *X_mel_/Y_san_* males with a null allele of *gt_mel_* (i.e., *gt_mel_^X11^/Y_san_*) do not show increased viability. This result is not surprising for two results. First, these males have no functional copy of *gt*, an essential gene. Second, *X_mel_* harbors at least eight more dominant (or semidominant) factors that also cause embryonic inviability (47), which may be specifically lethal to *mel/san* hybrids.

#### tll_mel_ exacerbates the hybrid inviability caused by gt_mel_

Hybrid defects are usually caused by interacting genetic elements (reviewed in (8, 64)). *giant* is an essential factor in the gap gene regulatory network, a set of interacting genes expressed in the blastoderm embryo to establish anterior-posterior patterning (60, 65–67); its function in segmentation as a transcriptional repressor of other gap genes (*Kruppel* and *knirps*; (67–69)) is conserved in arthropods. *giant* is itself repressed by the gene products of *hunchback* (66, 67, 70), *tailless* (71), and *hucklebein* (72). The proteins *Caudal* (73) and *Bicoid* (66, 74) activate *gt*, which localizes to two broad stripes, one towards the anterior and one towards the posterior pole of the embryo (reviewed in (75)). Given this knowledge, I hypothesized that gap genes interacting with *gt_mel_* could be additional candidates contributing to inviability in *mel/san* hybrids.

Even though *gt_mel_* is involved in generating abdominal ablations, hybrids with no functional *gt_mel_* allele also show abdominal ablations but at lower frequency (Figure 3B, (47)). This means that other alleles in the genome are involved in producing the maladaptive trait. I introgressed a *gt_mel_^X11^* allele into the background of 200 DGRP (*Drosophila* Genetic Reference Panel) lines (Figure S13) to assess whether autosomal variants segregating within *mel*, other than those in *gt_mel_*, would affect the frequency of abdominal ablations in hybrids. Using GWAS, I found a strong association between a 75.7kb haplotype in 3L which harbors nine genes: *cindr*, *CG15544*, *tll*, *Cpr100A*, *CG15545*, *CG15546*, *CG15547*, *CG12071*, and *Sap-r* (Figure 4). Of these nine, the only gene known to interact with *gt* is *tll*. Other haplotypes associated with the frequency of abdominal ablations show a much more modest association, often found in only one sex (Table S8).

**FIGURE 4.**
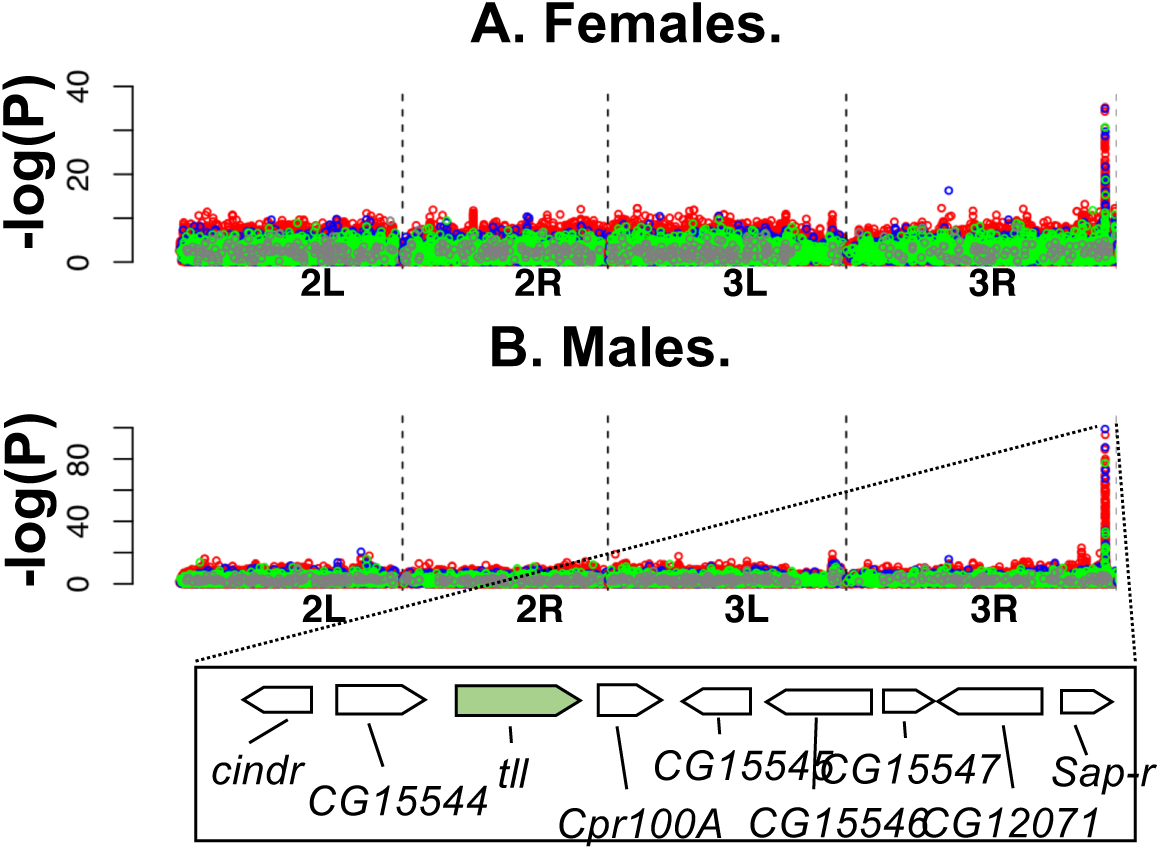
A *D. melanogaster* third-chromosome haplotype that encompasses *tll* is associated with the prevalence of abdominal ablations in *gt^X11^* mutants. Association study of autosomal genetic variants associated with the frequency of abdominal ablations in *gt^X11^/X^san^*hybrids using segregating variation within *D. melanogaster*. A haplotype of 54kb in the tip of the *X*-chromosome is strongly associated with the presence of abdominal ablations in both males (**A**) and females (**B**). Each panel shows a different chromosome arm. The haplotype 9 genes: *cindr*, *CG15544*, *tll*, *Cpr100A*, *CG15545*, *CG15546*, *CG15547*, *CG12071*, and *Sap-r.* Of these nine, *tll* is the only one known to interact with *gt*. Green: insertions, blue: deletions, red: SNPs, purple: multinucleotide polymorphisms.

To determine whether *gt_mel_* and *tll_mel_* genes interact genetically in causing HI, I generated double mutant females carrying loss of function mutations of *gt* and *tll* (*gt^X11^/FM7::GFP, tll_mel_^ΔGFP^/TM3 Ser Sb*) and crossed them to *san* males. First, I scored whether the presence of *tll_mel_* had any effect on hybrid female viability by itself. I found no effect of *tll_mel_* in hybrid female viability in an otherwise heterozygote F1 background (*FM7::GFP /X_san_, tll_mel_^ΔGFP^/3^san^*vs. *FM7::GFP /X_san_, TM3 Sb /3_san_*; Table 1). Next, I scored whether the presence of *tll_mel_* had an effect on hybrid female viability in a *gt_mel_^x11^* background. Hybrid *mel/san* females that have only a functional copy of *gt_san_* (i.e., carry a *gt_mel_^x11^* allele) and are hemizygous for *tll* (i.e., only have *tll_san_*) are more likely to survive to adulthood than *gt_mel_^x11^*-carrying females and a functional *tll_mel_* (*gt_mel_^X11^/X_san_, tll_mel_^lΔGFP^/3_san_* vs. *gt^X11^/X_san_,TM3, Sb /3_san_* Table 1A). These results suggest that while removing *tll_mel_* on its own has no major effect on HI, removing both *gt_mel_* and *tll_mel_* has a positive effect in viability that is larger than removing either allele individually.

**TABLE 1.**
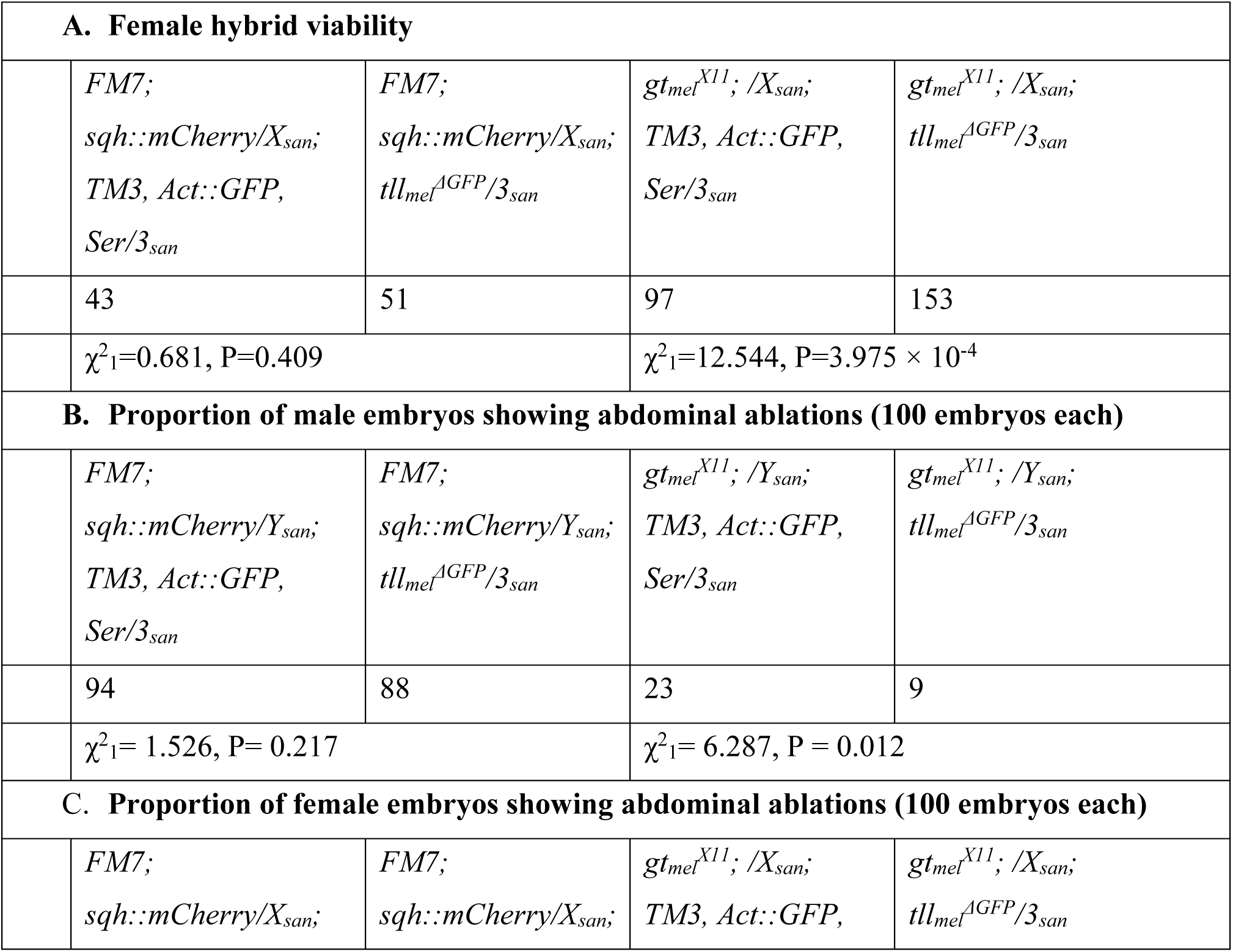
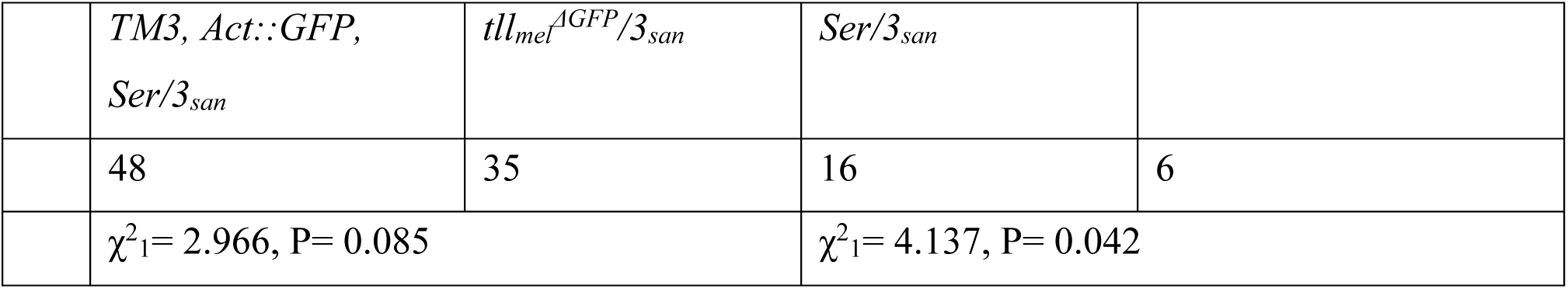
*tll_mel_* exacerbates the defects caused by *gt_mel_* in *mel/san* hybrids. **A.** Females that carry functional *gt_mel_* and *tll_mel_* alleles are more likely to die than females that carry an abrogation of *gt_mel_* (regardless of the genotype at *tll*) or that carry a functional *gt_mel_* and a null allele of *tll_mel_*. Tests with a different *tll* loss of function allele show similar results (Table S10). **B.** *mel/san* hybrid males with functional copies of *gt_mel_* and *tll_mel_* are more likely to show abdominal ablations than hybrid males with an abrogated copy of *tll_mel_*, an abrogated copy of *gt_mel_*, or abrogated copies of both. **C.** Similar to males, *mel/san* hybrid females with functional copies of *gt_mel_* and *tll_mel_* are more likely to show abdominal ablations than hybrid males with an abrogated copy of *tll_mel_*, an abrogated copy of *gt_mel_*, or abrogated copies of both. In all three cases, I used a *χ^2^* proportion test for the two pairwise comparisons.

*tll_mel_* also has a role in the frequency of abdominal ablations but only in the absence of *gt_mel_*. Abrogating *tll_mel_* in a F_1_ *gt_mel_/gt_san_* background has no detectable effect in the frequency of abdominal ablations in hybrid males or hybrid females with a functional copy of *gt_mel_*. In a *gt_mel_ ^x11^* background, abrogating *tll_mel_* decreases the proportion of male and female embryos showing abdominal ablations (Table 1B and 1C respectively). These results suggest —just as occurs with female viability— the absence of *gt_mel_* and *tll_mel_* together has a larger positive effect than the absence of each allele individually. Table S9 shows similar analyses with a different *tll* loss-of-function allele. Notably, in the conspecific crosses and the three possible interspecific crosses between *D. melanogaster* females and males from other species, *gt_mel_^X11^* and *tll_mel_^ΔGFP^* have no effects on viability (Table S10).

Finally, I tested the effect of disrupting the *tll_san_* in hybrids*. tll_san_^ΔdsRed^* did not rescue *tll_mel_^ΔGFP^* in hybrids. In *mel/san* hybrids, the *tll_san_^ΔdsRed^* deletion had no effect on female viability when tested in hybrids with multiple *mel* backgrounds (Table S11). This result suggests that removing *tll_san_* in *mel/san* hybrids has no effect on hybrid viability in an otherwise heterozygote hybrid background. Next, I tested the effect of *tll_san_^ΔdsRed^* in the null-*gt_mel_*, *gt_mel_^x11^* background. I found that the *tll_san_* deletion does not improve viability in *gt_mel_^x11^*-carrying *mel/san* females either (Table S12). These results suggests that even in the absence of a functional *gt_mel_* allele, removing *tll_san_* has no effect on hybrid female viability. Since the reciprocal deletion (removing *tll_mel_* and exposing the *tll_san_* allele) does improve female hybrid viability in *gt_mel_^x11^*-carriers, these results indicate that the presence of *tll_mel_*, but not of *tll_san_*, is involved in the HI of *mel/san* female hybrids.

##### Molecular Evolution

Gap genes *gt* and *tll* have phylogenetically conserved roles in pattern formation, as evidenced by their functionally conserved outputs in blastoderm embryos of distantly related *Drosophila* species (76) and beyond (77, 78). Yet, they have diverged sufficiently between *mel* and *san* such that they malfunction in hybrids. I therefore conducted analyses to assess the patterns and mechanism of divergence of the *gt* coding sequence in the *melanogaster* species subgroup **and across the *Drosophila* genus**.

Both *gt* and *tll* are highly conserved in their coding regions. The bZip domain that confers the Gt protein its ability to bind DNA is highly conserved across animals (79) and shows no fixed differences among *Drosophila* species (Figure S14). Gt shows only thirteen single amino acid substitutions in the *melanogaster* species subgroup (Figures S15 and S16), six of which occur on the branches connecting *mel* and *san*. *Giant* also contains three low complexity regions (including polyQ) that show extensive variation both within and between species (Figure S16). *tll* is also conserved in the *melanogaster* species subgroup; only four residues differ between the *tll* alleles in the *yakuba* clade and the *melanogaster* clade: Val509Asp, Arg1118Lys, Ser1208Thr, Leu1246Met (Figure S17). Only Val509Asp represents a change in the type of amino acid (a change from a nonpolar to an acidic residue).

I also investigated whether substitutions in these two genes might have been driven by adaptive evolution. Both genes, *gt* and *tll*, fall within the slowest-evolving quantile of genes comparing *mel* with *yak* or *san* (25% and 10% respectively) and, noting the limited power to detect selection as a result, show no signature of accelerated evolution (Table S13). At least one DMI partner(s) of *gt^mel^* is (are) unique to the *D. santomea* lineage The phylogenetic occurrence of developmental defects provides an additional hypothesis to test: I next evaluated whether the unknown genetic element(s) in the *D. santomea* genome that must interact with *gt_mel_* and *tll_mel_* (*gt_sim_*) to cause HI are also present in *D. teissieri* (a close relative of *D. santomea*, Figure 1A). The crosses ♀*mel* × ♂*tei*, ♀*sim* × ♂*tei* and ♀*mel* × ♂*tei* all produce viable adult females and males that die as late larvae/early pupae. Little embryonic lethality is observed in any of these two crosses and the rare embryos that die do not show abdominal ablations (Figure 1E, Figure S1). All *D. teissieri* lines showed similar levels of viability in each cross (Figure S18). Neither the presence or the absence of *gt_mel_* (Figure S5 and Figure S11 respectively) have any effect on the viability of *mel/tei* hybrids suggesting that the *tei* background is missing at least one allele that interacts with *gt_mel_* to cause HI. These results suggest that at least one of the partners of this incompatibility—one that remains unidentified— is specific to *san*, and evolved after *san* and *tei* diverged (Figure S19).

## DISCUSSION

Hybrid inviability is a strong barrier to gene exchange between species. While it is clear that this trait is often caused by epistatic interactions between alleles from different species, few examples have been identified to the gene level. Here, I identified two genes, *gt* and *tll,* which contribute to HI in hybrids between two *Drosophila* species. The two genes belong to the gap gene network, a highly conserved pathway that is in charge of establishing embryonic polarity in insects (76–78, 80–83). The *mel* alleles of these two genes are necessary and sufficient to cause a male abdominal ablation phenotype that is particularly common in hybrid males of the cross. I also find support for a third (or even more) elements that are exclusive to *D. santomea* and remain unidentified. Additional members of the gap gene network must have functionally diverged between the two species and contribute to HI. These are not the only alleles that contribute to inviability in the cross but are sufficient to cause the abdominal ablation defect that is particularly common in hybrid males of the cross (38, 47).

The involvement of *gt_mel_* and *tll_mel_* in HI indicates that one or more features of their function have diverged between relatively closely related species, despite their broad conservation across the Diptera ((76, 80, 81, 83); but see (82, 84, 85)), bees (78) and beetles (77, 86). The results shown here confirm speculation that HI can arise in phylogenetically conserved gene networks regulating development (30, 39, 87). The involvement of *gt_mel_* and *tll_mel_* in HI suggests that their function has diverged across *Drosophila* species. Natural selection has driven the evolution of regulatory elements of many developmental genes in *Drosophila* which has led to a rapid turnover (28, 76, 88, 89). Yet, neither *gt* nor *tll* show signatures of positive selection in their coding sequences. These results also suggest that the evolution of the different components involved in the DMI occurred at different times and is unlikely to have had any role on speciation. The deleterious effects caused by *gt_mel_* seem to be common to *D. melanogaster* and the species in the *D. simulans* clade, suggesting that the allele necessary for the incompatibility evolved before these species split between three and five million years ago (45, 46). Because the presence of *gt^mel^* has no quantifiable viability effect in *mel/tei* hybrids, at least one of the genetic factor(s) that interact with *gt^mel^* to cause abdominal ablation in hybrid embryos must have arisen after *D. santomea* and *D. teissieri* split between 1 and 2.5 million years ago (90, 91). An alternative divergence scenario is that at least one of the genetic components of the DMI evolved in the *tei* branch to suppress HI. Regardless of which of these two scenarios is correct, the components of the DMI must have evolved at different times in the two lineages, and the interactions with *giant* that cause abdominal ablation could not have been involved with any speciation event in the *melanogaster* species subgroup (Figure S11). Instead, these loci must have evolved independently in each lineage, accumulating differences as the genomes diverged after speciation, a scenario in accord with the Dobzhansky Muller model (1, 6, 7, 44, 92, 93). Mapping the allele(s) that interact with *gt^mel^*and *tll^mel^* in the *D. santomea* genome is the next step in describing how genomic divergence creates hybrid defects.

Previous comparative analyses of gap gene expression in dipterans indicates gene network evolution in spite of a conserved developmental phenotype (83), which suggests continual fine-tuning of the genetic interactions in the gap gene network within species. Coevolved compensatory changes have been proposed to cause HI in instances in which the phenotypic output of a gene network is under moderate stabilizing selection (94–97). Molecular functional evolution without phenotypic change, or developmental systems drift, has been hypothesized to underlie hybrid breakdown involving canalized traits such as embryogenesis and gametogenesis (30). The HI involving *gt_mel_* and *tll_mel_*may exemplify compensatory changes resulting in a stable phenotype when comparing pure species, but in an aberrant phenotype in hybrids.

The identification of two of the partners of a hybrid incompatibility in *mel/san* hybrids is just a first step on discovering why these hybrids die. The results shown here indicate that other genetic partners are involved. First, the *D. santomea* partner remains to be identified. Second, *gt_mel_* has differences levels of penetrance in crosses involving different *D. santomea* fathers, which in turns indicates the existence of genetic modifiers segregating in *D. santomea*. Besides the obvious step of idnetyfying these genetic partners, future work should focus on the evolution of gene expression in the gap gene network within the *melanogaster* group and examine the extent to which regulatory sequences are involved in hybrid incompatibilities. The importance of compensatory evolution in regulatory sequences remains a largely unexplored aspect of HI and more broadly, in the evolution of species boundaries.

The introduction of a developmental genetics perspective to speciation studies has the potential to shed new light on the study of hybrid inviability (98). Hybrid inviability is a natural experiment to test genetic interactions between diverging genomes: the molecular interactions that go awry in hybrids reveal evolutionary divergence of the genes involved, or the timing, location, or amount of their expression (97). The interactions between *gt_mel_*, *tll_mel_* and the unknown factors in the genome of *D. santomea*, had nothing to do with setting the speciation process in motion in the *melanogaster* species subgroup. They are also not involved in currently keeping species apart as *D. melanogaster* and *D. santomea* do not naturally hybridize. The results shown here should be viewed in the broad context of genome divergence and how genomes keep evolving long after speciation has occurred. This represents a path forward in terms of how to think about stability vs. change of different functional units within the genome and different developmental processes. The identification of *giant* and *tll* as genes involved in HI is the first indication that genes involved in early embryonic development, a canonical example of a conserved developmental process, functionally co-evolve at a pace sufficient to cause HI.

## METHODS AND MATERIALS

### 1. STOCKS

#### 1.1. Drosophila melanogaster mutants

With two exceptions (see 1.1.2. *tll* alleles and 1.1.5. *gt* transgenics), all *Drosophila melanogaster* mutant stocks used in this study were originally obtained from the Bloomington Stock Center (BSC) and are listed in Table S1.

1.1.1. gt *alleles*. I obtained multiple *gt* mutant alleles from the Bloomington Stock Center (BSC) but focused on a null allele: *gt_mel_^X11^* (*y^1^ sc^1^ gt^X11^/FM6*; BSC stock #1529). The *gt_mel_^X11^* has a frameshift mutation that abrogates the bZIP DNA-binding domain (99). I confirmed that this stock did not complement a deficiency of *giant* (*Df(1)Exel6231*; BDSC stock #7706) and verified *gt^X11^* was indeed a loss-of function allele. To differentiate between embryos carrying the *gt* allele or the balancer, I rebalanced each stock with a *FM7 Actin::GFP* to allow for genotypic distinctions in all further crosses.
1.1.2. tll *alleles*. I used two *tll* alleles. The first one was a loss-of-function allele of *tll*: *tll^1^* (100) which was procured from the Bloomington Stock center (*st^1^ e^1^ tll^1^/TM3, Sb^1^*; stock 2729). I also obtained a CRISPR-mediated *tll GFP* disruption: *tll ^Δ^*^GFP^. Construct design, construction were performed by Rainbow Transgenics (Camarillo, CA). The design for the disruption is shown in Figure S20. The construct was then injected into a *y w^1118^ D. melanogaster* stock. One hundred and five larvae were injected and one showed a GFP integration. Heterozygote transformants were identified by the GFP marker. These heterozygote flies (*tll ^Δ^*^GFP^/*tll^+^*) were then crossed to a *TM3, Sb*/*TM6B* stock to obtain *tll ^Δ^*^GFP^ */TM3, Sb* flies. The resulting mutant failed to complement the *tll^1^*mutant.
1.1.3. *Attached-X stocks*. I used two different attached-*X* stocks: *C(1)RM* and *C(1)DX*. These two stocks differ in the way that the two *X*-chromosomes are attached to each other. *C(1)RM*, or reverse metacentric, is a fusion of the *X*-chromosomes joint at their heterochromatic regions (101). *C(1)DX* is a fusion of *X*-chromosomes with a reverse acrocentric fusion (101), and the two two *X*-chromosomes at fused at the centromere in opposite directions.
1.1.4. X-Y *translocations.* (*Dp(1;Y)*) duplications and deficiencies are described below in sections 2.2 and 2.3 respectively.

#### 1.2. Other species stocks

Initial surveys of HI alleles used outbred lines (i.e., lines derived from combined individuals from several isofemale lines) for interspecific crosses (e.g.,(47, 102)). In this report, I used isofemale lines collected in nature. This allowed me to survey whether HI was a line-specific or a species-wide phenomenon.

1.2.1 *Isofemale lines*. For interspecific crosses involving *D. melanogaster gt_mel_^x11^/FM7::GFP* and *D. melanogaster df(BSC)/FM7::GFP* I used 25 *D. santomea* isofemale lines collected in the island of São Tomé. I also quantified hybrid viability for hybrids from hybrids between females from these two *D. melanogaster* mutant crosses with 25 *D. mauritiana*, 25 *D. teissieri*, and 25 *D. simulans* lines. All lines, including their collection details, are listed in Table S2.
1.2.3. *Other mutant stocks*. Finally, I used five mutant stocks from three other species from *D. melanogaster*. Attached *X* stocks from *D. simulans* (*y w*, (103) reviewed in (104)) were donated by D. Presgraves. Attached *X* from *D. yakuba* (*y*) were produced by J. Coyne (105). Attached *X* from *D. santomea* (*gn*) were donated by A. Llopart (49, 105, 106). *yellow* stocks from *D. simulans, and D. mauritiana* were obtained from the San Diego stock Center (now Cornell Stock Center).

### 2. INTERSPECIFIC GENETIC CROSSES

#### 2.1. Wild-type crosses

Pure species males and females of each species were collected as virgins within 8 hours of eclosion under CO_2_ anesthesia and kept for three days in single-sex groups of 20 flies in 30mL, corn meal food-containing vials. Flies were kept at 24°C under a 12 h light/dark cycle. On day four, I assessed whether there were larvae in the media. If the inspection revealed any progeny, the vial was discarded.

On the morning of day four after collection, I placed forty males and twenty females together at room temperature (21°–23°C) to mate *en masse* on corn meal media. To produce hybrid adults, vials were inspected every five days to assess the presence of larvae and/or dead embryos. In order to maximize the lifespan of the parental, I kept all the vials laying on their side. I transferred all the pure species adults to a new vial (without anesthesia) every five days. This procedure was repeated until the cross stopped producing progeny (i.e., females were dead). Once L_2_ larvae were observed in a vial, I added a solution of 0.05% propionic acid and a KimWipe (Kimberly Clark, Kimwipes Delicate Task, Roswell, GA) to the vial. I counted the hybrids as they hatched by anesthetizing them with CO_2_.

To quantify embryonic lethality, I mixed males and females as described above in a 30mL plastic vial with corn-meal or molasses food. After two days, I transferred the adults from the vials that showed larvae to an oviposition cage with apple juice media and yeast. The plates were inspected every 48 hours for the presence of viable eggs. To quantify embryonic lethality, I counted the number of egg cases (viable embryos) and the number of brown eggs (dead embryos) in each oviposition cage. Rates of embryonic lethality were calculated as the proportion of brown eggs over the total number of eggs. For all interspecific crosses within the *yakuba* species complex (*D. santomea*, *D. yakuba*, and *D. teissieri*), I only used lines that have been shown to be infected by *Wolbachia* (wB) to minimize any possible effect of endosymbionts on hybrid inviability (51).

#### 2.2. Genome wide association studies (GWAS)

##### 2.2.1. Hybrid inviability GWAS

I identified polymorphisms segregating in *D. melanogaster* associated with hybrid inviability and penetrance of abdominal ablations in *mel/san* F1 hybrids. I leveraged the *Drosophila* Genetic Reference Panel (DGRP) genotype information (54, 55) to identify variants within *D. melanogaster* associated with the presence of abdominal ablations when crossed to *D. santomea* males. In total, I used 200 *D. melanogaster* lines, all of which are listed in Table S1. In order to identify SNPs associated with abdominal ablations in hybrids, I mated females from each of the *D. melanogaster* lines with *D. santomea* males following the procedure described in 2.1 (immediately above). The response phenotype was the percentage of larvae that showed abdominal ablations (scored as described above).

I submitted the percentage of ablated embryos of the 200-line study, not sexed, to the web portal dgrp2.gnets.ncsu.edu for analysis. Since I could not differentiate between female and male embryos (DGRP lines do not carry a *y*^-^ marker), I collected a single ablation score per line, combining the two sexes. Associations between the phenotype (i.e. percentage of ablated embryos) and genome wide polymorphisms within *D. melanogaster* were calculated by the DGRP algorithm, using a linear mixed model, which accounts for any effects of *Wolbachia* infection, common polymorphic inversions, and cryptic relatedness in the DGRP lines, as described in detail in (54). This GWAS incorporates information from 1.9 million SNP variants. The genome-wide significant threshold at the 5% significance level was determined after a Bonferroni correction for multiple testing (107) and adjusting the critical P-value for significance to as 2.60 × 10^-8^ (0.05/1,900,000).

##### 2.2.2. GWAS restricted to the autosomes

Next, I assessed if any of the autosomal segregating variants in the DGRP modified the effects of *gt_mel_^X11^*. To this end, I introgressed an *FM7::GFP* balancer and the null allele of giant, *gt_mel_^X11^* into 200 lines from the DGRP. The introgression protocol is shown in Figure S13 and involved introgressing *FM7::GFP* (10 generations of backcrossing) by following the *Bar* and *GFP* markers of the balancer (Figure S13A). On generation 11, and for each DGRP stock, *gt_mel_^X11^/FM7* females, were crossed to a male from each of the DGRP stocks that carried a *FM7:GFP* (from the first round of introgression). F1 females with yellow mouth parts carried *gt_mel_^X11^* and *FM7::GFP* and were again crossed to DGRP males carrying *FM7:GFP*. This procedure was repeated for 10 generations (Figure S13B). The result from this crosses was having *FM7::GFP*/*y_mel_^-^ gt_mel_^X11^* in 200 different genetic backgrounds.

Next, I assessed the effects of the different DGRP genetic backgrounds on the penetrance of *gt_mel_^X11^* in *mel/san* hybrids. I crossed females from each of these 200 stocks to *D. santomea* CAR1600 males. I separated the progeny into four different categories. Females have black mouthparts (i.e., *FM7::GFP/X_san_* and *y_mel_^-^ gt_mel_^X11^/X_san_*; *y_san_* rescue *y_mel_^-^*); males (either *FM7::GFP* /*Y_san_* or *y_mel_^-^ gt_mel_^X11^*/ *Y_san_*) have brown mouthparts. Balancer carriers (i.e., *FM7::GFP/X_san_* and *FM7::GFP*/ *Y_san_)* have a *GFP* marker. The goal of this experiment was to find autosomal factors that would affect the frequency of abdominal ablations in a *gt_mel_^X11^* background in both females and males; I scored the percentage of ablated embryos in *y_mel_^-^ gt_mel_^X11^*/ *Y_san_* males and *y_mel_^-^ gt_mel_^X11^*/ *X_san_* hybrid females apart. Because of the experimental design (introgression of a full *y_mel_^-^ gt_mel_^X11^X*-chromosome on multiple autosomal backgrounds), I did not study possible *X*-linked modifiers of the penetrance of *gt_mel_^X11^*. Association mapping was done as described immediately above but splitting the two sexes and excluding markers from the *X*-chromosome (section 2.2.1. Hybrid inviability GWAS).

#### 2.3. Duplication mapping

Duplication mapping identifies dominant (or semidominant) alleles in the *D. melanogaster X-*chromosome (*X^mel^)* that cause inviability in hybrids resulting from interspecific crosses. The technique uses stocks provided by Bloomington Stock Center in which segments of the *X-*chromosome have been duplicated and attached to the *Y-*chromosome by BAC recombineering (57). I used two classical *Drosophila* techniques—attached-*X* females (described in section 1.1.2), and *X-Y* chromosome fusions—to finely characterize the identity of HI alleles in the *X_mel_*. *Drosophila melanogaster* attached-*X* females carry two *X-*chromosome fused together which carry recessive alleles for easy identification. I used two genotypes of attached-*X* females: *C(1)DX* and *C(1)RM*. Unless otherwise noted, all crosses used *C(1)DX*. These females can carry a *Y_mel_* chromosome and remain morphologically female. When these females carry both an attached-*X* and a Y chromosome, they produce attached-*X* gametes and *Y_mel_* gametes.

Previous experiments have shown that when *C(1)DX* (or *C(1)RM*) females are crossed with *D. santomea* males the only viable genotype are F_1_ hybrid males with genotype *X_san_ Y_mel_* (Figure S1, (47)). *Drosophila melanogaster C(1)DX* females can also be crossed to *D. simulans* (108)*, D. mauritiana* (109), and *D. teissieri*. In the first two cases, the cross produces viable hybrid males with an *X*-chromosome from the father, and a *Y_mel_*. In the cross with *D. teissieri* males, *C(1)DX* females produce viable larvae from both sexes that die before molting into pupae.

I addressed whether the introduction of small pieces of *X_mel_* in *mel/san* hybrid males cause HI. I used a panel of small *X_mel_*-chromosome fragments attached to the *Y*-chromosome [*Dp(1;Y)*] that tilled the cytological bands 2, 3 and 4 in the *X*-chromosome (12 duplications). These segments also carry two phenotypic markers: *y_mel_^+^*, and *Bar*.

Since the procedure to produce interspecific hybrid males for the four species (*D. santomea*, *D. teissieri*, *D. mauritiana*, and *D. simulans*) is identical, I only describe the protocol for only one of them, *D. santomea*. I crossed attached*-X^mel^*females [*C(1)DX*] to *D. melanogaster Dp(1;Y)* males that carry small fragments of *X_mel_* on their *Y-*chromosome. The female progeny [*mel C(1)DX/Dp(1;Y)*] will carry both the attached-*X* and the *Y-*chromosome with the small fragment of *X* to be tested. These virgin females are then crossed to *D. santomea* males to produce F_1_ hybrid males harboring an *X_san_* and a (*Y_mel_, Dp(1;Y)*) chromosomes. The crossing scheme to identify these dominant alleles on the *X_mel_* chromosome is shown in Figure S1. The effect of the *X_mel_* fragment was assessed by counting how many individuals survive the transition between three developmental stages (embryo, larvae, pupae). This approach (shown in Figure S1) has been used to identify alleles from *X_mel_* that cause hybrid inviability (35, 47, 109).

A parallel set of *X*-duplications, was not attached to the *Y*-chromosome but the third chromosome instead (*Dp(1;3)*; (57)). I introgressed these duplications into a *C(1)DX, TM3*/+ background by repeated backcrossing (4 generations) to produce *C(1)DX*; *Dp(1;3)*/ *Dp(1;3)* females. The effect of the *Dp(1;3)* was measured in the same manner as described above for *Dp(1;Y)* by measuring transition rates across developmental stages. Interspecific crosses using *C(1)DX, TM3* or *C(1)DX, TM6B* stocks yielded very low numbers of progeny (110), which precluded the collection of embryos.

#### 2.4. Deficiency mapping

Next, I studied whether the absence of *X_mel_* chromosomal segments improved or decrease hybrid inviability. Traditionally, deficiency mapping has focused on the later and has been used to find recessive alleles in the genome of the other species that are lethal when the *D. melanogaster* allele is deleted (10, 44, 61). I took a different approach and focused on dominant alleles: those that when removed increased hybrid viability. I used *mel* females from stocks containing known genomic deletions, or “deficiencies” (*df*, Bloomington *Drosophila* Fly Stock Center) maintained as heterozygotes against a balancer (*Bal*) chromosome carrying a dominant homozygous lethal mutation, to *D. santomea* (*san*) males. Seven deficiencies encompass *gt_mel_* (listed in Table S1). Virgin *D. melanogaster* females were crossed to *D. santomea* males following previously described procedures (44, 111). (Behavioral isolation seems to be complete in the reciprocal direction (38).) Crosses were kept until no more progeny was produced out of each vial, usually 45 days after they were set up. The effect of each hemizygous region on the viability of hybrid female offspring was measured by comparing the ratio of *df/san* to *Bal/san* hybrid females. The significance of the departure was assessed by a χ ^2^ test followed by a Sidak’s multiple comparison correction. P-values were considered significant if to P < 0.007 (( = 0.05 adjusted for 7 multiple comparisons). If the deletion has no effect on hybrid viability, the ratio of F_1_ *df/san* to the total number of progeny (F_1_ *Bal/san* + F_1_ *df/san*) will not differ from 0.5. If the deletion reveals alleles in the *san* genome that cause complete inviability, the ratio will be equal to 0 (only progeny carrying the *Balancer* will survive). If the *D. melanogaster* deficiency uncovers a recessive region of the *D. santomea* genome that compromises hybrid fitness but does not cause complete lethality, the ratio (F_1_ *df/san* /Total) will be greater than 0 and significantly lower than 0.5. Finally, and the target of this study, if the ratio (F_1_ *df/san* /Total) is significantly higher than 0.5, then the *df* is removing a dominant (or semidominant) contributor to HI. This last category must be seen with caution as Balancer chromosomes carry deleterious alleles that might bias the ratio upwards. To minimize the potentially deleterious effect of any given balancer, I used seven different *X*-chromosome balancers to replicate crosses involving *gt^X11^*and *df(gt)*: *FM6::GFP*, *Binscy*, *Basc*, Bascy, *Binsn, Binsncy, FM4* and *FM7a*.

### 3. INTRASPECIFIC CROSSES: DOSAGE EFFECTS

I tested whether any of the mutants caused phenotypic defects or inviability by dosage effects. This is important because *Dp(1;Y)* carrying males have two copies of the genes under study while wild-type males only have one copy. Similarly, *df*-carrying females only have a single copy of a gene (i.e., they are hemizygous) while wild-type females have two copies of that gene.

I studied whether any of the used duplications cause inviability in males for being diploid (when they are normally hemizygous). All crosses were done as described in section 2. 2. (Duplication mapping) but instead of using heterospecific males, I used males from 25 different *D. melanogaster* isofemale lines. The list of isofemale lines used for these experiments is shown in Table S2. If there is a dosage effect (i.e., carrying *Dp(1;Y)* and thus two copies of a gene while being male is deleterious), one might expect inviability and/or developmental defects in these crosses.

I did a similar analysis to assess for potential haploinsufficiency in *df*-carrying females. I tested whether any of the used deficiencies cause inviability in females for being hemizygous (when they are normally diploid) in the same twenty-five *D. melanogaster* backgrounds described immediately above. I measured the ratios of *df*- and *Balancer*-carrying females using the methods described above.

### 4. CUTICLE PREPARATION

I generated cuticles for wild-type (i.e., progeny produced by crossing wild-type stocks), *Dp(1;Y)*-carrying and *gt^x11^*-carrying hybrids. The details to produce the three types of cuticles are described as follows.

#### 4.1 Wild-type hybrids

To collect sex-specific hybrid cuticles, I used *D. melanogaster* a *y^1^ w* stock. *Drosophila melanogaster y^1^ w* females were crossed to *D. santomea* males. Inseminated females were allowed to deposit on apple juice plates overnight and embryos were aged for 24 hours before scoring. To prepare cuticles, I used a slightly modified the protocol described in (38, 47). Briefly, I dechorionated embryos using double-sided scotch tape. To devitellinize brown (dead) embryos. I made a cut on the vitelline membrane and removed the rest of the cuticle with a tungsten dissection needle and placed them in a 3:1 solution of acetic acid and glycerol for 48 hours. After this period, cuticles were mounted on a pre-clean glass slide (VMR VistaVision^TM^, VWR; cat. no. 16004-422; Radnor, PA) on 20μl of a 1:1 solution of Hoyer’s media (kindly donated by Dr. Daniel McKay) and acetic acid. Embryos without pigmentation of the mouth hooks were scored as male with the genotype *y^1^ w* / *Y_san_*. Embryos with black mouth parts were identified as females (*y^1^/ y_san_*) as the *y^1^* allele is complemented by the homologous *y_san_*. Embryos were visualized and imaged with an Olympus BX61 dark-field microscope at the Microscopy facility of the Pathology department at UNC.

#### 4.2. *Dp(1;Y)* carrying male hybrids

To collect *X_san_ Y_mel_* (*Dp1;Y*), I followed a similar procedure to that described immediately above (section 4.1). Individuals with brown mouth parts were concluded to be females carrying *C(1)DX* (*y^-^/y^-^* homozygotes). Individuals with black mouth parts had two possible genotypes: metafemales carrying three *X* chromosomes (*C(1)DX y^-^* and *X_san_; y^-^/y^-^/y_san_*) or *X_san_/Y_mel_, Dp(1,Y)* males (*y_san_*). Since metafemales (i.e., *C(1)DX y^-^*/*X_san_* females) are thought to show a low rate of embryonic defects (Matute and Gavin-Smyth 2014b), the pooling of these two categories underestimates the penetrance of the alleles responsible for the ablation. This bias is not a concern as it should occur at a similar rate in all crosses involving *C(1)DX*. All rates of penetrance using *C(1)DX, Dp1;Y* females should then be considered an underestimation.

#### 4.3 *gt_mel_^x11^* carrying female hybrids

I used a similar approach to collect cuticles for hybrids carrying a *gt*-null allele (*gt_mel_^x11^*). To score hybrid defects on *gt*^-^-carriers, I used a *y^1^ sc^1^ gt^X11^* genotype. This stock was purchased as *y^1^ sc^1^ gt^X11^*/*FM6* (Table S1, Row 1). I rebalanced the stock over a *y^1^ FM7* chromosome carrying an *Gal4*-Actin::*UAS-GFP* reporter. To produce *gt^x11^*/*san* cuticles, *gt^x11^*/*FM7Actin::GFP D. melanogaster* females were crossed to *D. santomea* males. *GFP* minus embryos that failed to hatch were separated and prepared for cuticle mounting using standard procedures (see (Gavin-Smyth and Matute 2013)). Both, *GFP* minus embryos (*gt^x11^*-carriers) and *GFP* plus embryos (*FM7* carriers) were separated by the color of their mouth-hooks as described above to identify by sex. Cuticles of other interspecific crosses (e.g., *mel/tei, mel/mau, mau/sim*) were collected, prepared, and imaged using the same scheme.

### 5. GENOME SEQUENCING

Next, I studied the patterns of polymorphism in *gt* across the nine species of the *melanogaster* species subgroup. This involved (*i*) collecting flies in nature, (*ii*) sequencing their genomes, and (*iii*) aligning them. These three steps are described in detail as follows.

#### 5.1. Stock collection

I collected lines from five species in the *melanogaster* group in their natural habitat. *Drosophila santomea* and *D. yakuba* were collected in the volcanic island of São Tomé. *Drosophila teissieri* was collected in the highlands of the island of Bioko, Equatorial Guinea. *Drosophila simulans* was collected in Bioko, São Tomé and Zambia. In all cases, single females were collected with banana traps, anesthetized with FlyNap (triethylamine, Carolina Biological Supply Co.) for 2-5 minutes. Individual females were then placed in plastic vials with instant potato fly media (Carolina Biological Supply Co.) and were allowed to oviposit until their death. Vials with progeny were hand carried to the USA (USDA permit: P526-150127-009) and progeny was transferred to a corn-meal diet in 100 mL vials. *Drosophila mauritiana* stocks were kindly donated by D. Presgraves.

#### 5.2. Genomic data

All the genomes of the lines used in this study were previously published. I downloaded available raw reads (FASTQ files) for *D. yakuba* (50, 91, 112, 113), *D. santomea* (50, 91), *D. teissieri* (50, 91), *D. mauritiana* (114, 115), *D. sechellia* (50, 116), *D. simulans* (114, 117), and *D. melanogaster* (118, 119) from NCBI and mapped them to the corresponding reference genome (see below). All the accession numbers are listed in Table S14.

#### 5.3. Read mapping and variant calling

Reads were mapped to the closest reference genome using bwa version 0.7.12 (120, 121). Reads from *D. yakuba*, *D. teissieri*, and *D. santomea* were mapped to the *D. yakuba* genome version 1.04 (122), and reads from *D. simulans, D. sechellia*, and *D. mauritiana* were mapped to the *D. simulans w^501^* genome (123). I used Samtools version 0.1.19 (124) to merge Bam files. I used GATK version 3.2-2 RealignerTargetCreator and IndelRealigner functions to identify indels and polymorphic sites (125, 126). Read mapping and SNP genotyping were done independently for the *D. yakuba* and *D. simulans* clades using GATK UnifiedGenotyper but in both cases I used similar parameters and files. The parameter het was set to 0.01. I also used the following filters to all resulting vcf files: QD = 2.0, FS_filter = 60.0, MQ_filter = 30.0, MQ_Rank_Sum_filter = -12.5, and Read_Pos_Rank_Sum_filter = -8.0. If a site had a coverage below five reads or above than the 99^th^ quantile of the genomic coverage distribution for the given line, that site was assigned an ‘N’.

#### 5.4. Sequencing, Indel identification

I studied the positions of indels in the *giant* locus across the *melanogaster* species subgroup. To genotype indels, I used GATK UnifiedGenotyper with the -glm INDEL flag for just the sequence orthologous to the *D. melanogaster giant* gene. I then generated fasta files for the *giant* locus. No coverage thresholds were used for indel genotypes.

#### 5.5. Genomic Alignments

I generated genome alignments that included *D. melanogaster*, *D. simulans* and *D. yakuba*. The *D. yakuba* and *D. simulans* reference genomes were separately aligned to the *D. melanogaster* genome using nucmer version 3.23 with parameters –r and –q. Next, I used the dmel6.01 annotation: ftp.flybase.net/genomes/Drosophila_melanogaster/dmel_r6.01_FB2014_04/gff/ dmel-all-r6.01.gff.gz (127) to identify the *giant* coding region (*D. melanogaster X* chromosome: 2,427,113 – 2,429,467). These alignments were the used to extract polymorphism data for this region for 8 species in the *melanogaster* species subgroup. In total, I included data for 903 sequences. The subsequent alignment was visually inspected using Mesquite version 3.04 (128) to ensure indels were aligned and did not disrupt codons.

### 6. DETECTION OF NATURAL SELECTION

#### 6.1. PAML

Next, I studied whether the *giant* locus had evolved though natural selection. The first approach to detect positive selection was to count the number of synonymous (dS) and non-synonymous (dN) substitutions in each branch and calculate the ratio between these two variables. First, I generated a consensus sequence for each of the species in the *melanogaster* species subgroup. Next, I ran PAML version 4.8 (129, 130) to calculate dN/dS ratios. I used four sets of parameters: basic model (model=0), free ratios (model=1), 3 ratios (model=2, tree = ((mel, (sim, sech, mau) $2)$1, ((yak, san), tei) $3);), and 2 ratios (model=2, tree = ((mel, (sim, sech, mau))$1, ((yak, san), tei) $2)). A dN/dS ratio significantly higher than 1 means positive selection, while a dN/dS ratio significantly lower than 1 means negative/purifying selection ((129, 130) but see (131)). dN/dS values not significantly different from zero represent neutral evolution.

## Supplementary Information

### SUPPLEMENTARY TABLES

**TABLE S1.**
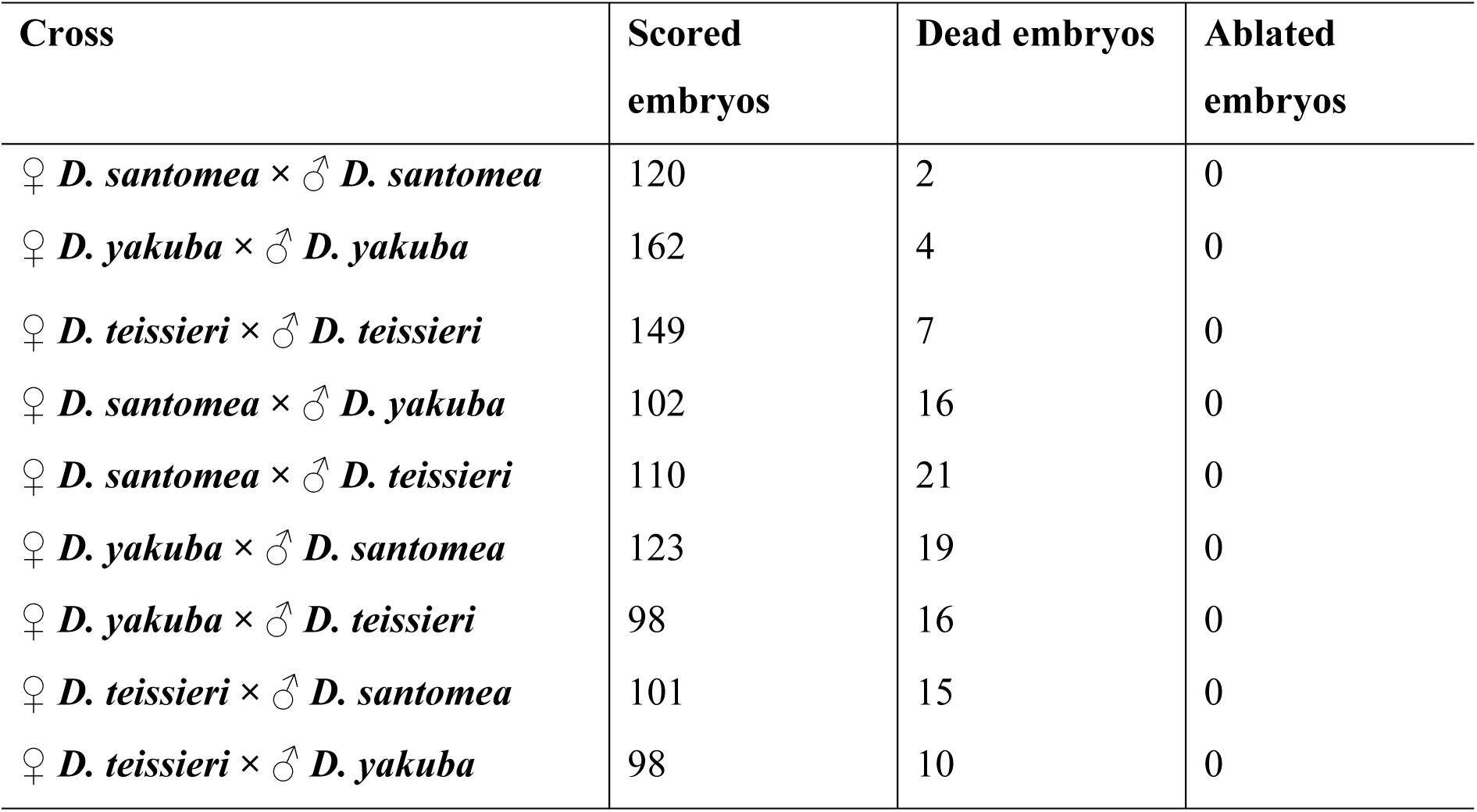
Rates of ablation in F1 hybrids between the species of the *yakuba* species complex. None of the nine possible F1 hybrids show abdominal ablations.

**TABLE S2.**
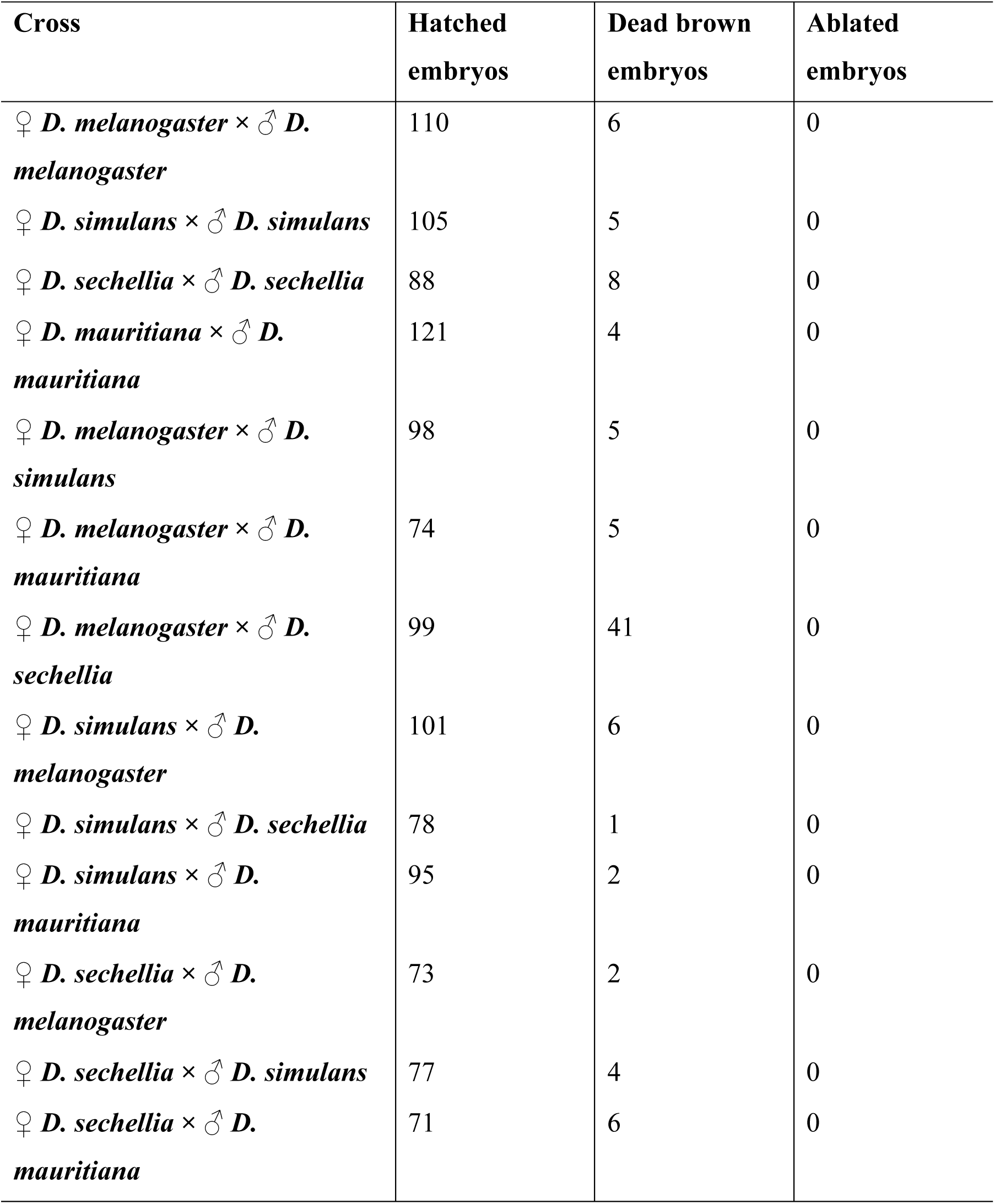
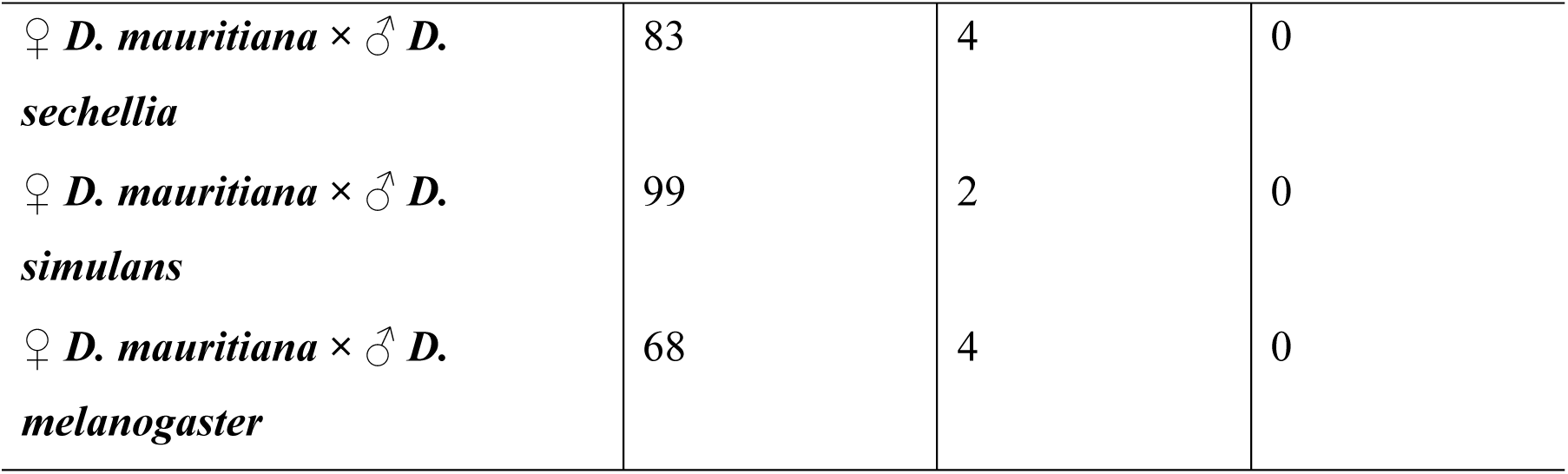
Hybrids between *D. melanogaster* females and *D. simulans* and *D. mauritiana* show no abdominal ablations or embryonic lethality. The reciprocal crosses (♀*D. simulans* × ♂ *D. melanogaster;* ♀*D. mauritiana* × ♂*D. melanogaster*) show high levels of female embryonic lethality but eggs do not turn brown (i.e., the assay of brown embryos cannot detect early embryo lethality).

| Cross | Hatched embryos | Dead brown embryos | Ablated embryos |
| --- | --- | --- | --- |
| $\text{♀} D. melanogaster \times \text{♂} D. melanogaster$ | 110 | 6 | 0 |
| $\text{♀} D. simulans \times \text{♂} D. simulans$ | 105 | 5 | 0 |
| $\text{♀} D. sechellia \times \text{♂} D. sechellia$ | 88 | 8 | 0 |
| $\text{♀} D. mauritiana \times \text{♂} D. mauritiana$ | 121 | 4 | 0 |
| $\text{♀} D. melanogaster \times \text{♂} D. simulans$ | 98 | 5 | 0 |
| $\text{♀} D. melanogaster \times \text{♂} D. mauritiana$ | 74 | 5 | 0 |
| $\text{♀} D. melanogaster \times \text{♂} D. sechellia$ | 99 | 41 | 0 |
| $\text{♀} D. simulans \times \text{♂} D. melanogaster$ | 101 | 6 | 0 |
| $\text{♀} D. simulans \times \text{♂} D. sechellia$ | 78 | 1 | 0 |
| $\text{♀} D. simulans \times \text{♂} D. mauritiana$ | 95 | 2 | 0 |
| $\text{♀} D. sechellia \times \text{♂} D. melanogaster$ | 73 | 2 | 0 |
| $\text{♀} D. sechellia \times \text{♂} D. simulans$ | 77 | 4 | 0 |
| $\text{♀} D. sechellia \times \text{♂} D. mauritiana$ | 71 | 6 | 0 |
| ♀ *D. mauritiana* × ♂ *D. sechellia* | 83 | 4 | 0 |
| ♀ *D. mauritiana* × ♂ *D. simulans* | 99 | 2 | 0 |
| ♀ *D. mauritiana* × ♂ *D. melanogaster* | 68 | 4 | 0 |

**TABLE S3.**
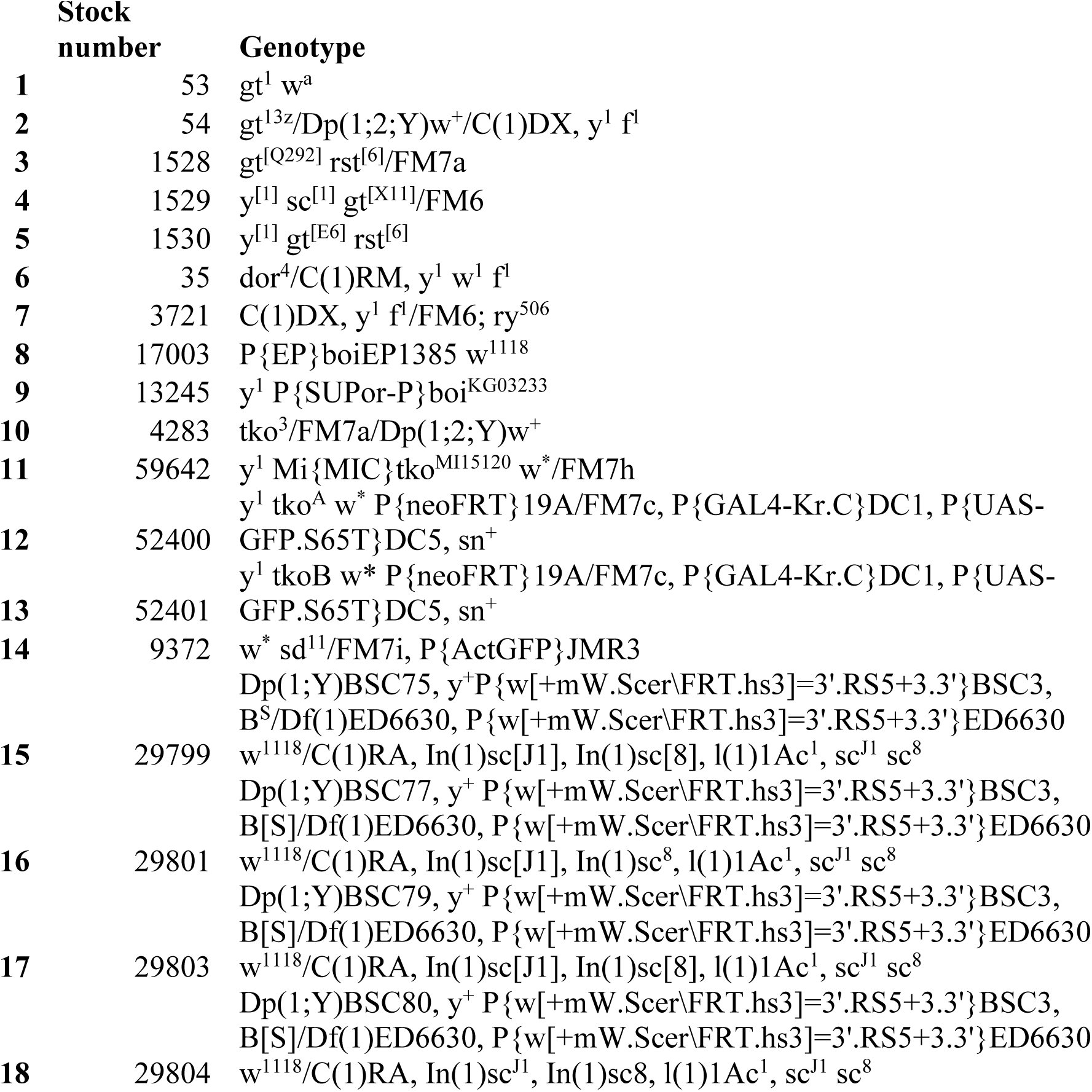
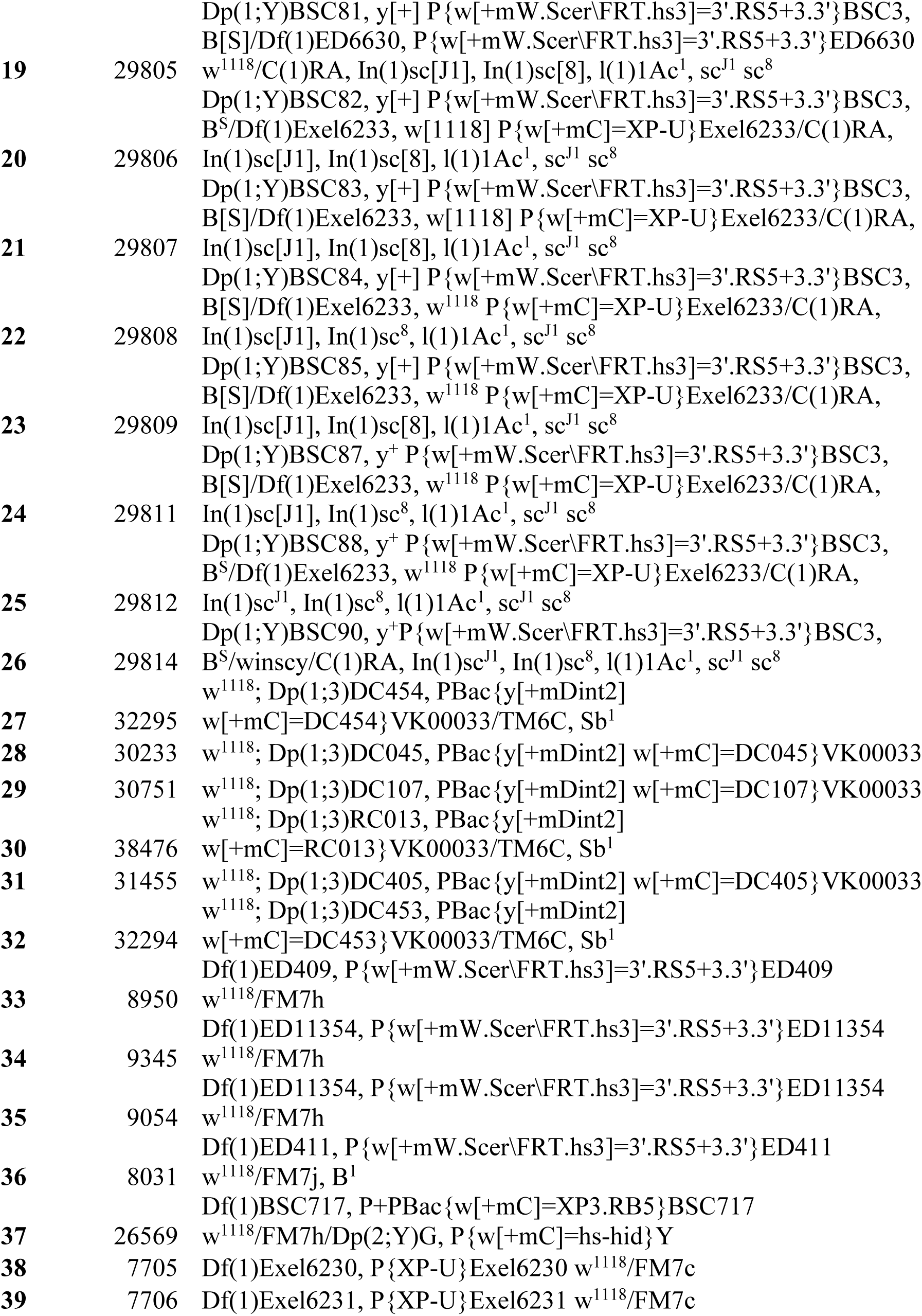
Mutant stocks used in this study.

**TABLE S4.**
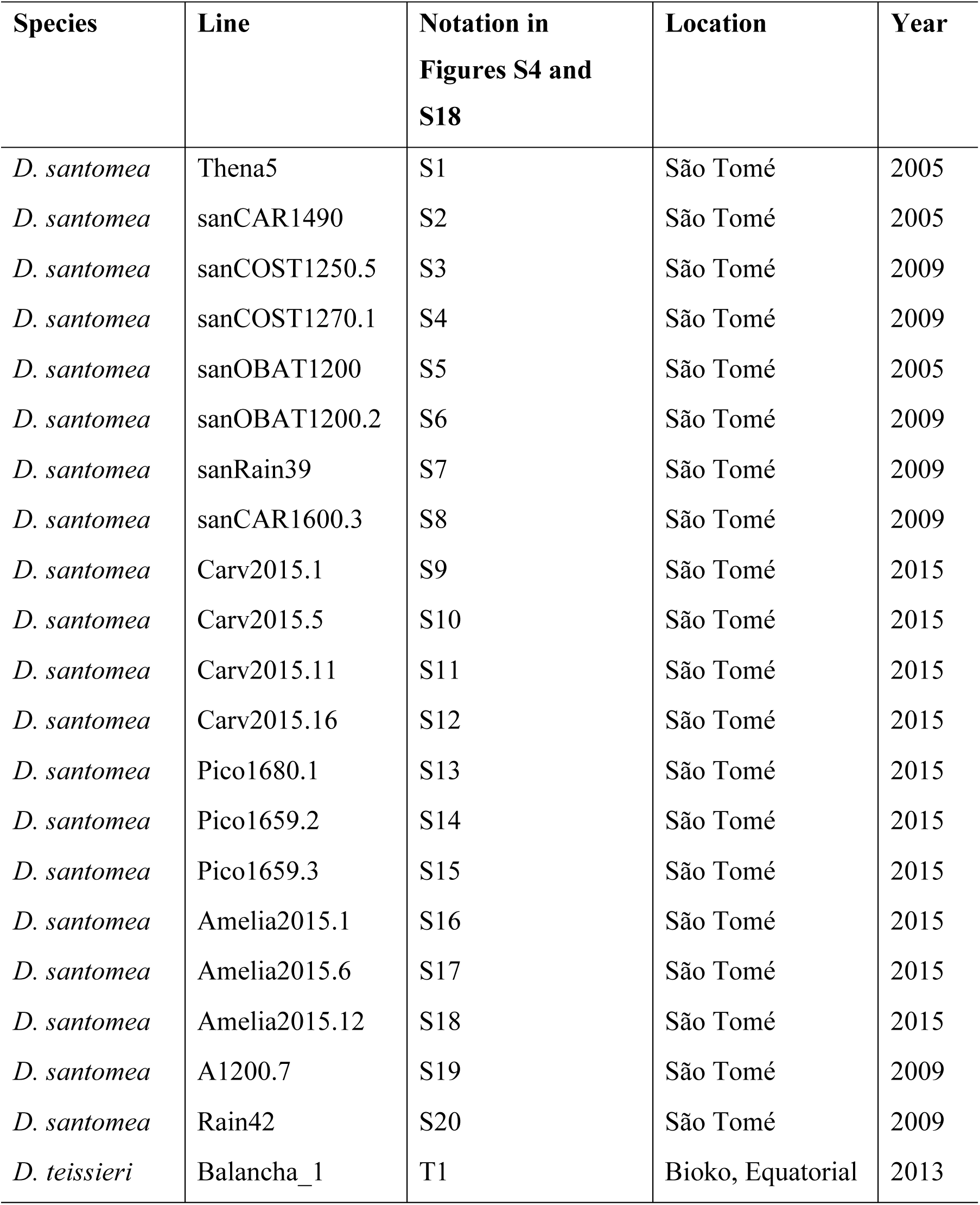
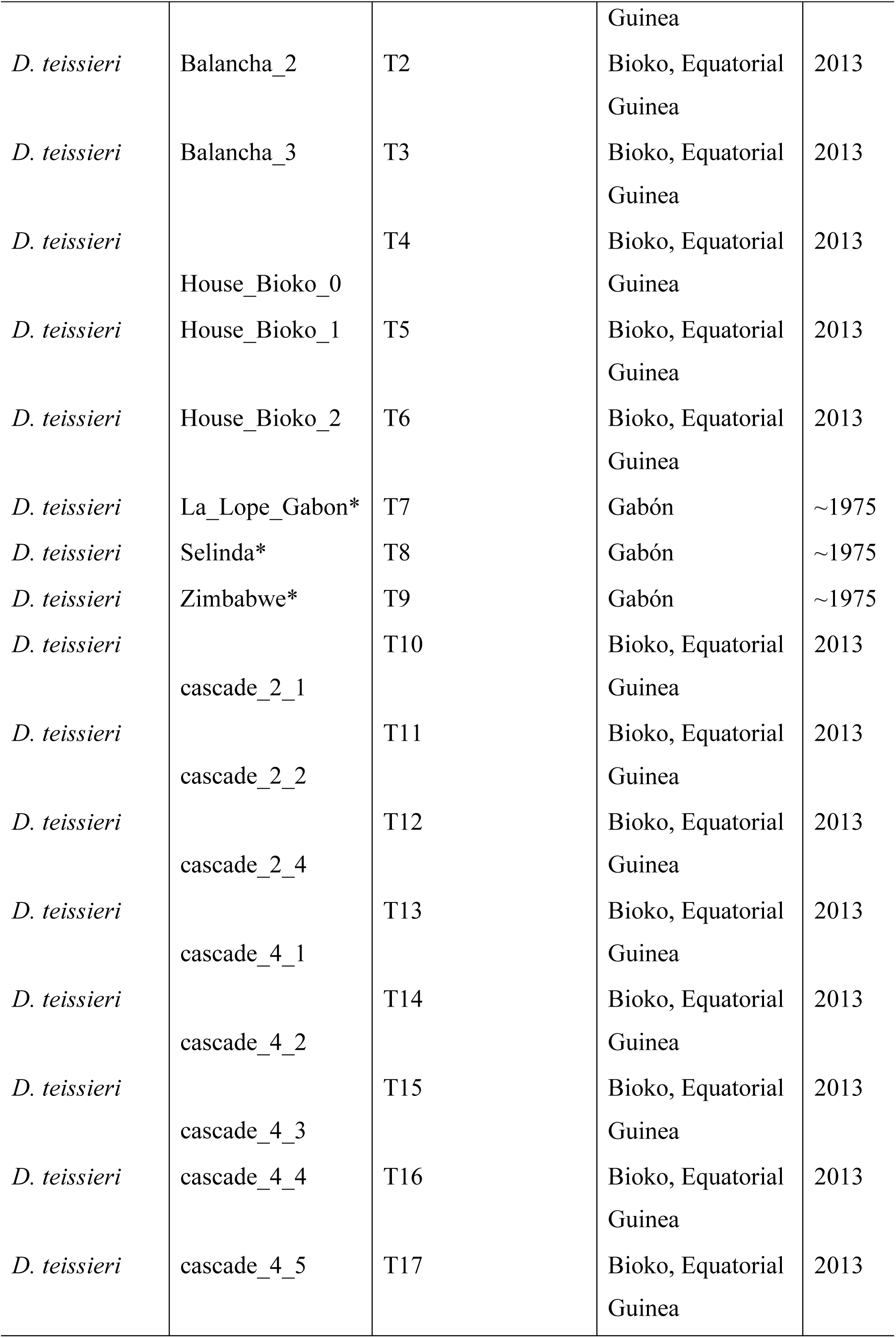
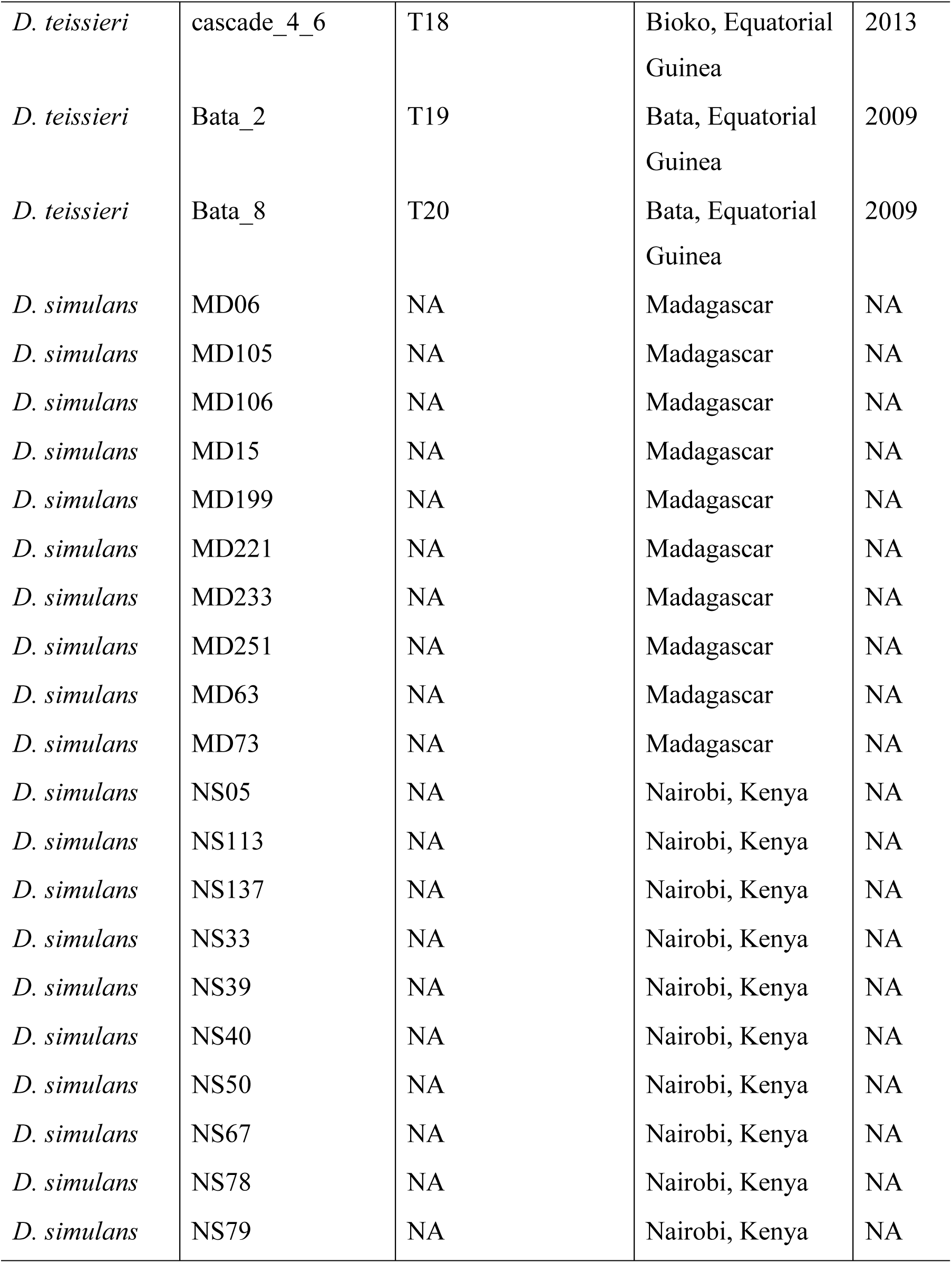
Isofemale lines from four species used to assess whether HI in *mel/san* hybrids was a species-specific phenomenon or a line specific phenomenon. All lines were collected by D. R. Matute with the exception of the three *D. teissieri* lines marked with an asterisk. Those three lines were donated by J.R. David. MISSING THE MELANOGASTER LINES (ALSO SIM>????)

**TABLE S5.**
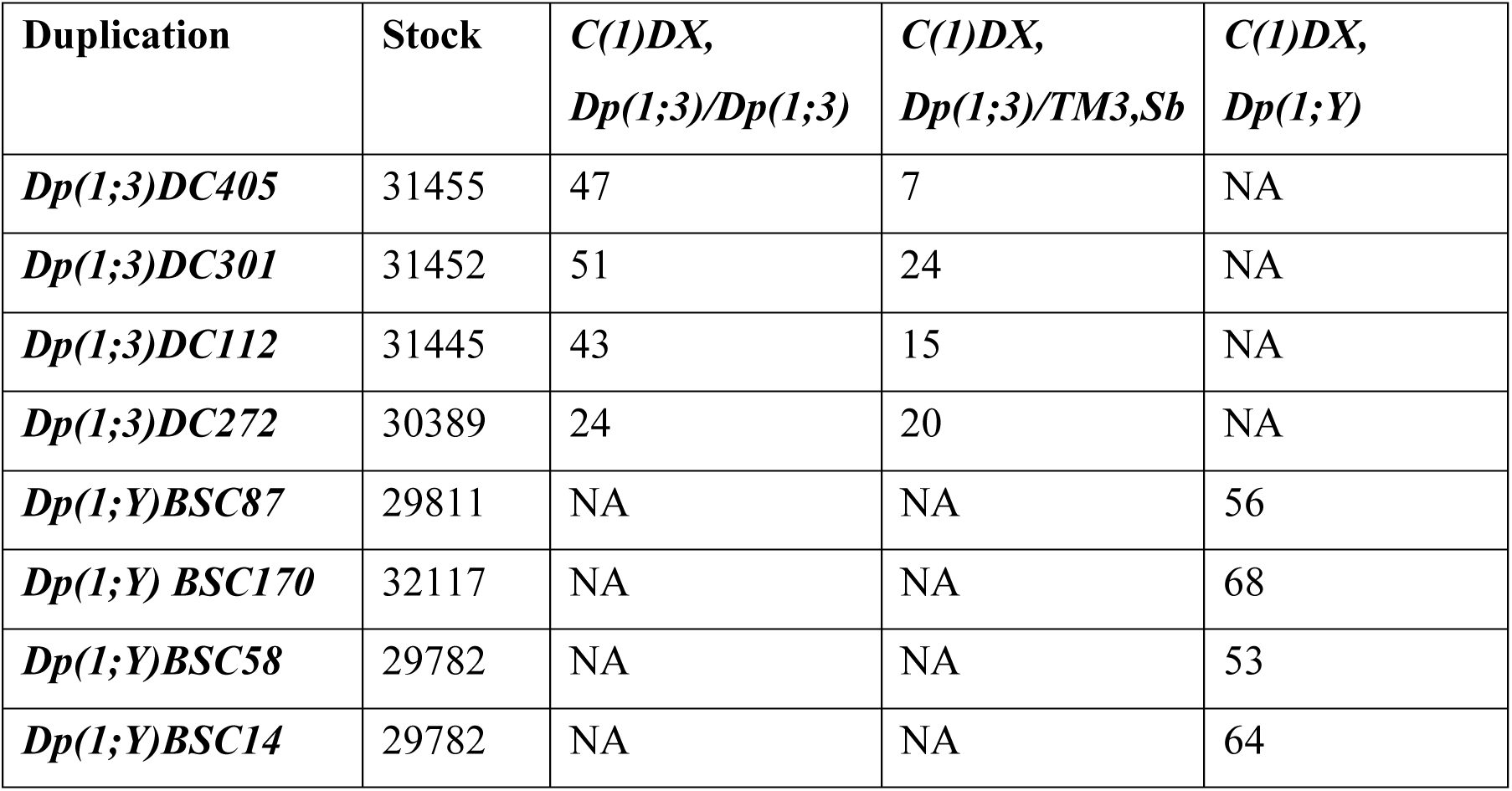
*C(1)DX, Dp(1;3)/TM3* lay few eggs than *C(1)DX, Dp(1;Y)* females in conspecific *mel* matings. Each column shows the median number of eggs produced by a singly-mated female when mated to a *w^1118^* male over 4 days (*N* = 15 females).

**TABLE S6.**
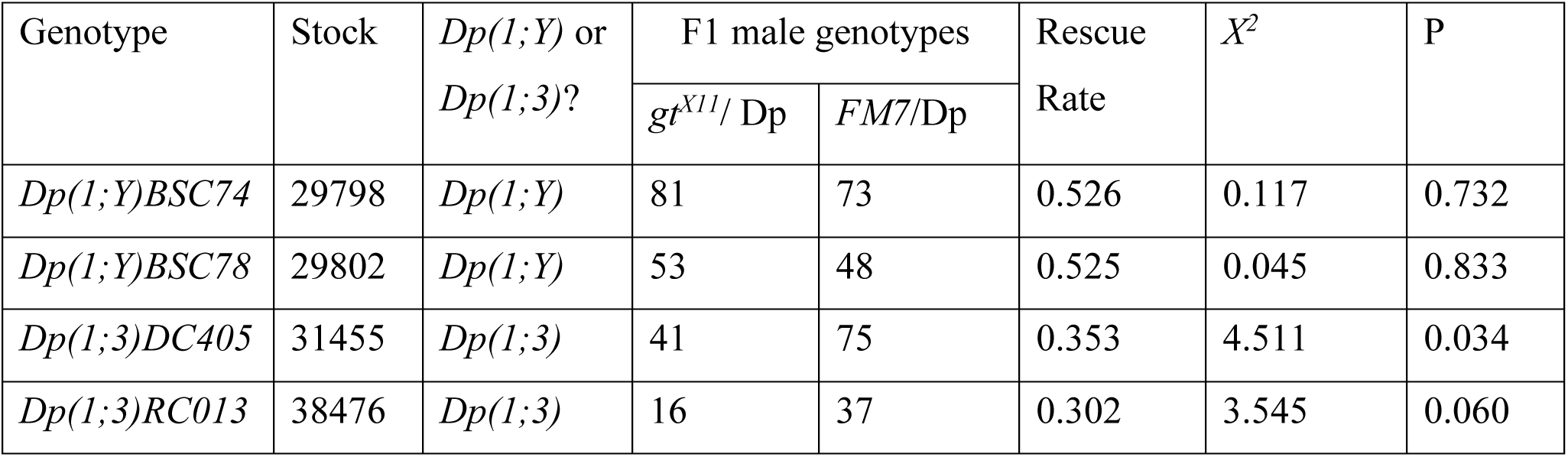
*Y*-linked duplications of the *X*-chromosome rescue a *gt_mel_* null allele more effectively than duplications on the third chromosome in a pure *mel* background. All crosses involved *gt_mel_^X11^/FM7* females and hemizygote males for two different types of duplications (*Dp(1;Y)* or *Dp(1;3)*). Cells show the total progeny from 15 crosses. In the case of *Dp(1;Y)* duplications, the father had a *X_mel_/Y_mel_Dp(1;Y)* genotype. In the case of *Dp(1;3*) duplication, the father had a *X_mel_/Y_mel_*, *Dp(1,3)/ Dp(1,3)* genotype. I only scored *gt^X11^*-carriers.

**TABLE S7.**
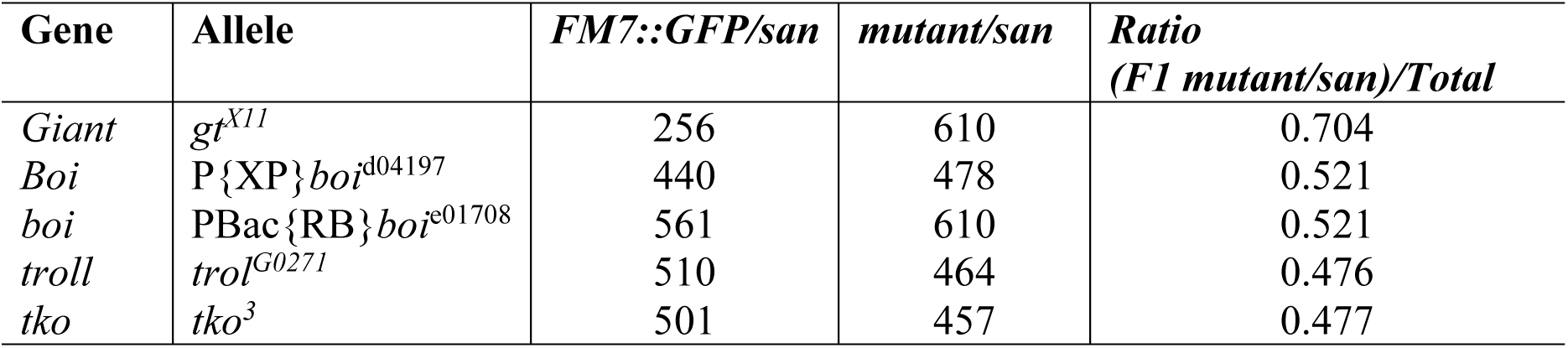
Complementation mapping using loss-of function and hypomoprhic alleles show that *gt* mutants are the only alleles in cytological region 3A3 that lead to an increase of female viability in *mel/san* hybrid. *mutant/san* refers to the number of F1 hybrids carrying the mutant allele. *gt^X11^* is a loss of function (amorphic) allele (2, 40). Both *boi^d04197^* ^and^ *boi^e01708^* shows significantly lower RNA production than wild-type flies (41). *trol^G0271^*is a hypomorph (42). *tko^3^* is a null allele. *tko^3^*/*tko^3^* homozygote females and *tko^3^* hemizygote males die as larvae but are able to complete embryogenesis (43).

**TABLE S8.** Haplotypes associated with HI in *mel/san* hybrids. This table only shows autosomal haplotypes besides the one that contains *tll^mel^* (.xls file).

**TABLE S9.**
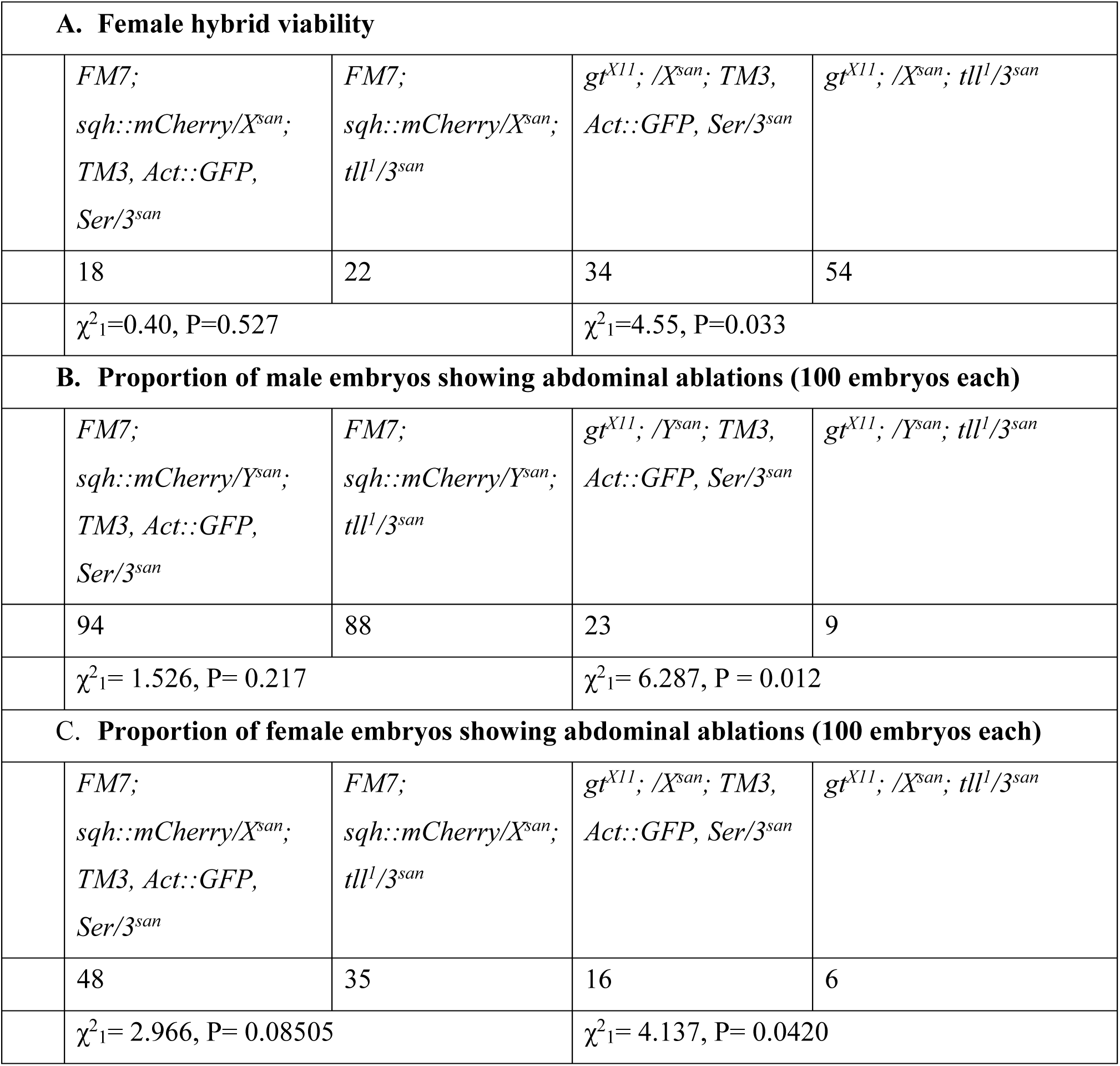
*tll^mel^*exacerbates the defects caused by *gt^mel^* in *mel/san* hybrids as revealed by a second *tll* mutant. The analysis is similar to that shown in Table 1. **A.** Female viability **B.** Frequency of abdominal ablations in hybrid males. **C.** Frequency of abdominal ablations in hybrid females.

| A. Female hybrid viability |  |  |  |  |
| --- | --- | --- | --- | --- |
|  | *FM7;*<br>*sqh::mCherry/X<sup>san</sup>;*<br>*TM3, Act::GFP,*<br>*Ser/3<sup>san</sup>* | *FM7;*<br>*sqh::mCherry/X<sup>san</sup>;*<br>*tll<sup>l</sup>/3<sup>san</sup>* | *gt<sup>X11</sup>; /X<sup>san</sup>; TM3,*<br>*Act::GFP, Ser/3<sup>san</sup>* | *gt<sup>X11</sup>; /X<sup>san</sup>; tll<sup>l</sup>/3<sup>san</sup>* |
|  | 18 | 22 | 34 | 54 |
| | $\chi^2_1=0.40$, P=0.527 | | $\chi^2_1=4.55$, P=0.033 | |
| B. Proportion of male embryos showing abdominal ablations (100 embryos each) |  |  |  |  |
|  | *FM7;*<br>*sqh::mCherry/Y<sup>san</sup>;*<br>*TM3, Act::GFP,*<br>*Ser/3<sup>san</sup>* | *FM7;*<br>*sqh::mCherry/Y<sup>san</sup>;*<br>*tll<sup>l</sup>/3<sup>san</sup>* | *gt<sup>X11</sup>; /Y<sup>san</sup>; TM3,*<br>*Act::GFP, Ser/3<sup>san</sup>* | *gt<sup>X11</sup>; /Y<sup>san</sup>; tll<sup>l</sup>/3<sup>san</sup>* |
|  | 94 | 88 | 23 | 9 |
| | $\chi^2_1= 1.526$, P= 0.217 | | $\chi^2_1= 6.287$, P = 0.012 | |
| C. Proportion of female embryos showing abdominal ablations (100 embryos each) |  |  |  |  |
|  | *FM7;*<br>*sqh::mCherry/X<sup>san</sup>;*<br>*TM3, Act::GFP,*<br>*Ser/3<sup>san</sup>* | *FM7;*<br>*sqh::mCherry/X<sup>san</sup>;*<br>*tll<sup>l</sup>/3<sup>san</sup>* | *gt<sup>X11</sup>; /X<sup>san</sup>; TM3,*<br>*Act::GFP, Ser/3<sup>san</sup>* | *gt<sup>X11</sup>; /X<sup>san</sup>; tll<sup>l</sup>/3<sup>san</sup>* |
|  | 48 | 35 | 16 | 6 |
| | $\chi^2_1= 2.966$, P= 0.08505 | | $\chi^2_1= 4.137$, P= 0.0420 | |

**TABLE S10.** Double mutant analysis of survival in hybrids between *D. melanogaster* females and three males from four different species (*D. melanogaster*, *D. simulans*, *D. mauritiana*, and *D. teissieri*). Individuals that are hemizygote for both alleles do not show different viability from individuals that are hemizygote for one allele and heterozygote for the other one, or heterozygote for both alleles.

| *tll<sup>l</sup>* |  |  |  |  |  |  |
| --- | --- | --- | --- | --- | --- | --- |
| | *FM7; sqh::mCherry/X<sup>san</sup>; TM3, Act::GFP, Ser/3<sup>san</sup>* | *FM7; sqh::mCherry/X<sup>s</sup><sub>an</sub>; tll<sup>l</sup>/3<sup>san</sup>* | *gt<sup>X11</sup>; /X<sup>san</sup>; TM3, Act::GF P, Ser/3<sup>san</sup>* | *gt<sup>X11</sup>; /X<sup>san</sup>; tll<sup>l</sup>/3<sup>sa</sup><sub>n</sub>* | $\chi^2$, df =3 | P-value |
| *D. melanogaster* | 95 | 104 | 121 | 98 | 1.869<br>5 | 0.599<br>9 |
| *D. simulans* | 34 | 31 | 28 | 29 | 0.338<br>64 | 0.952<br>6 |
| *D. mauritiana* | 52 | 41 | 39 | 41 | 1.143<br>6 | 0.766<br>6 |
| *D. teissieri* | 19 | 22 | 15 | 20 | 0.715<br>74 | 0.869<br>5 |
| *tll<sup>ΔGFP</sup>* |  |  |  |  |  |  |
| | *FM7; sqh::mCherry/X<sup>san</sup>; TM3, Act::GFP, Ser/3<sup>san</sup>* | *FM7; sqh::mCherry/X<sup>s</sup><sub>an</sub>; tll<sup>l</sup>/3<sup>san</sup>* | *gt<sup>X11</sup>; /X<sup>san</sup>; TM3, Act::GF P, Ser/3<sup>san</sup>* | *gt<sup>X11</sup>; /X<sup>san</sup>; tll<sup>l</sup>/3<sup>sa</sup><sub>n</sub>* | $\chi^2$ | |
| *D. melanogaster* | 104 | 87 | 99 | 78 | 2.142<br>8 | 0.543<br>3 |
| *D. simulans* | 42 | 36 | 39 | 44 | 0.460<br>75 | 0.927<br>4 |
| *D. mauritiana* | 51 | 38 | 49 | 43 | 1.181<br>5 | 0.757<br>5 |
| *D. teissieri* | 20 | 25 | 21 | 26 | 0.567<br>22 | 0.903<br>9 |

**TABLE S11.**
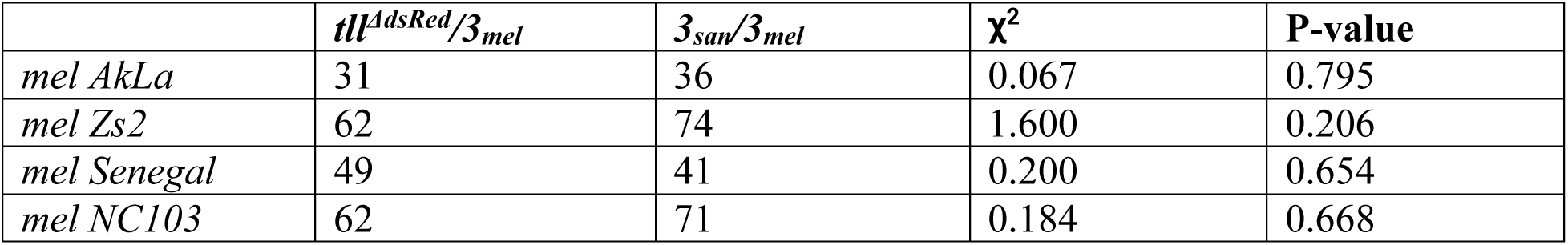
A *tll_san_^ΔdsRed^* has no effect on HI in crosses between male-carriers and *mel* females from four different backgrounds.

| | $tll^{dsRed}/3_{mel}$ | $3_{san}/3_{mel}$ | $\chi^2$ | P-value |
| --- | --- | --- | --- | --- |
| *mel AkLa* | 31 | 36 | 0.067 | 0.795 |
| *mel Zs2* | 62 | 74 | 1.600 | 0.206 |
| *mel Senegal* | 49 | 41 | 0.200 | 0.654 |
| *mel NC103* | 62 | 71 | 0.184 | 0.668 |

**TABLE S12.**
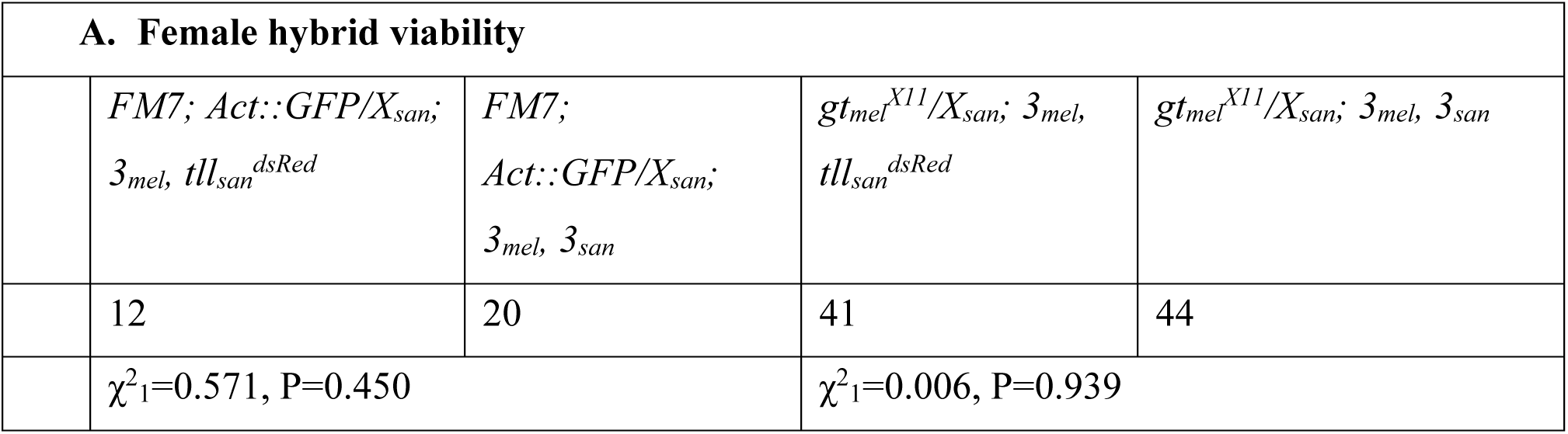
Abrogating the *tll_san_* allele has no viability effect in *gt_mel_^X11^ mel*/san hybrids.

**TABLE S13.**
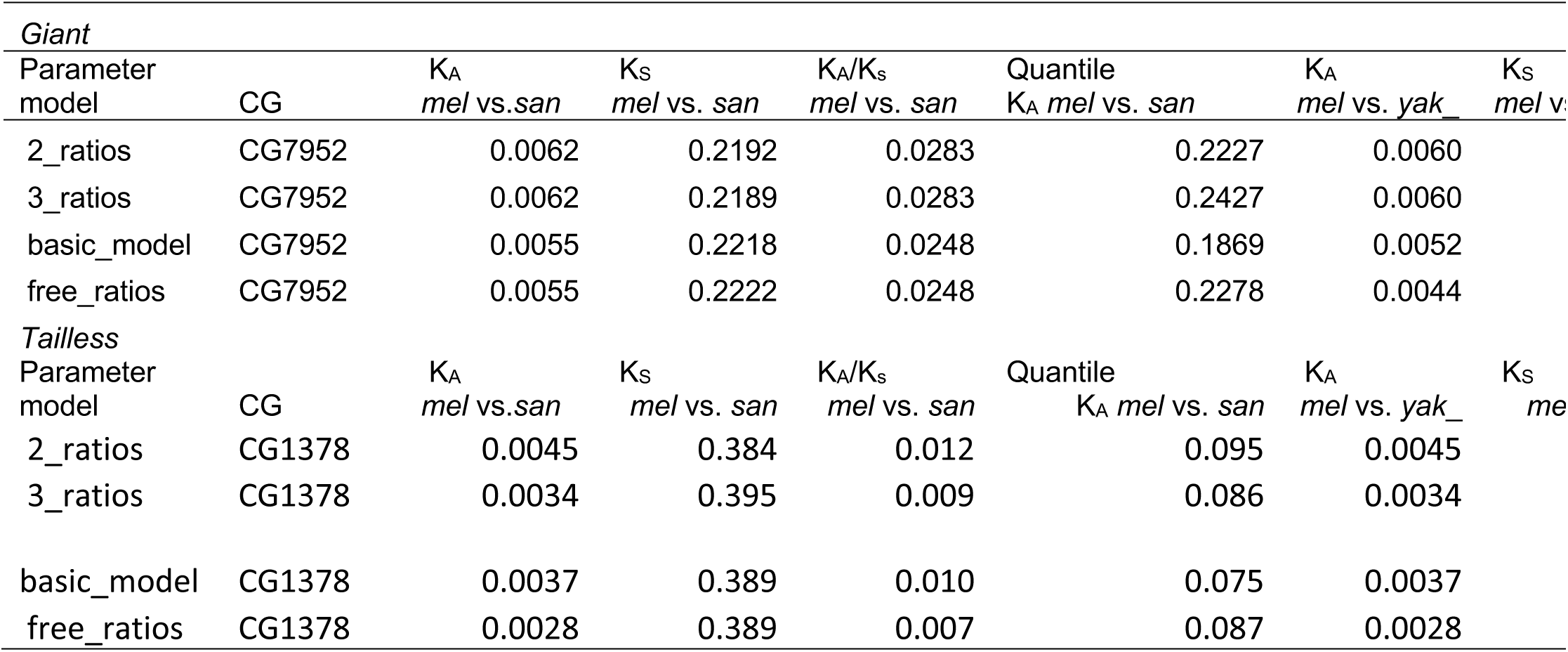
*gt* and *tll* are slowly evolving genes as measured by the rate of aminoacid substitutions. I show K_A_/K_S_ values for two species comparisons (*D. melanogaster* vs. *D. santomea*, and *D. melanogaster* vs. *D. yakuba*) and the quantiles of the K_A_/K_S_ values compared to the rest of the genome. I used four different parametrizations in PAML (listed in the first column). *D. melanogaster*: *mel*, *D. santomea*: *san*; *D. yakuba*: *yak*.

| *Giant* |  |  |  |  |  |  |  |
| --- | --- | --- | --- | --- | --- | --- | --- |
| Parameter model | CG | $K_A$<br>*mel* vs.*san* | $K_S$<br>*mel* vs. *san* | $K_A/K_S$<br>*mel* vs. *san* | Quantile<br>$K_A$ *mel* vs. *san* | $K_A$<br>*mel* vs. *yak* | $K_S$<br>*mel* vs. *yak* |
| 2_ratios | CG7952 | 0.0062 | 0.2192 | 0.0283 | 0.2227 | 0.0060 |  |
| 3_ratios | CG7952 | 0.0062 | 0.2189 | 0.0283 | 0.2427 | 0.0060 |  |
| basic_model | CG7952 | 0.0055 | 0.2218 | 0.0248 | 0.1869 | 0.0052 |  |
| free_ratios | CG7952 | 0.0055 | 0.2222 | 0.0248 | 0.2278 | 0.0044 |  |
| *Tailless* |  |  |  |  |  |  |  |
| Parameter model | CG | $K_A$<br>*mel* vs.*san* | $K_S$<br>*mel* vs. *san* | $K_A/K_S$<br>*mel* vs. *san* | Quantile<br>$K_A$ *mel* vs. *san* | $K_A$<br>*mel* vs. *yak* | $K_S$<br>*mel* vs. *yak* |
| 2_ratios | CG1378 | 0.0045 | 0.384 | 0.012 | 0.095 | 0.0045 |  |
| 3_ratios | CG1378 | 0.0034 | 0.395 | 0.009 | 0.086 | 0.0034 |  |
| basic_model | CG1378 | 0.0037 | 0.389 | 0.010 | 0.075 | 0.0037 |  |
| free_ratios | CG1378 | 0.0028 | 0.389 | 0.007 | 0.087 | 0.0028 |  |

**TABLE S14.**
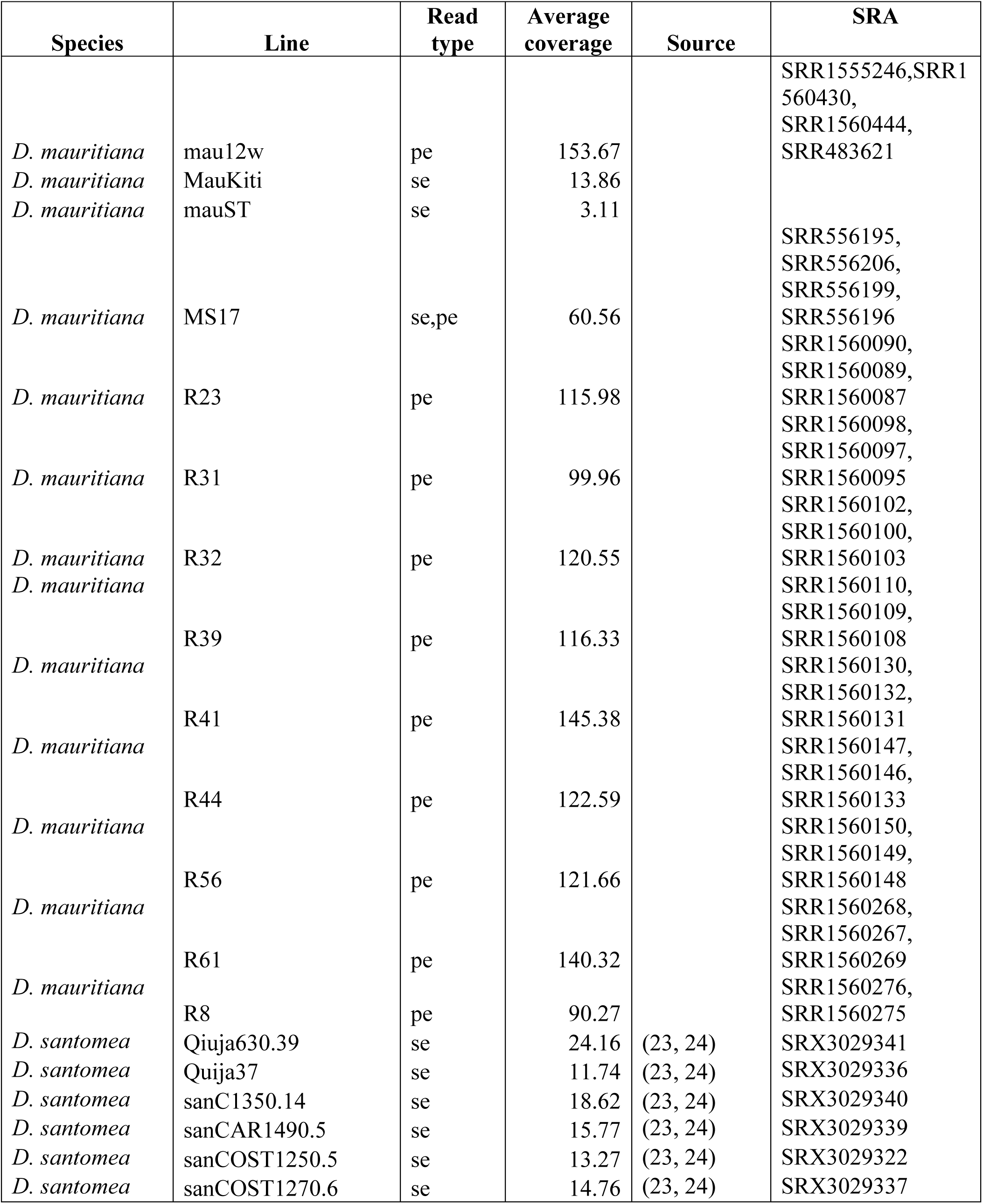
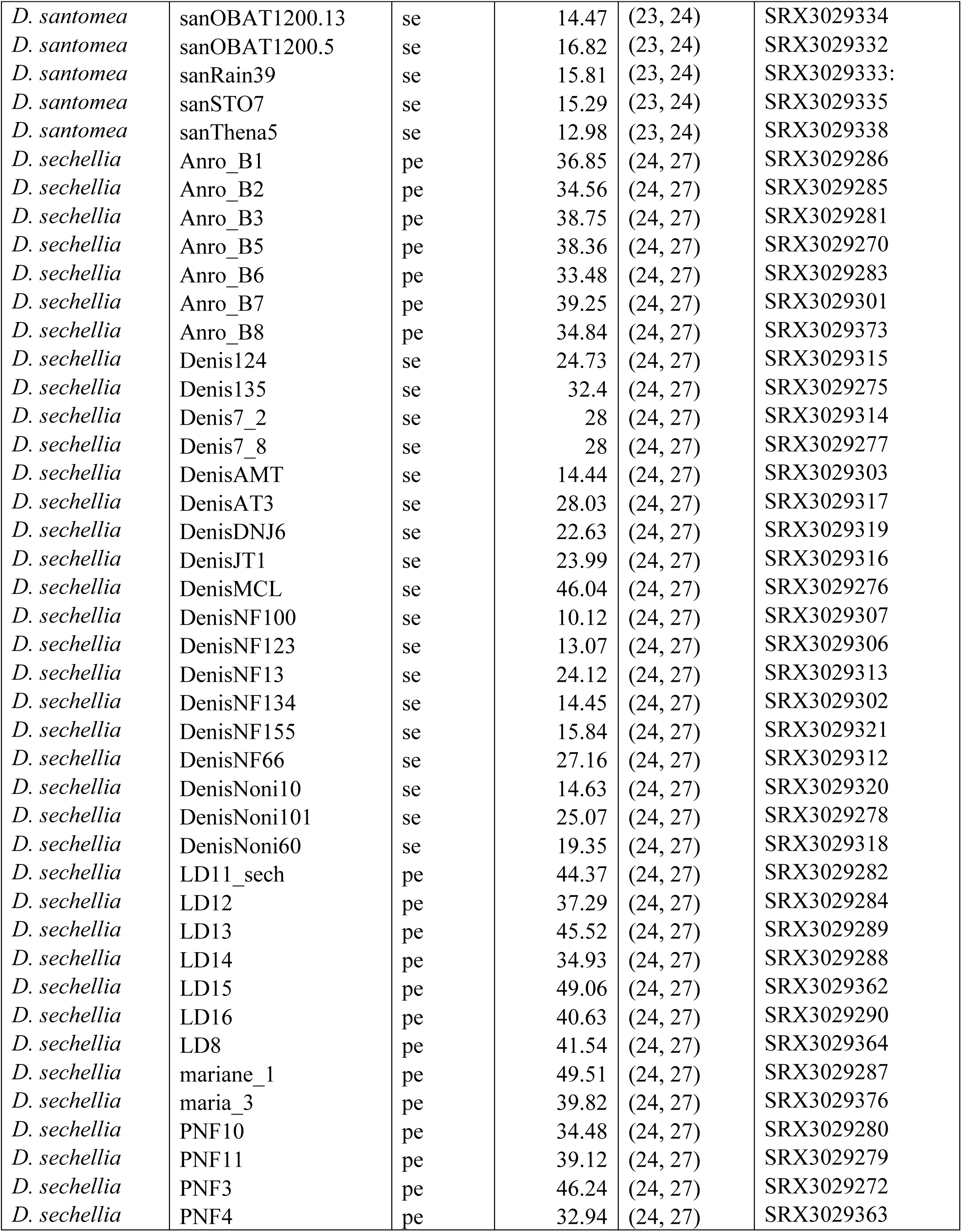
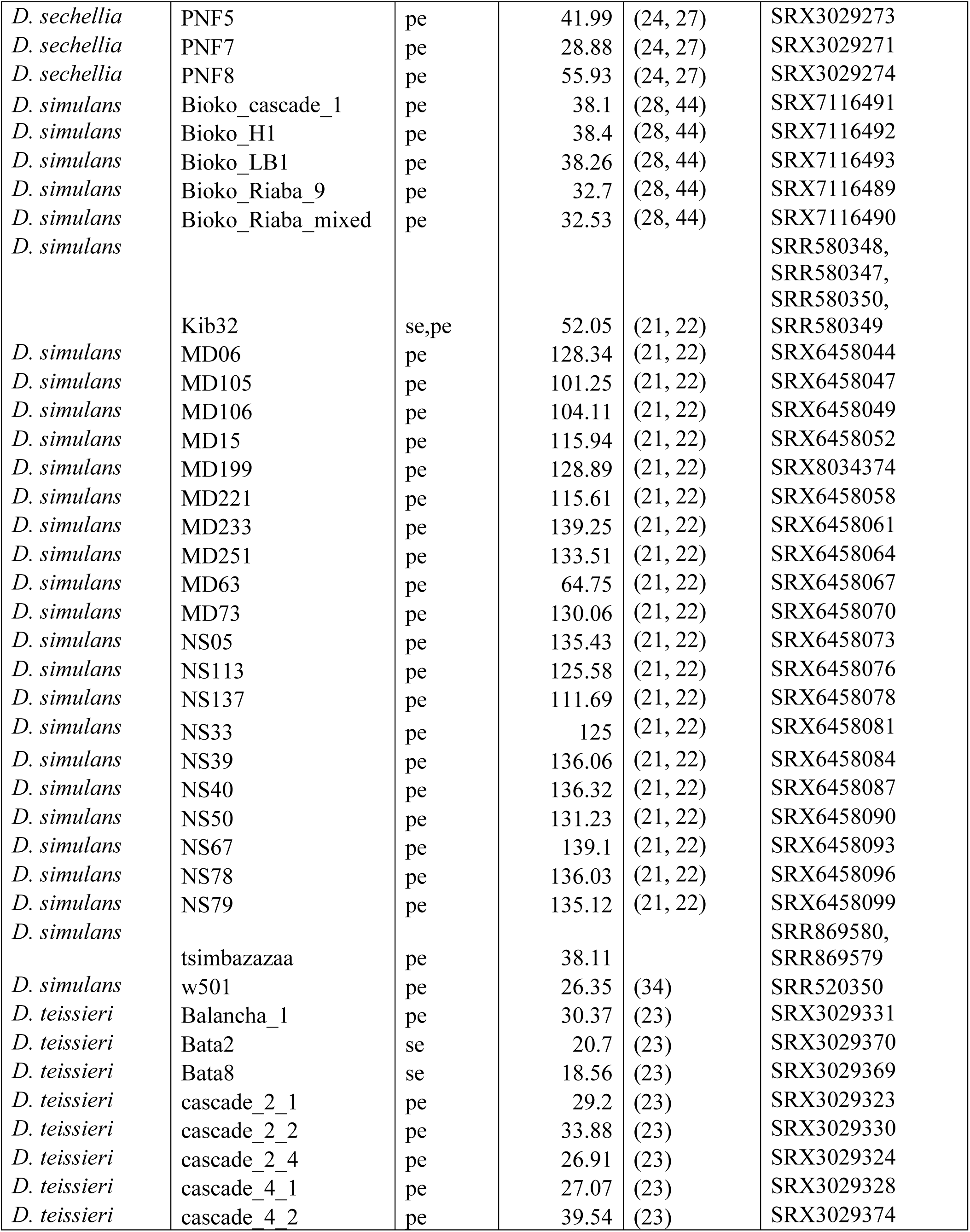
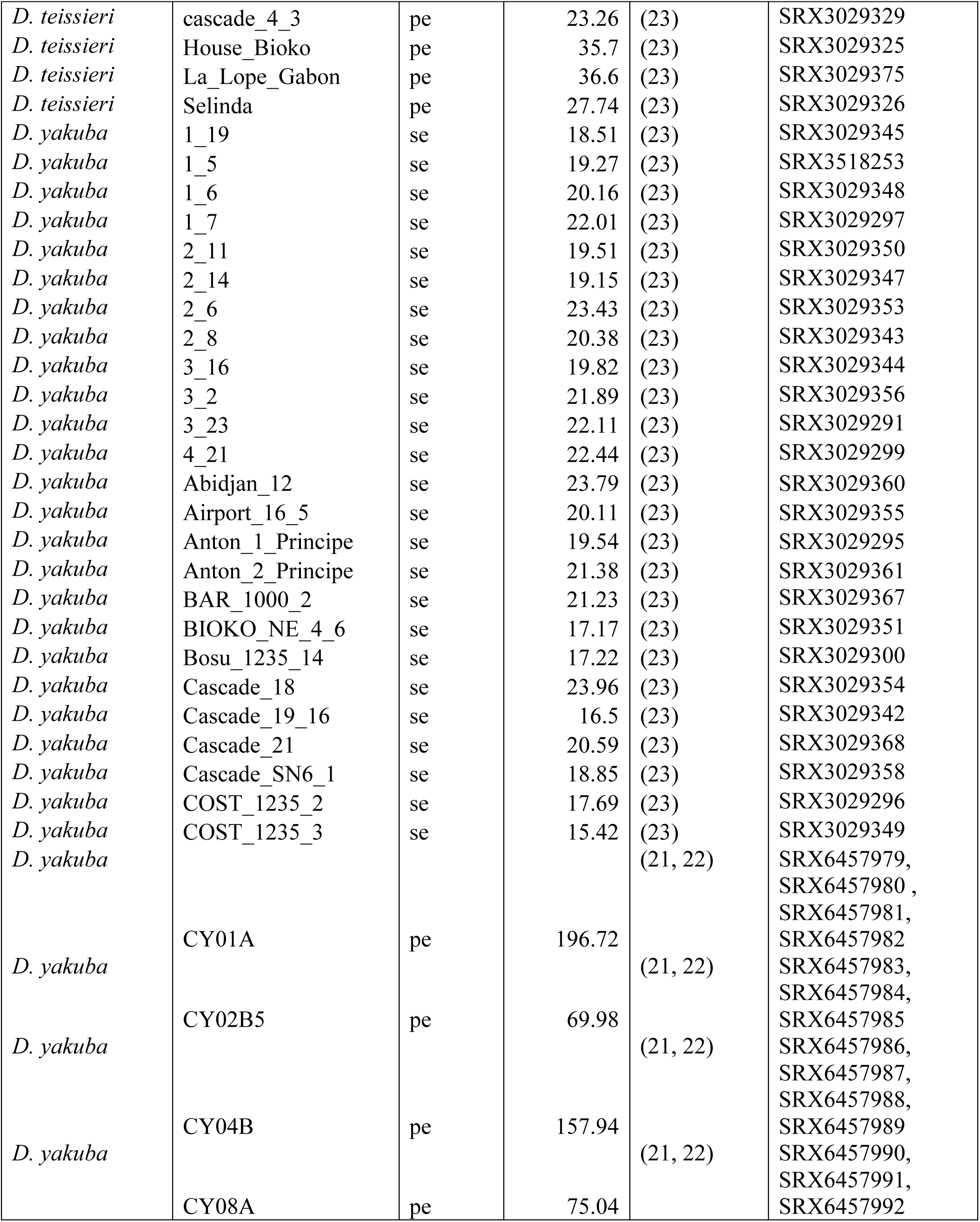
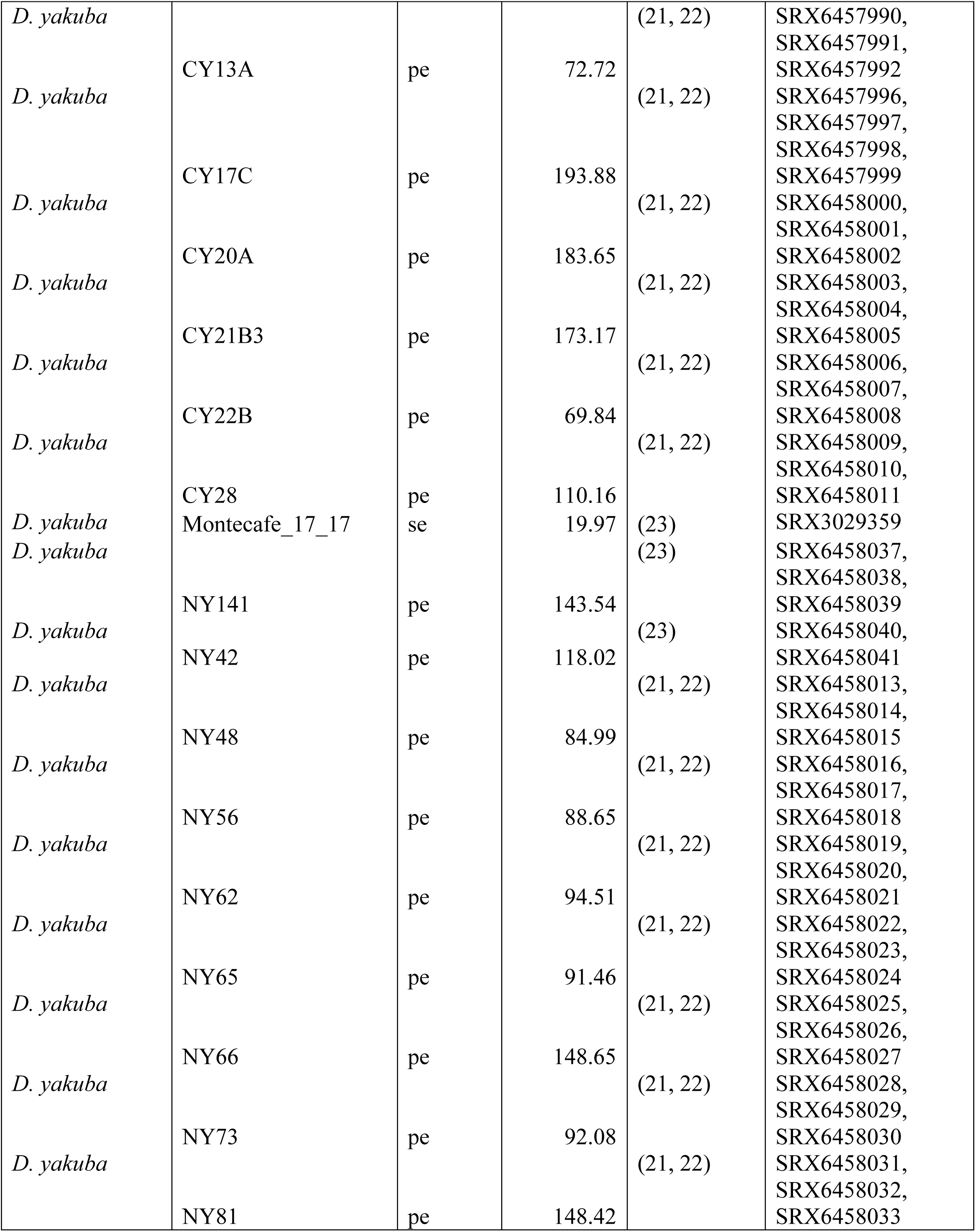
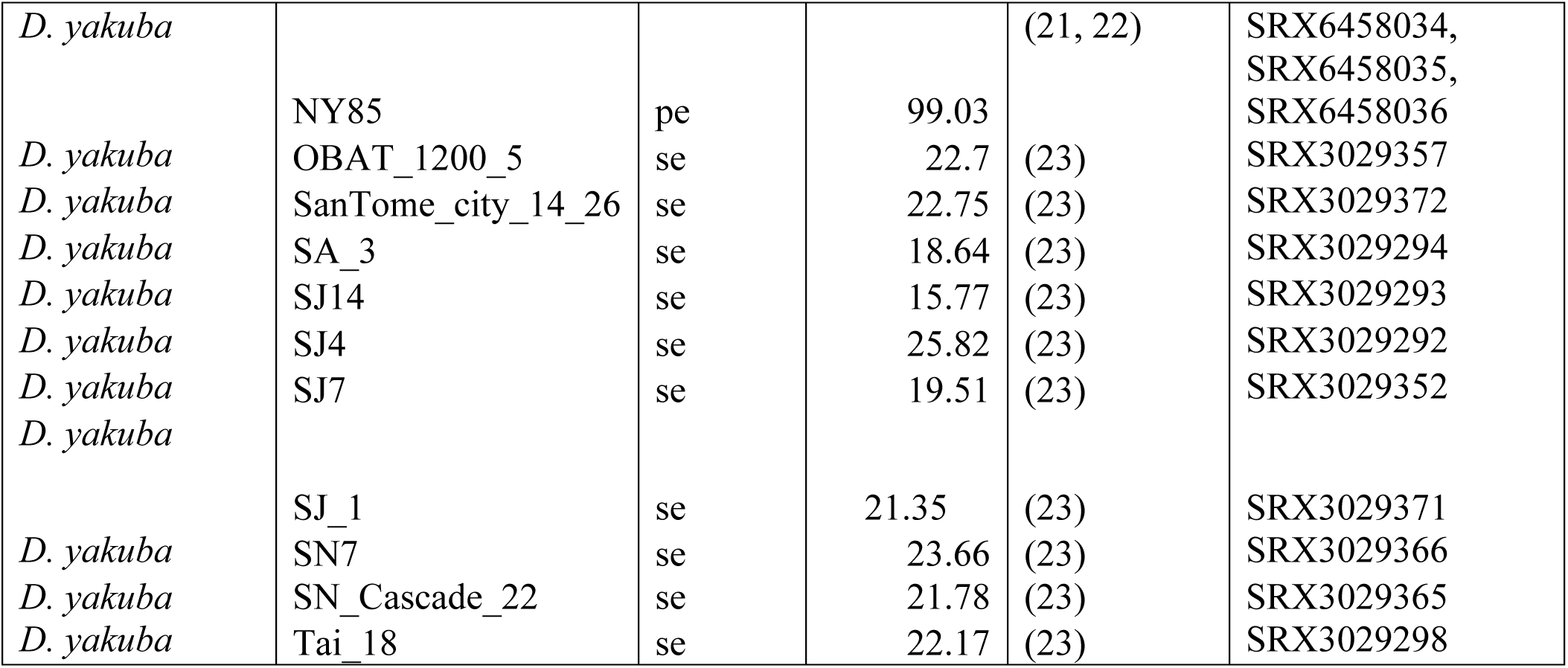
Sequencing details and coverage for all the lines included in this study.

### SUPPLEMENTARY FIGURES

**FIGURE S1.**
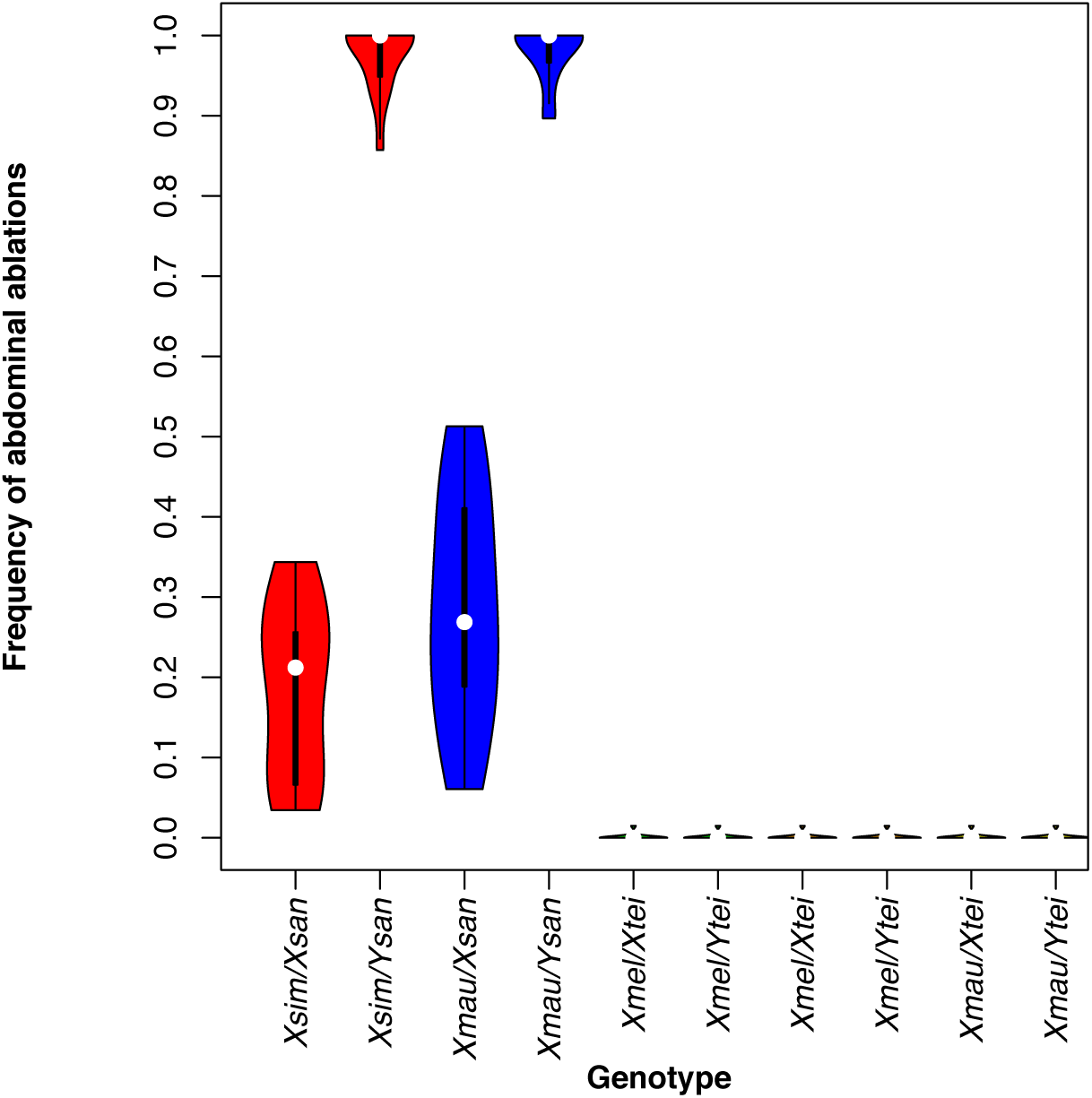
*X_sim_* and *X_mau_* cause abdominal ablations in hybrid males with *D. santomea*. Hybrid males from the **♀***sim ×* **♂***san* and **♀***mau ×* **♂** *san* crosses show high frequency of abdominal ablations similar to those observed in **♀***mel ×* **♂** *san* hybrids (Figures 1 and 2C). Hybrid females from the same crosses show a lower frequency of ablations. The nature of the defect in these hybrid males is identical to that seen in *mel/san* hybrid males, a characteristic ablation of abdominal segments (Figure 1C).

**FIGURE S2.**
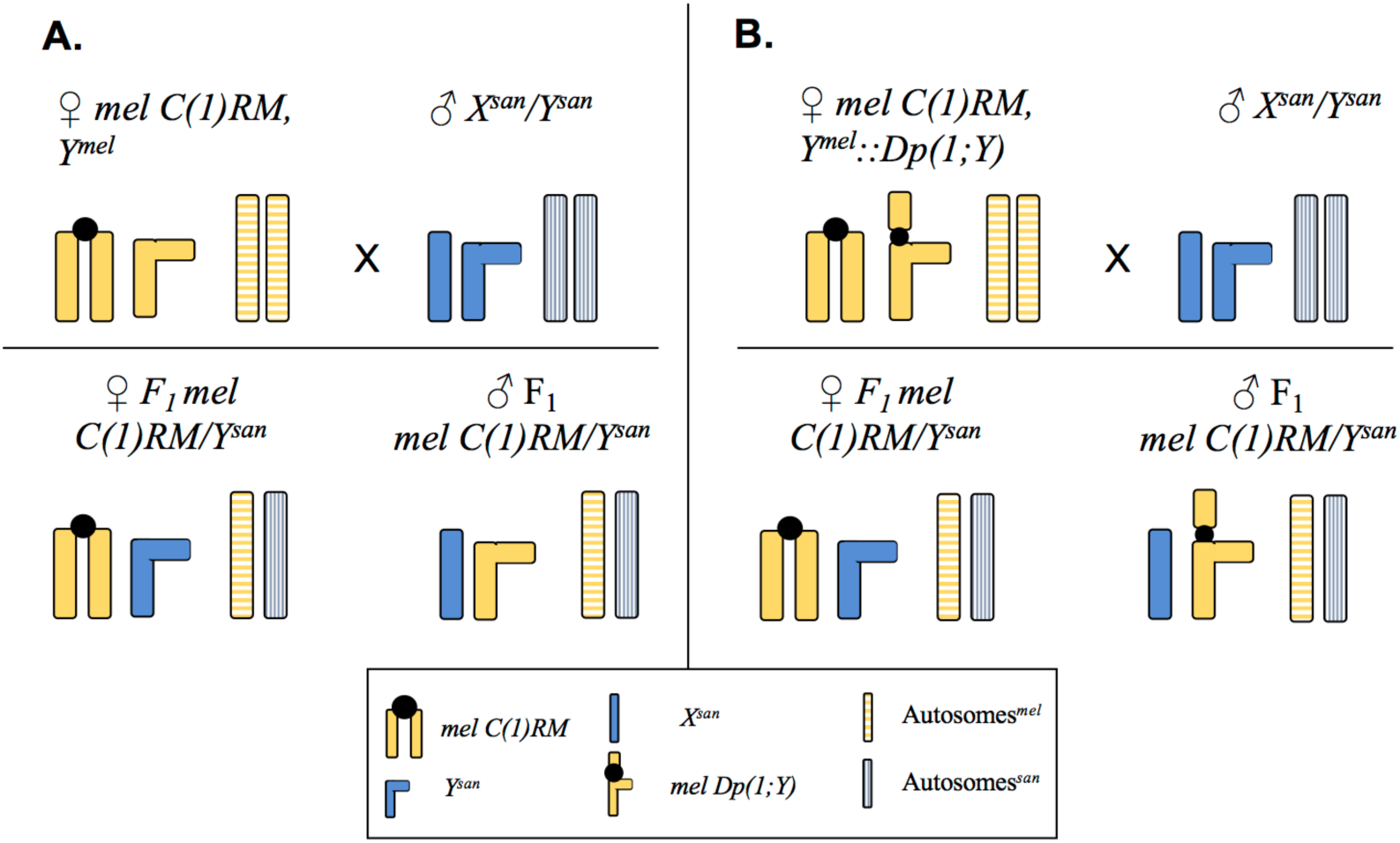
Crossing design to assess whether of *Y*-linked pieces of the *X_mel_* chromosome cause hybrid inviability and abdominal ablations. Blue bars represent *D. santomea* chromosomes; yellow bars represent *D. melanogaster* chromosomes. Solid colors: sex chromosomes, stripped bars: autosomes. This approach is a modified version of (12, 14, 45).

**FIGURE S3.**
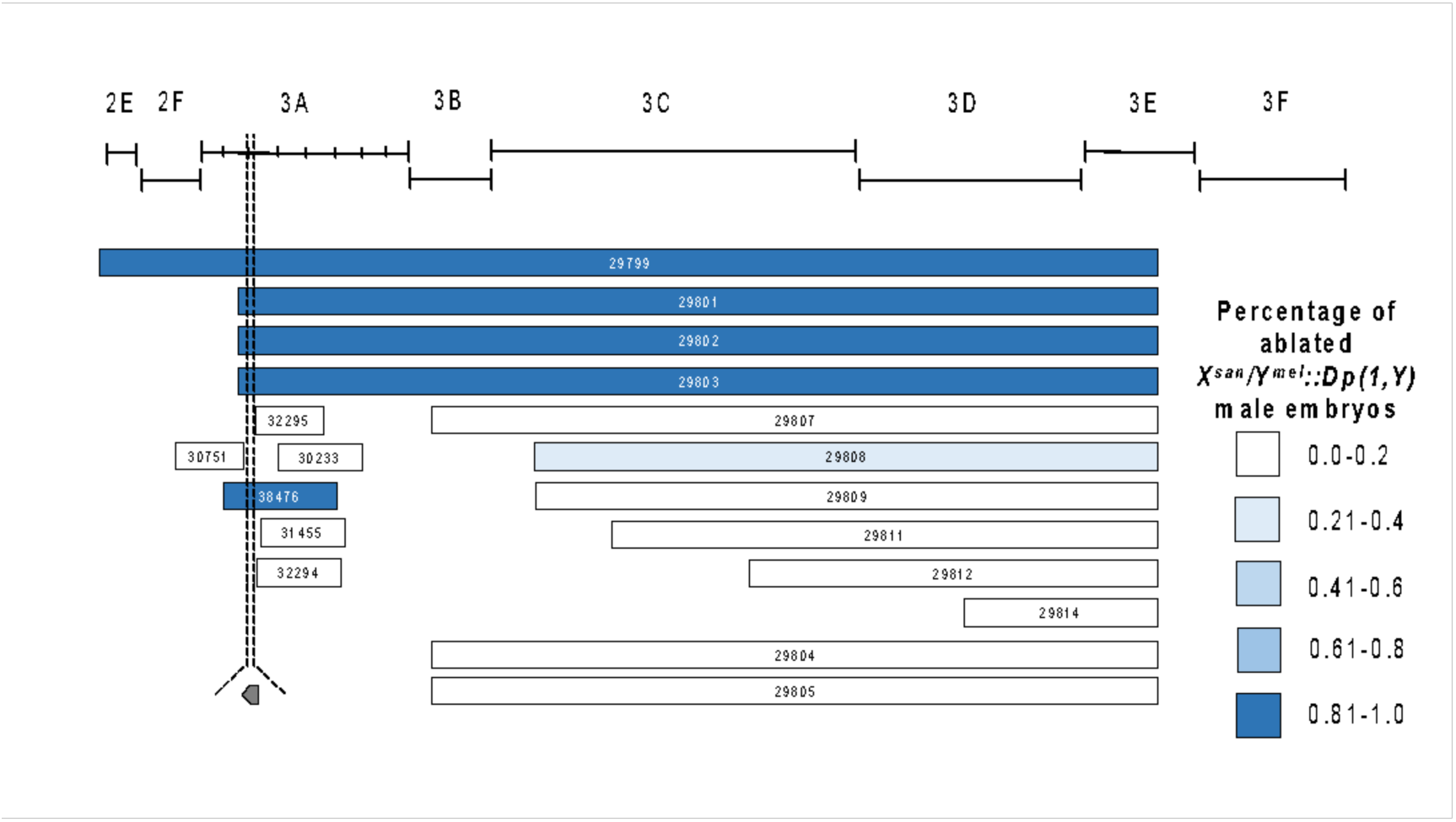
Frequency of abdominal ablations caused by the *X_mel_* translocations shown in Figure 2 in *X_san_/Y_mel_* hybrid males. Each Bloomington stock number is shown within the bar. The precise genotype of each stock is shown in Table S3.

**FIGURE S4.**
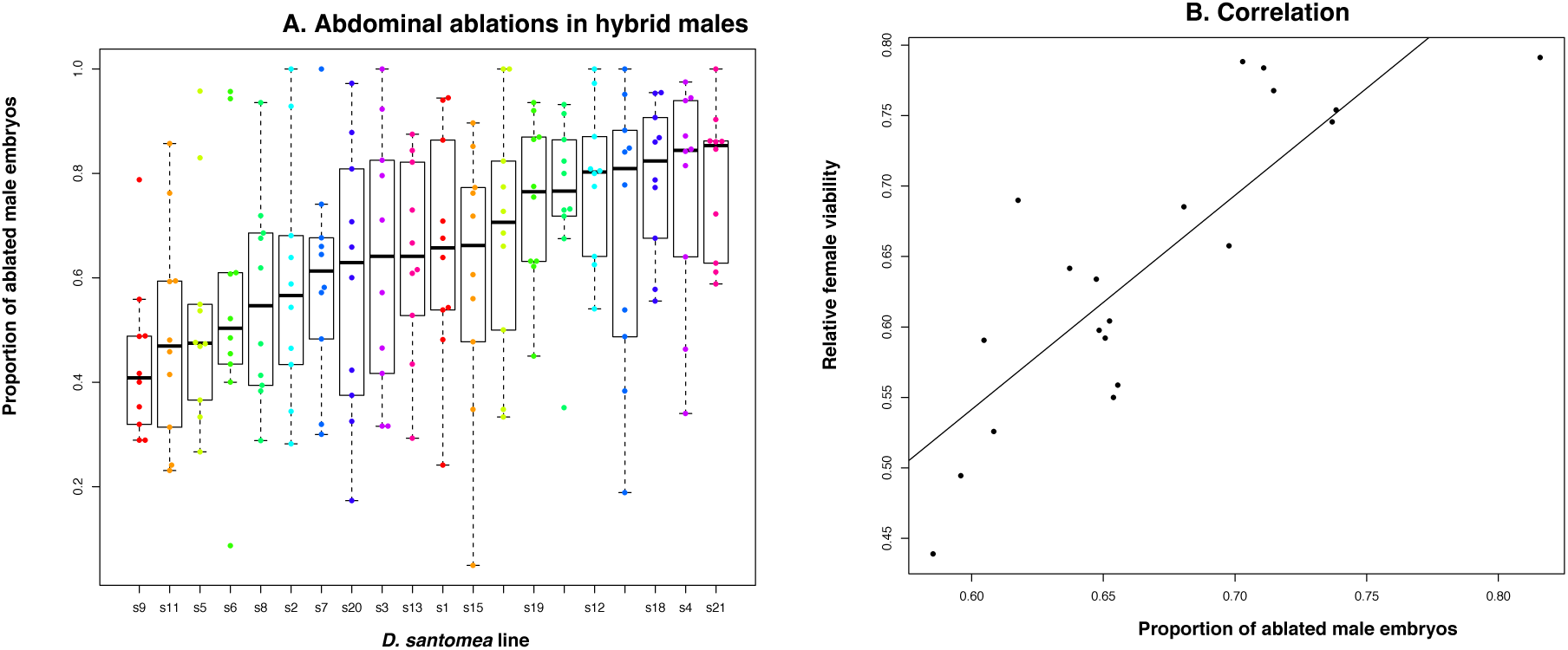
The presence of *gt_mel_* causes HI in *mel/san* hybrids produced with all *D. santomea* lines but the magnitude of the inviability varies. I measured frequency of abdominal ablations in hybrid *X_mel_/Y_san_* males (**A**). The magnitude of the frequency of abdominal ablations and of hybrid female inviability are correlated among lines (panel **B**; ρ = 0.1734, P = 0.0141). Boxes in the boxplot are ordinated from the lower median (left) to the highest (right). S1: Thena5; S2: sanCAR1490; S3: sanCOST1250.5; S4: sanCOST1270.1; S5: sanOBAT1200; S6: sanOBAT1200.2; S7: sanRain39; S8: sanCAR1600.3; S9: Carv2015.1; S10: Carv2015.5; S11: Carv2015.11; S12: Carv2015.16; S13: Pico1680.1; S14: Pico1659.2; S15: Pico1659.3; S16: Amelia2015.1; S17: Amelia2015.6; S18: Amelia2015.12; S19: A1200.7; S20: Rain42.

**FIGURE S5.**
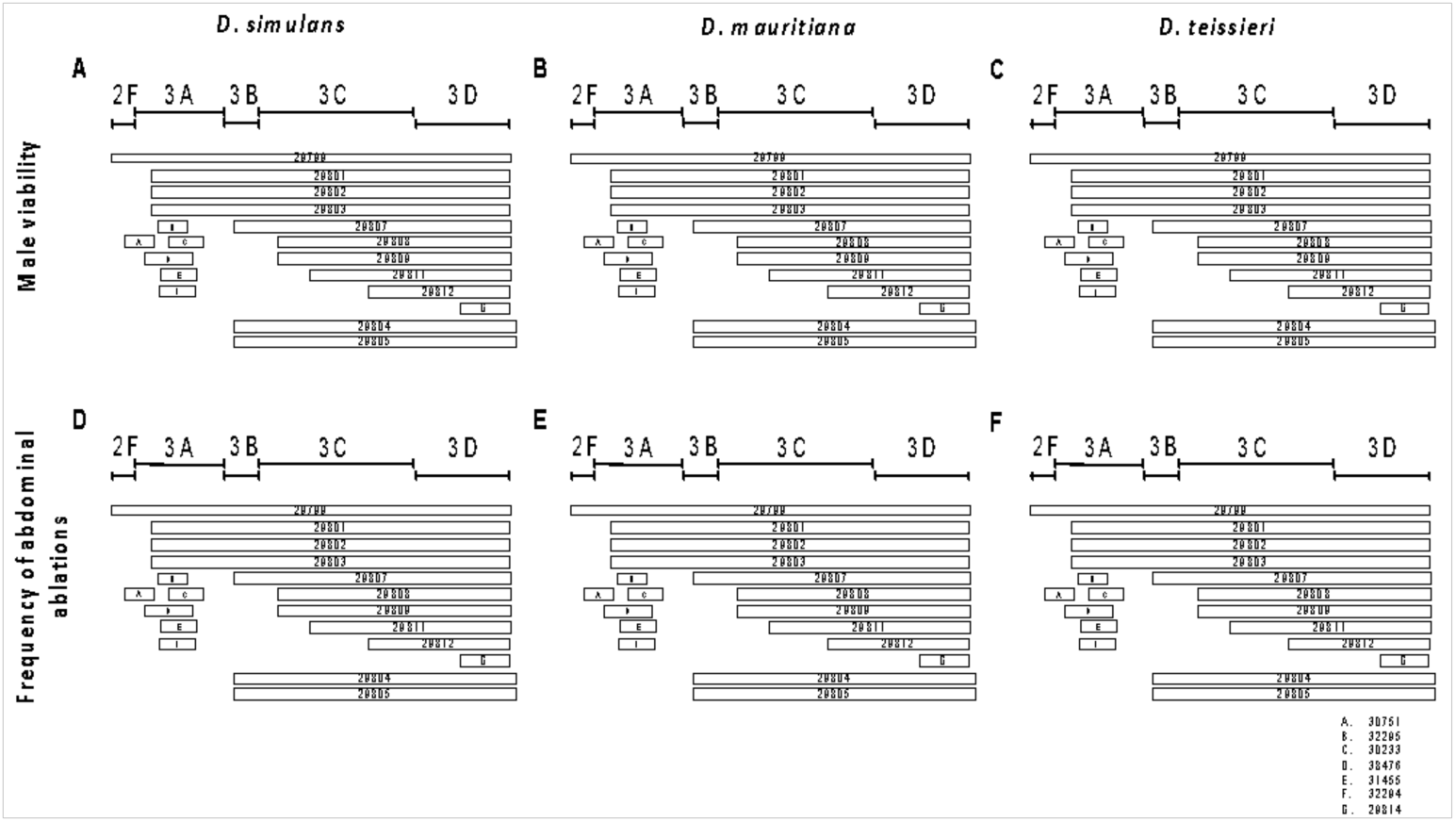
Crosses between *D. melanogaster* females harboring *Dp(1;Y)* duplications and males from the *simulans* species group (*D. mauritiana*, and *D. simulans*) show no evidence of male embryo lethality. *Dp(1:Y)* duplications containing *gt_mel_* cause no embryonic defects and do not cause heightened hybrid inviability. The color code is the same as used in Figure 2C. The lack of gray bars indicates that none of the duplications caused hybrid inviability in any of the crosses.

**FIGURE S6.**
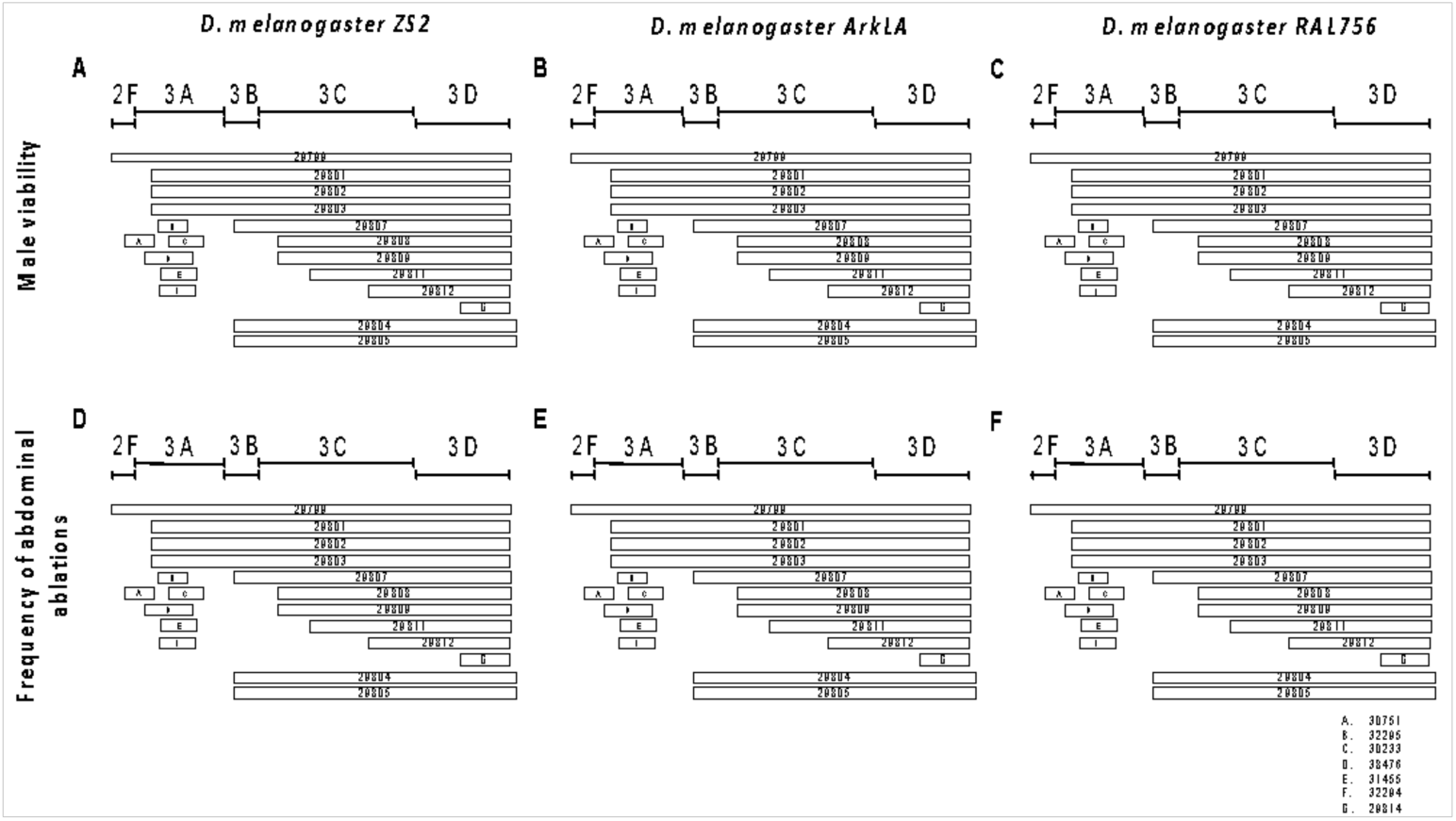
Crosses between *D. melanogaster* females harboring *Dp(1;Y)* duplications and males from different lines of *D. melanogaster* show no evidence of male embryo lethality or abdominal ablations. *Dp(1:Y)* duplications containing *gt^mel^* cause no embryonic defects and do not cause heightened HI. The color code is the same as used in Figure 2C. The lack of gray bars indicates that none of the duplications caused hybrid inviability in any of the crosses.

**FIGURE S7.**
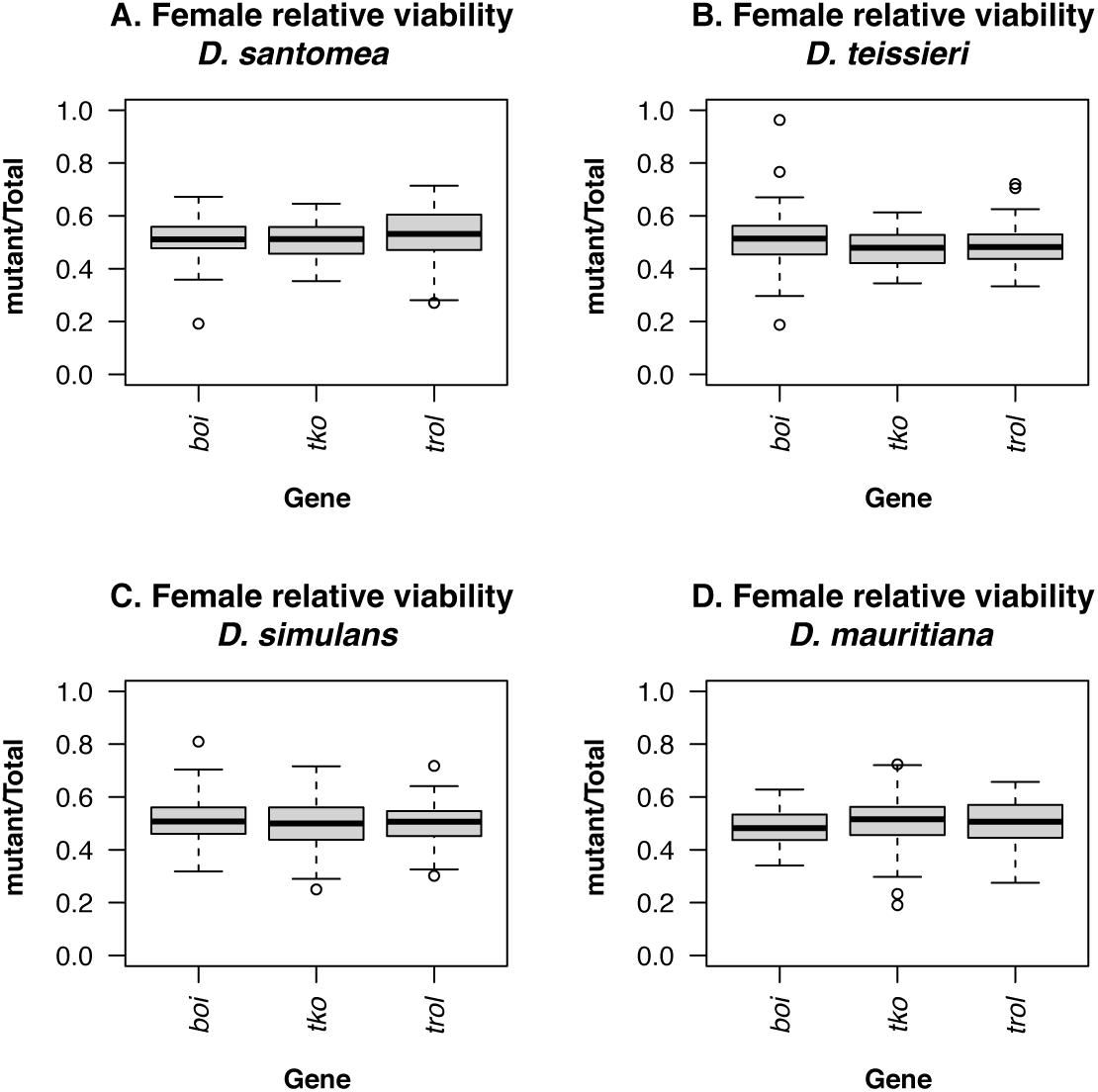
*mel/san* hybrid females carrying a *D. melanogaster* either a balancer chromosome or a mutant (hypomorphic or null) alleles for *boi*, *trol*, or *tko* show similar levels of hybrid viability. This is in contrast to males also carrying a *D. melanogaster* chromosome but a *gt* null allele (*gt_mel_^X11^*) which show a significant reduction in abdominal ablations (shown in Figure 2). These three genes have no effect in hybrid female viability between *D. melanogaster* and three more species (*D. teissieri* [B], *D. simulans* [C], and *D. mauritiana* [D]).

**FIGURE S8.**
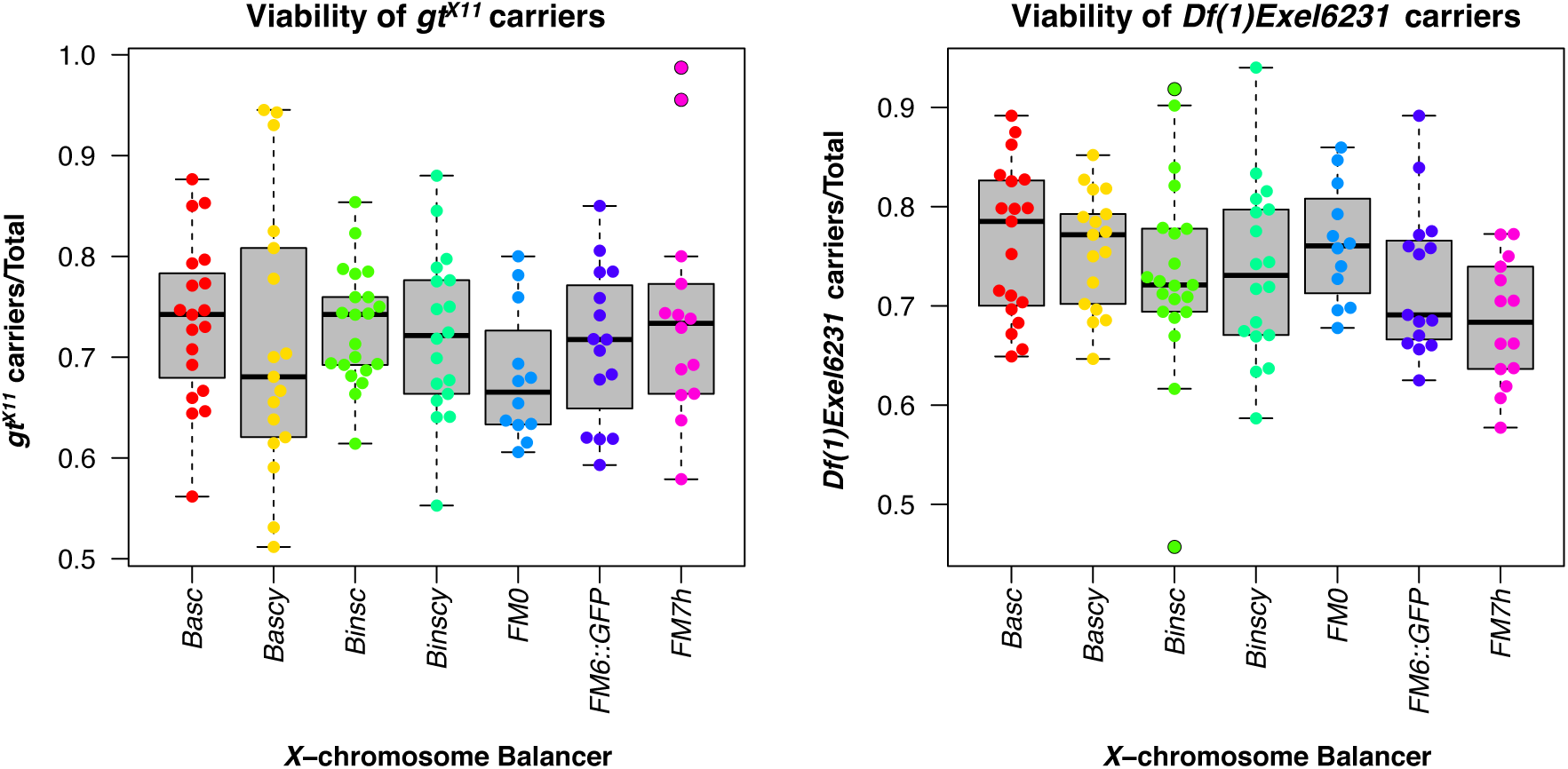
The *X*-chromosome balancer identity has no effect on the quantification of hybrid inviability in *mel/san* hybrid females. I found no heterogeneity in the relative viability of two different *gt* null mutants (A. *gt^X11^;* B. *Df(1)Exel6231*) when different balancer chromosomes are used (One-way ANOVA, F_6,109_= 0.694, P= 0.655). I used seven different *X*-chromosomes balancers and none of them had a major effect on the quantification of the relative frequency of viability in any of the hybrid crosses.

**FIGURE S9.**
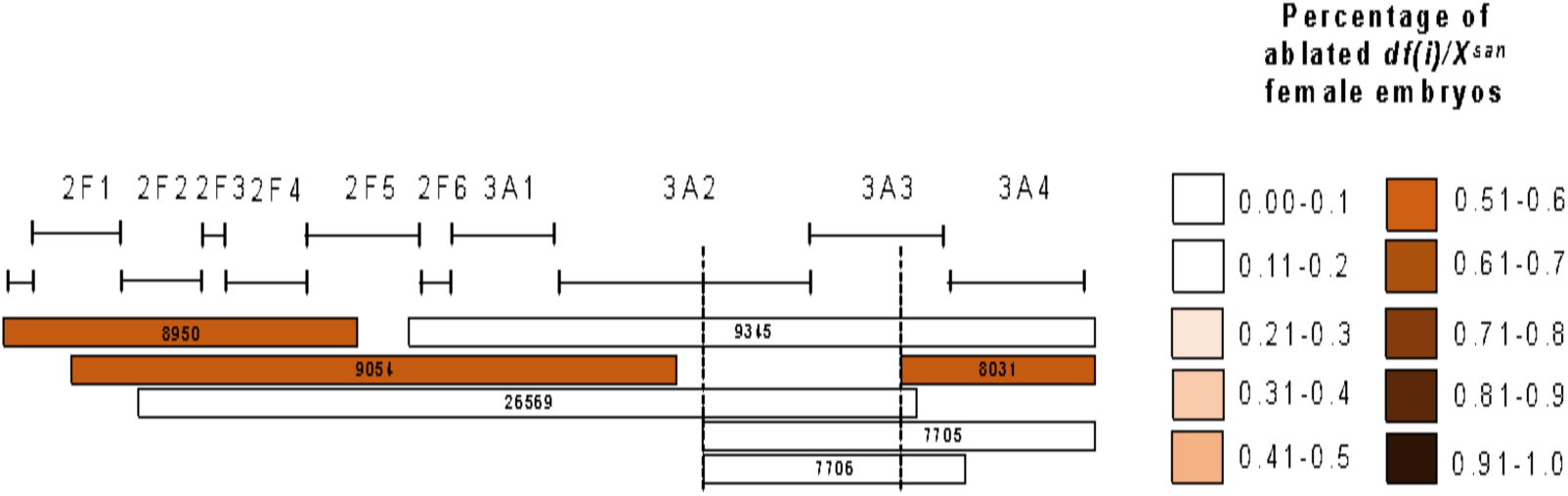
Frequency of abdominal ablations in each deficiency cross shown in Figure 3A. Deficiency chromosomes that contain a functional *gt_mel_* (e.g., 8031, 9054, and 8950) were more prone to show abdominal ablations than deficiencies that harbored other genes. Conversely, deficiency chromosomes with no functional copy of *gt_mel_* show reduced rates of abdominal ablations. The mean proportion of abdominal ablation in *X_mel_/X_san_* hybrid females is 0.420.

**FIGURE S10.**
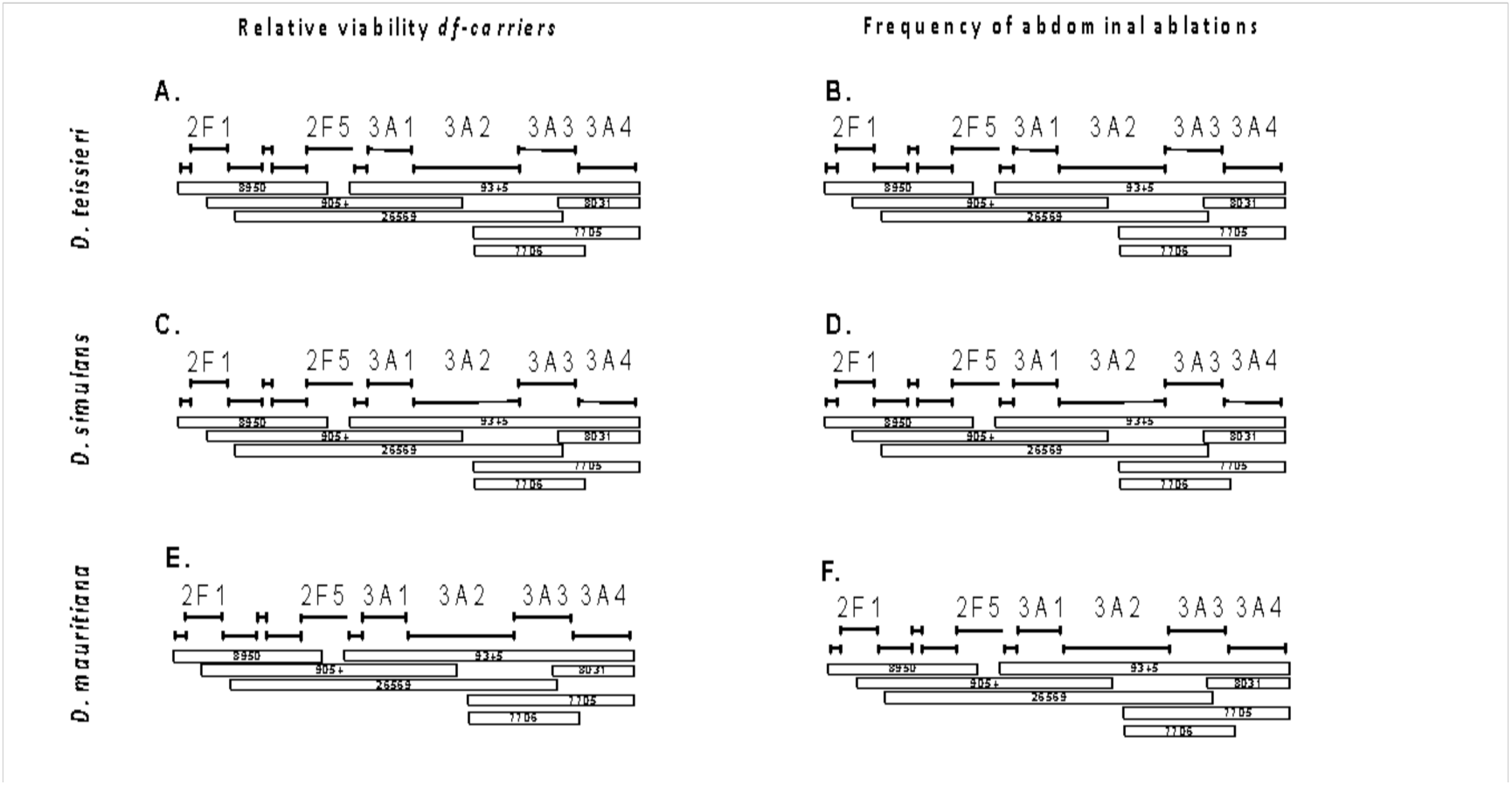
*gt^mel^*has no effect on the viability of *mel/tei*, *mel/mau*, and *mel/sim* hybrid females. I used the same deficiency chromosomes reported to detect the effect of *gt_mel_* in *mel/san* hybrid females to detect a potential effect of *gt_mel_* in hybrids between *D. melanogaster* and other species (Figure 3A). In all three cases, *df/FM7::GFP* crossed to males of each of the species led to 1:1 ratios in female progeny. The color code is the same as in Figure 3A but since no deficiency departed from the 1:1 expected (i.e., no *gt_mel_* effect on hybrid viability), there are no gray bars.

**FIGURE S11.**
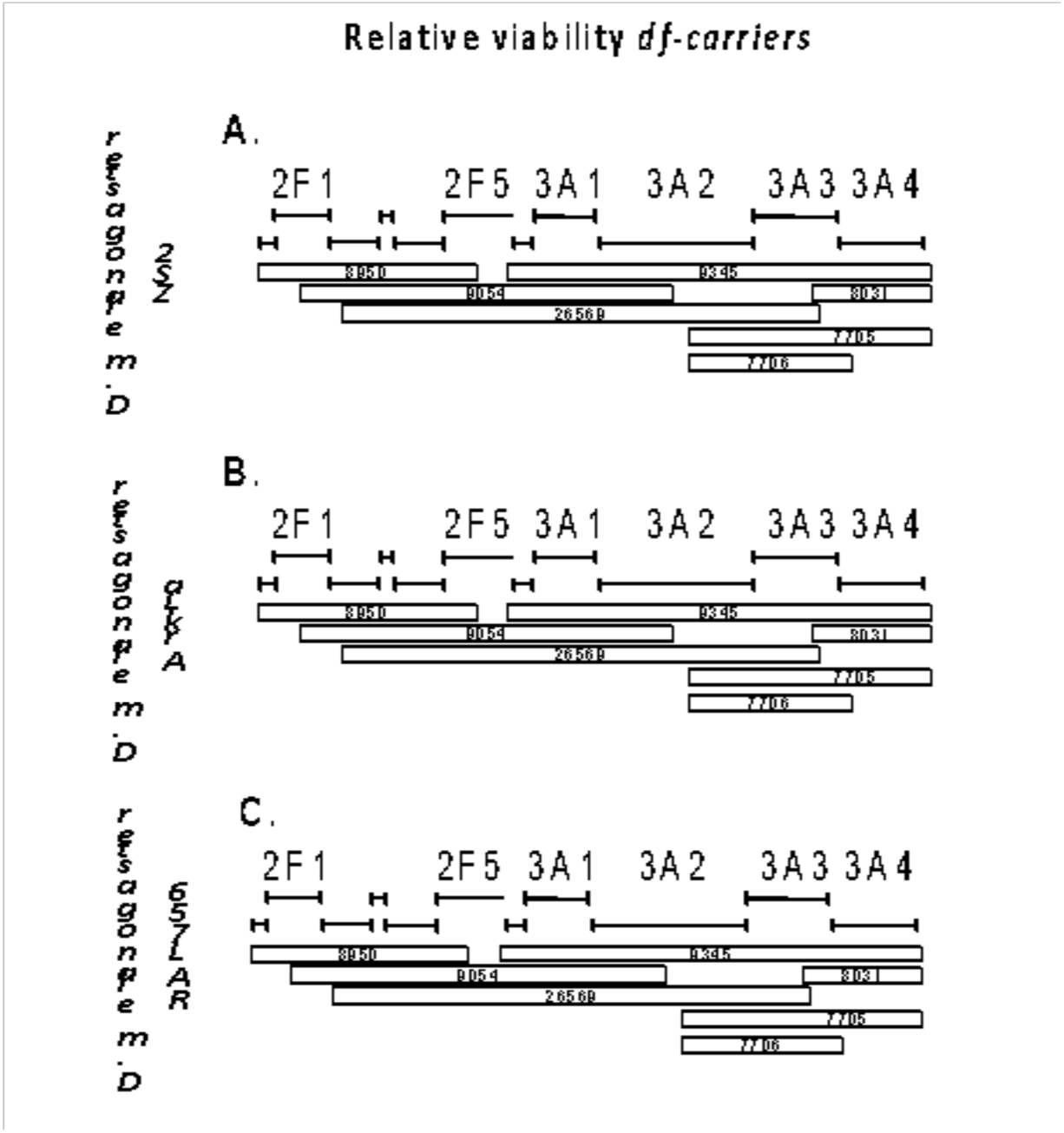
*gt_mel_* has no effect on the viability of pure-species *D. melanogaster* F1 females. I used the same deficiency chromosomes reported to detect the effect of *gt_mel_* in *mel/san* hybrid females (Figure 3A). In all three types of crosses (three isofemale lines), *df/FM7::GFP* crossed to males of each of the species led to 1:1 ratios in female progeny. The color code is the same as in Figure 3A but since no deficiency departed from the 1:1 expected (i.e., no *gt_mel_* effect on hybrid viability), there are no gray bars. No cross showed any abdominal ablation.

**FIGURE S12.**
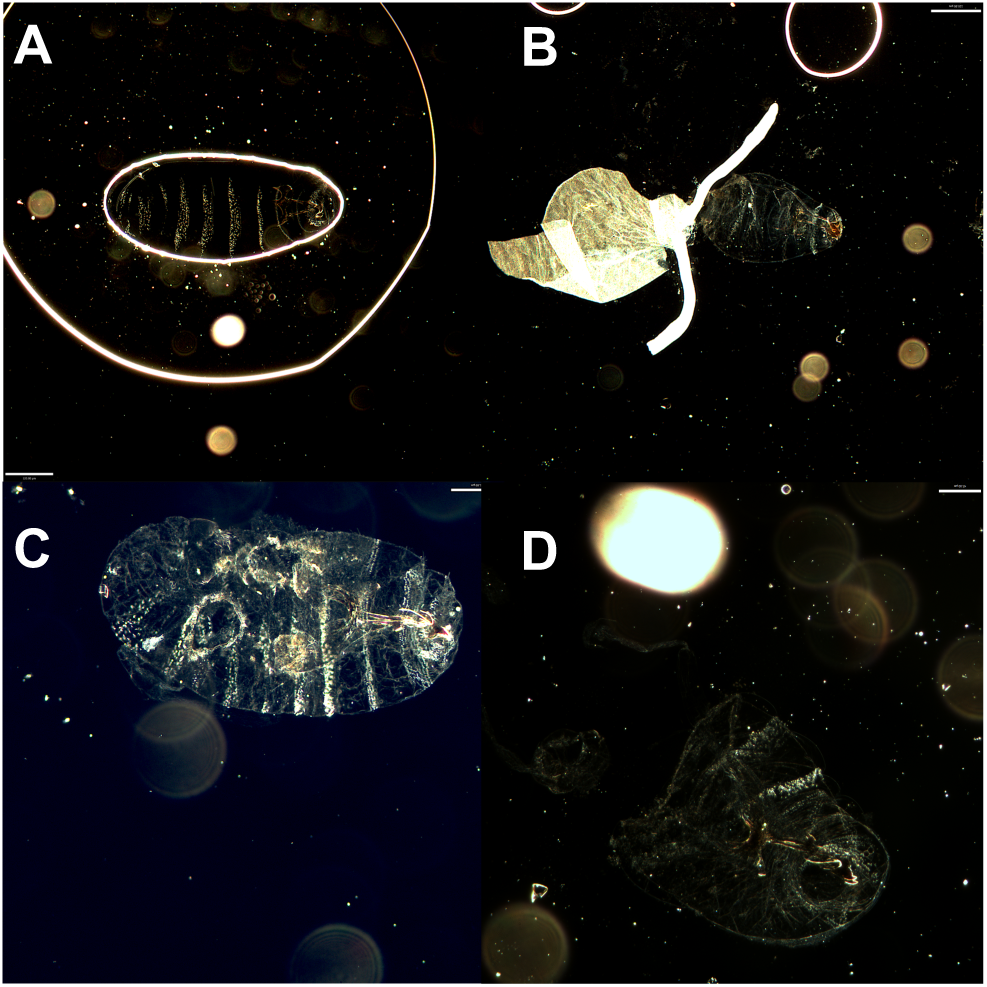
Hybrid male embryos carrying a *gt_mel_^X11^ D. melanogaster* allele show a variety of developmental defects. *gt_mel_^X11^/Y_san_ males* are inviable and show a variety of developmental defects (A-C). A small proportion of individuals also show abdominal ablations (D).

**FIGURE S13.**
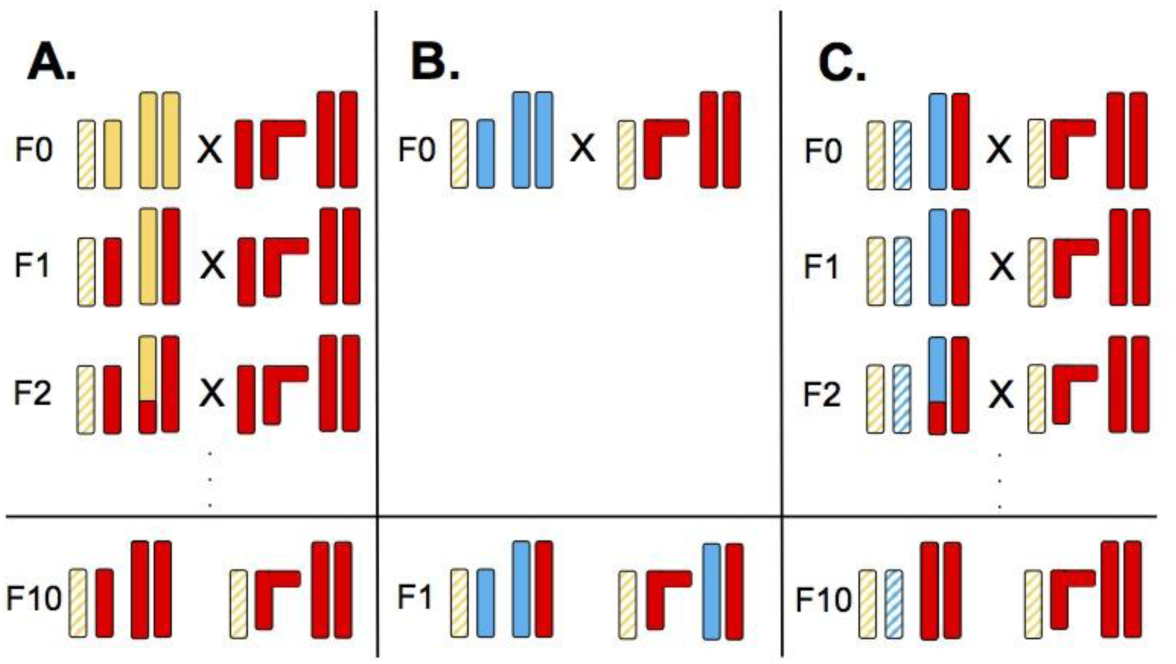
Introgression of the *FM7::GFP* and *gt^X11^* alleles into 200 lines of the DGRP lines. Each bar represents a chromosome. Short bars are the *X*-chromosome while longer bars represent the autosomes (only one set of autosomes shown). The bar with dashed lines represents the *FM7::GFP* balancer. Solid yellow represent the *FM7::GFP* background. Red bars represent each of the DGRP genetic backgrounds. Dashed blue lines represent the the *gt^X11^* chromosome, while solid blue lines represent the autosomal background of the *gt^X11^* stock. **A.** The first step of the experimental design involves introgressing the *FM7::GFP* balancer into each of the 200 DGRP backgrounds. After ten generations of repeating backcrossing, I obtained both females and males that carried the balancer and the DGRP background. **B.** Males from the cross shown in **A** (carriers of the *FM7::GFP* balancer) were crossed the *gt^X11^/FM7::GFP* females. This cross produces females that carry the *gt^X11^* chromosome, the *FM7::GFP* balancer and a mixed genetic background. These females were crossed to males that had the DGRP autosomal background and a *FM7::GFP* balancer (**C**). I repeated this backcrossing approach for ten generations. After, 11 generations I obtained *gt^X11^/FM7::GFP* females with DGRP autosomal backgrounds. These females were then crossed to *D. santomea* and the percentage of ablated progeny were scored.

**FIGURE S14.**
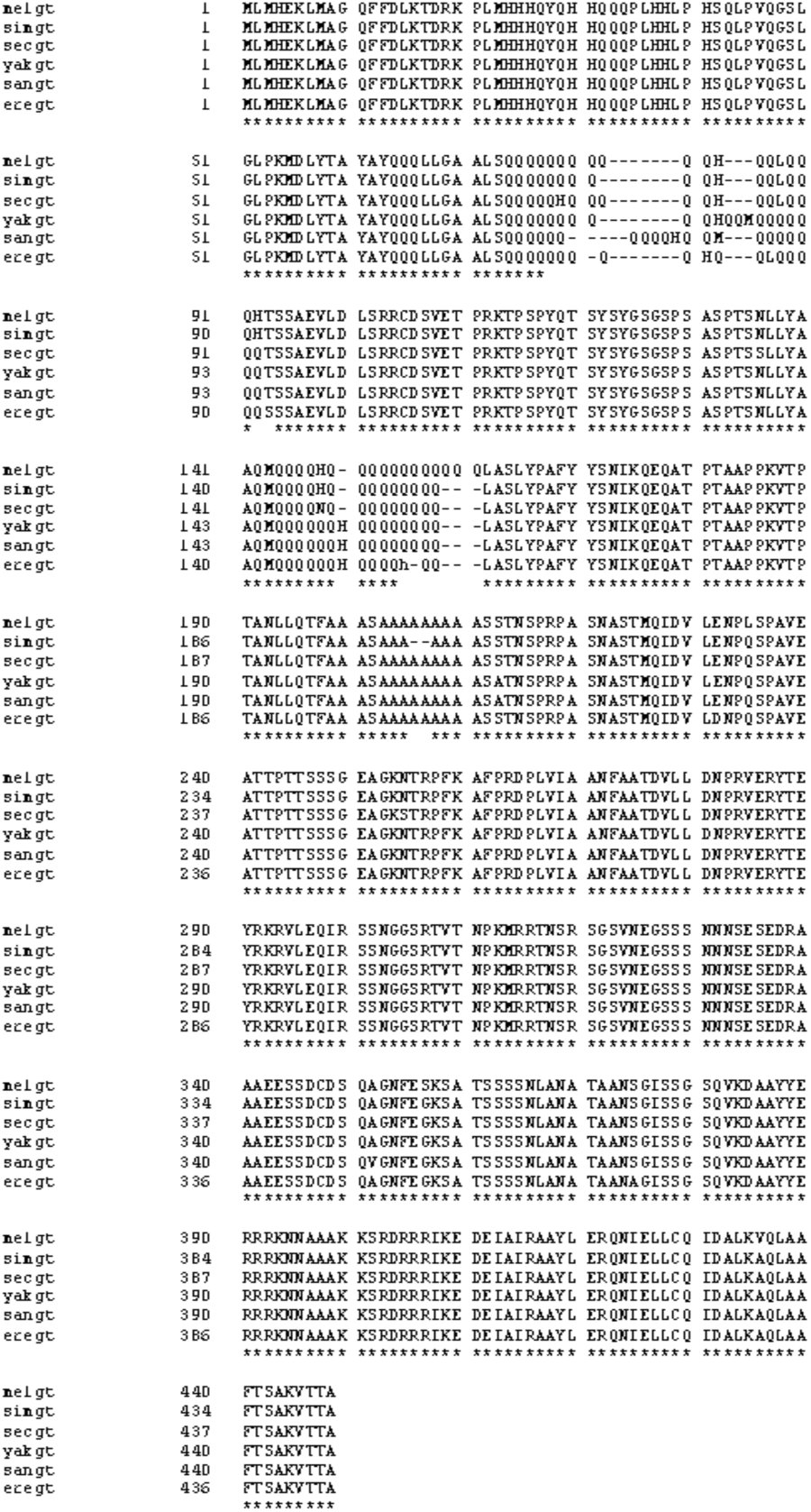
*gt* alleles from six different species in the *melanogaster* species complex. I found no major differences at the coding portion of *gt* between the *melanogaster* species supercomplex (*D. melanogaster*, *D. simulans*, and *D. sechellia*) and the *D. santomea/D. yakuba* species pair is the structure of poly-glutamine repeats. melgt: *D. melanogaster*, simgt: *D. simulans*, sechgt: *D. sechellia*, sangt: *D. santomea*, eregt: *D. erecta*. Asterisks show residues that are conserved across the whole group.

**FIGURE S15.**
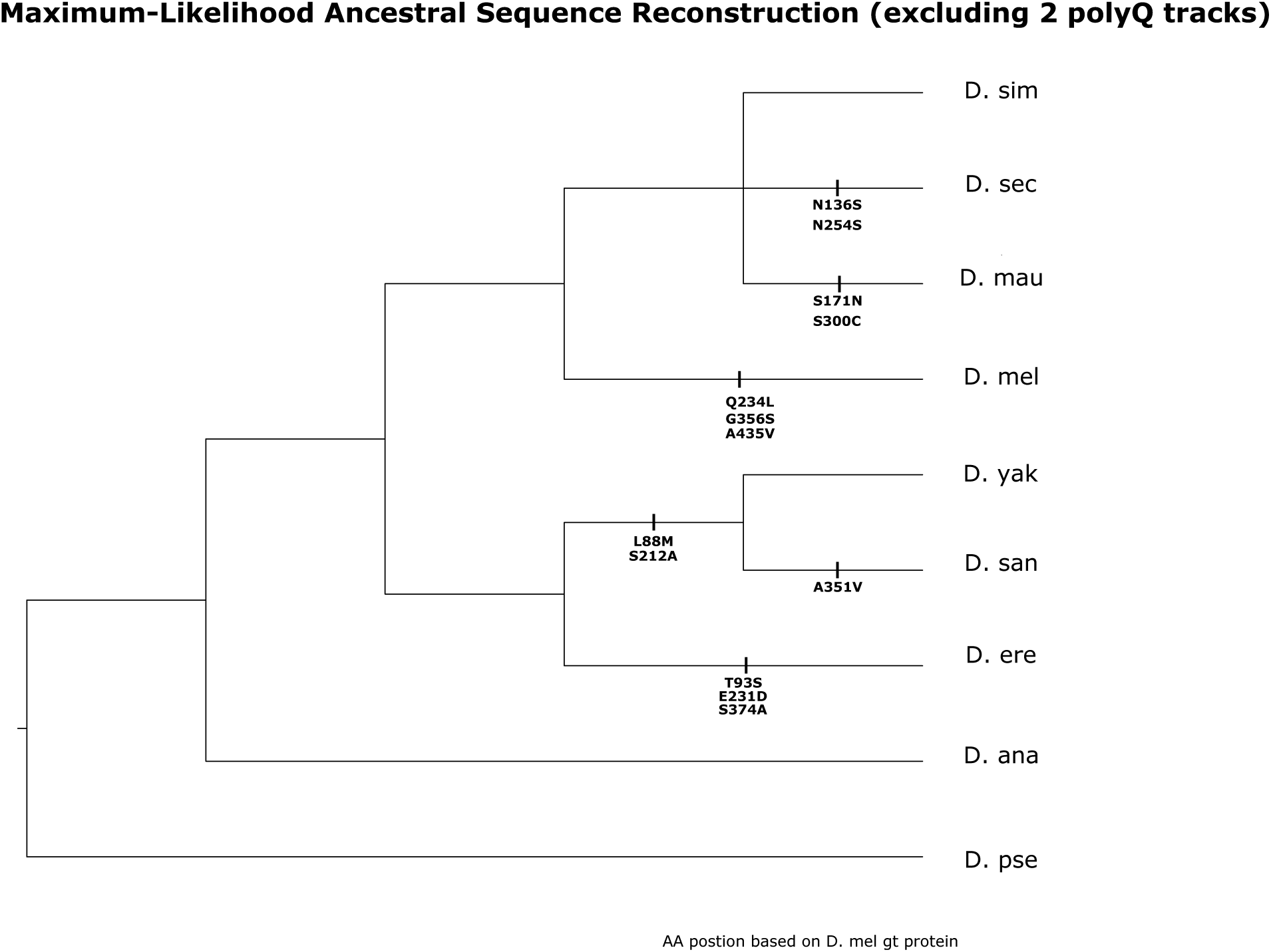
Maximum-likelihood ancestral sequence reconstruction of GT protein, excluding polyQ. All ancestral sites could be reconstructed with high confidence (posterior probability > 0.98), except for the two polyQ tracks. Aminoacid positions are all based on the *D. melanogaster GT* protein.

**FIGURE S16.**
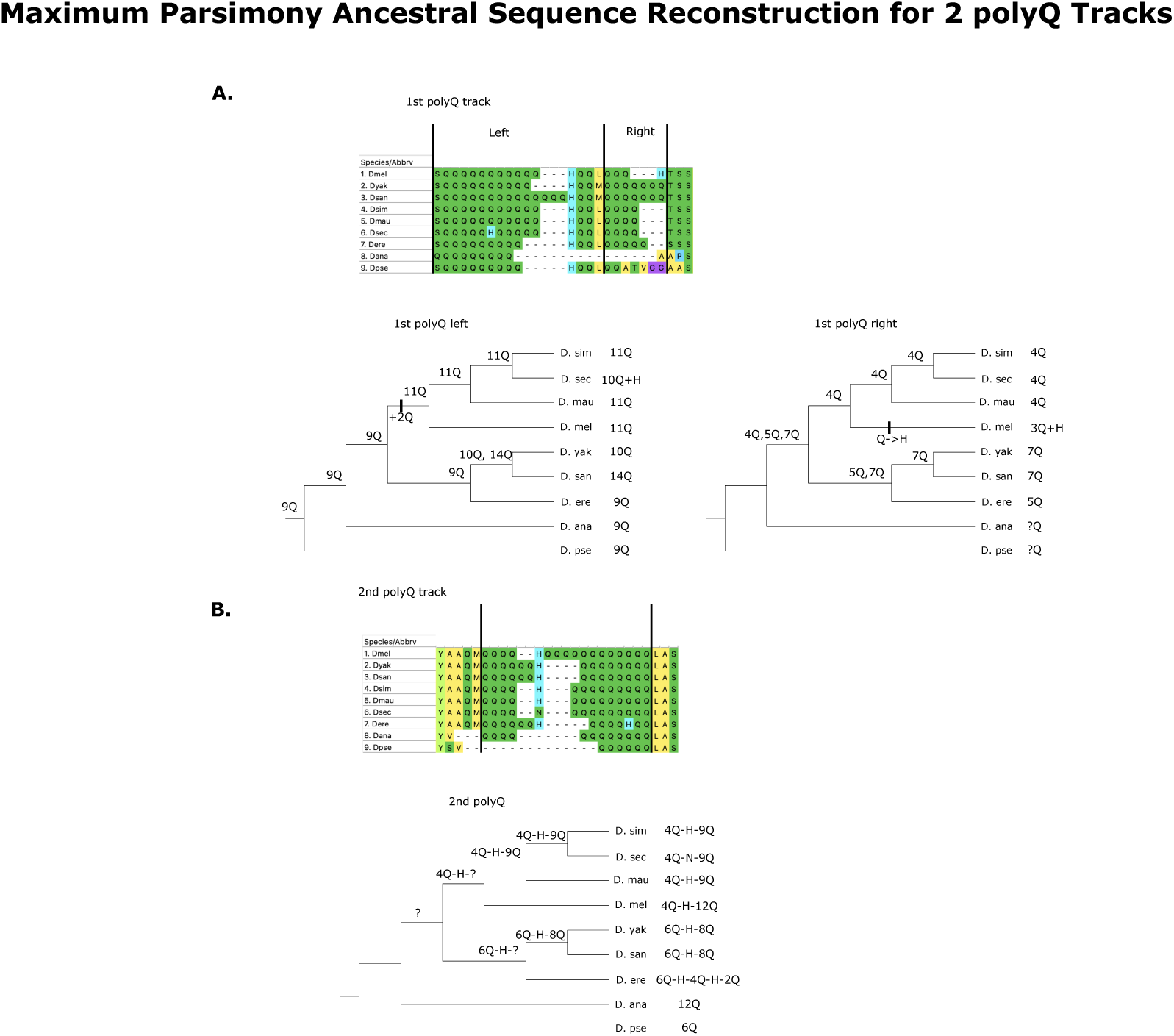
The polyglutamine repeats (polyQ) show differences among the *melanogaster* subspecies complex species. Maximum-parsimony ancestral sequence reconstruction of 2 polyQ tracks of gt protein. The conserved H within polyQ tracks helps to delineate both polyQ tracks to two parts. Due to the lack of proper substitution models, this maximum-parsimony based ancestral reconstruction for polyQ tracks may subject to bias due to alignment error, and arbitrary choice of polyQ unit.

**FIGURE S17.**
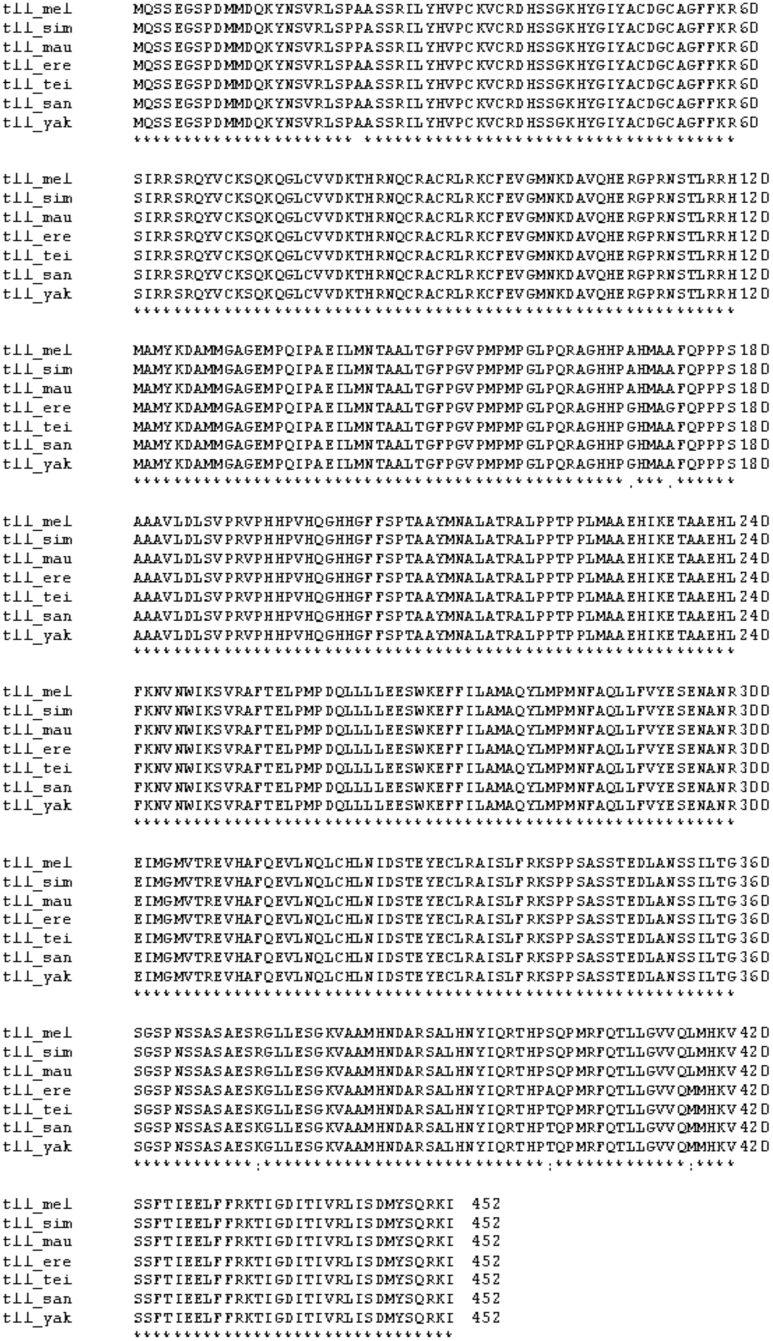
*tll* alleles from six different species in the *melanogaster* species complex. I found no major differences at the coding portion of *tll* between the *melanogaster* species supercomplex (*D. melanogaster*, *D. simulans*, and *D. sechellia*) and the *D. santomea/D. yakuba* species pair. tll_mel: *D. melanogaster*, tll_sim: *D. simulans*, tll_mau: *D. mauritiana*, tll_ere: *D. erecta*, tll_tei: *D. teissieri,* tll_san: *D. santomea*, tll_yak: *D. yakuba*. Asterisks show residues that are conserved across the whole group.

**FIGURE S18.**
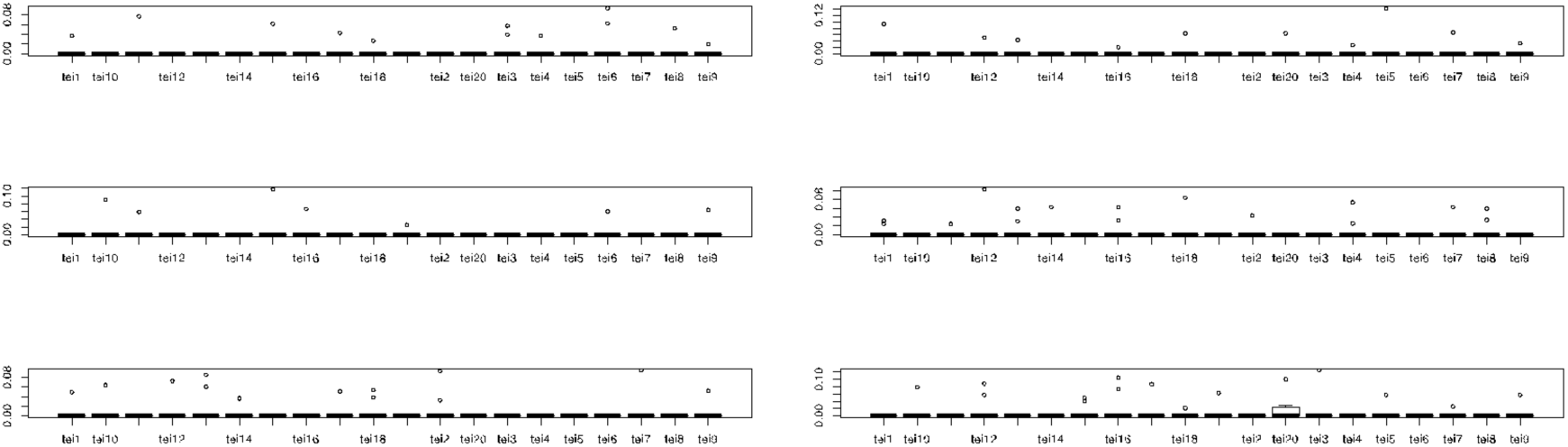
Embryonic hybrid inviability does not occur in *mel/tei* hybrids. No line of *D. teissieri* showed either embryonic inviability or abdominal ablations when crossed with *D. melanogaster*, *D. simulans*, or *D. mauritiana* females. The vast majority of assays revealed no dead embryos. **A.** *mel/tei* male hybrids. **B.** *sim/tei* male hybrids. **C.** *mau/tei* male hybrids. **D.** *mel/tei* female hybrids. **E.** *sim/tei* female hybrids. **F.** *mau/tei* female hybrids. tei1: Balancha_1; tei2: Balancha_2, tei3: Balancha_3, tei4: House_Bioko_0, tei5: House_Bioko_1, tei6: House_Bioko_2, tei7: La_Lope_Gabon, tei8: Selinda, tei9: Zimbabwe, tei10: cascade_2_1, tei11: cascade_2_2; tei12: cascade_2_4, tei13: cascade_4_1, tei14: cascade_4_2, tei15: cascade_4_3, tei16: cascade_4_4, tei17: cascade_4_5, tei18: cascade_4_6, tei19: Bata_2, tei20: Bata_8.

**FIGURE S19.**
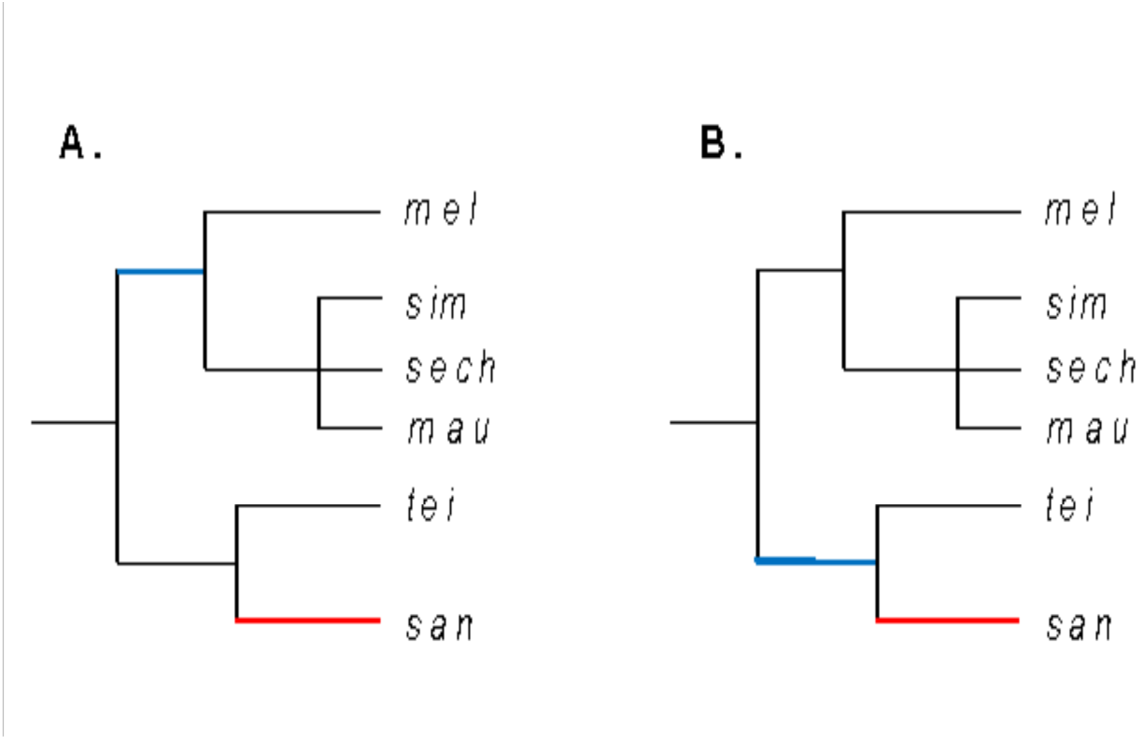
Divergence in *giant* in the *melanogaster* species subgroup. **A**. The evolutionary timing of *gt^mel^* and its interactor leading to HI. The *gt* allele responsible for HI in hybrids with *D. santomea* evolved before *mel, sim, and mau* had a common ancestor (Blue branch). At least one of the interactors of *gt^mel^* is not shared with *D. teissieri* which indicates that such element must have evolved after the last common ancestor of *D. santomea* and *D. teissieri* speciated (Red branch).

**FIGURE S20.**
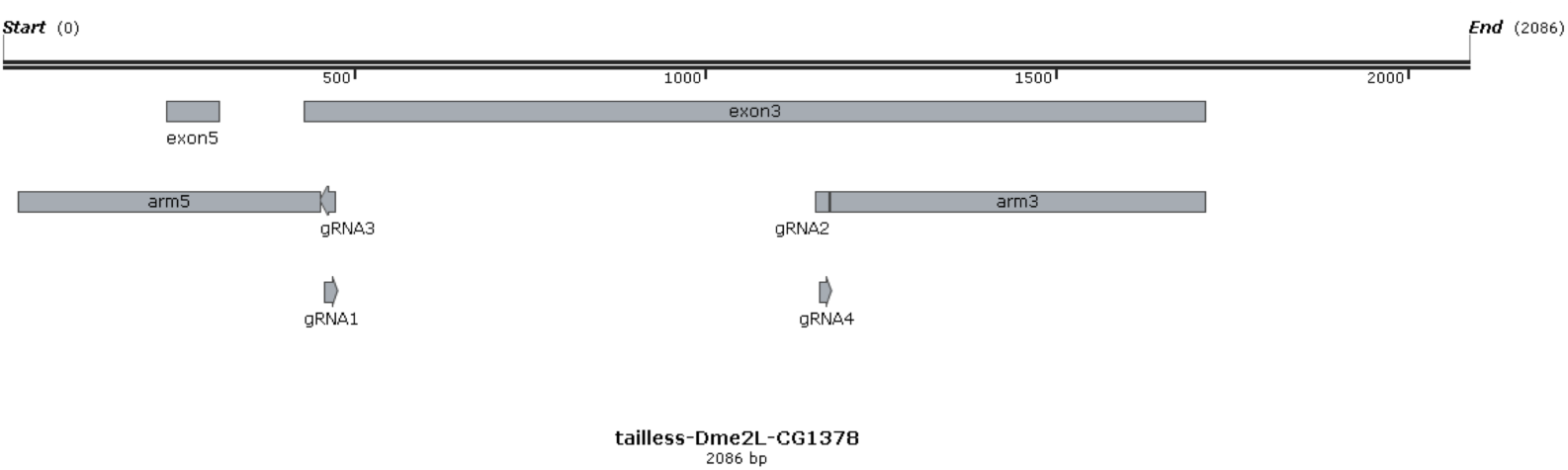
Experimental design to generated a GFP-mediated disruption of *tll_mel_*.

## REFERENCES

1. J. A. Coyne, H. A. Orr, Speciation, Sunderland, MA (Sinauer Associates, 2004).

2. J. M. Coughlan, D. R. Matute, The importance of intrinsic postzygotic barriers throughout the speciation process. Philosophical Transactions of the Royal Society B: Biological Sciences 375, 20190533 (2020).

3. C. Darwin, The origin of species by means of natural selection (Pub One Info, 1859).

4. J. J. Weir, An Unnoticed Factor in Evolution. Nature 31, 194–194 (1885).

5. G. H. Shull, The Species Concept from the Point of View of a Geneticist. American Journal of Botany 10, 221–228 (1923).

6. T. Dobzhansky, Genetic nature of species differences. The American Naturalist 71, 404–420 (1937).

7. H. J. Muller, Isolating mechanisms, evolution and temperature (1942).

8. S. Maheshwari, D. A. Barbash, The Genetics of Hybrid Incompatibilities. Annual Review of Genetics 45, 331–355 (2011).

9. C. Wellbrock, et al., Signalling by the oncogenic receptor tyrosine kinase Xmrk leads to activation of STAT5 in Xiphophorus melanoma. Oncogene 16, 3047–3056 (1998).

10. D. C. Presgraves, A fine-scale genetic analysis of hybrid incompatibilities in *Drosophila*. Genetics 163, 955–972 (2003).

11. N. J. Brideau, et al., Two Dobzhansky-Muller Genes interact to cause hybrid lethality in Drosophila. Science 314, 1292–1295 (2006).

12. K. Bomblies, et al., Autoimmune response as a mechanism for a Dobzhansky-Muller-type incompatibility syndrome in plants. PLoS Biology (2007). 10.1371/journal.pbio.0050236.

13. H. Y. Lee, et al., Incompatibility of Nuclear and Mitochondrial Genomes Causes Hybrid Sterility between Two Yeast Species. Cell 135, 1065–1073 (2008).

14. S. Tang, D. C. Presgraves, Evolution of the Drosophila nuclear pore complex results in multiple hybrid incompatibilities. Science 323, 779–782 (2009).

15. P. M. Ferree, D. A. Barbash, Species-Specific Heterochromatin Prevents Mitotic Chromosome Segregation to Cause Hybrid Lethality in Drosophila. PLOS Biology 7, e1000234 (2009).

16. N. Phadnis, et al., An essential cell cycle regulation gene causes hybrid inviability in Drosophila. Science (New York, N.Y.) 350, 1552–5 (2015).

17. M. P. Zuellig, A. L. Sweigart, Gene duplicates cause hybrid lethality between sympatric species of Mimulus. PLOS Genetics 14, e1007130 (2018).

18. D. L. Powell, et al., Natural hybridization reveals incompatible alleles that cause melanoma in swordtail fish. Science 368, 731–736 (2020).

19. B. M. Moran, et al., “A Lethal Genetic Incompatibility between Naturally Hybridizing Species in Mitochondrial Complex I” (2021).

20. P. R. V. Satyaki, et al., The Hmr and Lhr Hybrid Incompatibility Genes Suppress a Broad Range of Heterochromatic Repeats. PLoS Genetics 10 (2014).

21. N. Phadnis, H. A. Orr, A single gene causes both male sterility and segregation distortion in Drosophila hybrids. Science (2009). 10.1126/science.1163934.

22. N. A. Johnson, Hybrid incompatibility genes: Remnants of a genomic battlefield? (2010).

23. K. L. Gordon, I. Ruvinsky, Tempo and Mode in Evolution of Transcriptional Regulation. PLOS Genetics 8, e1002432 (2012).

24. M. Manzanares, et al., Conservation and elaboration of Hox gene regulation during evolution of the vertebrate head. Nature 408, 854–857 (2000).

25. S. J. Gaunt, Conservation in the Hox code during morphological evolution. International Journal of Developmental Biology 38, 549–552 (2002).

26. S. Santini, J. L. Boore, A. Meyer, Evolutionary conservation of regulatory elements in vertebrate Hox gene clusters. Genome research 13, 1111–1122 (2003).

27. A. P. Lee, E. G. Koh, A. Tay, S. Brenner, B. Venkatesh, Highly conserved syntenic blocks at the vertebrate Hox loci and conserved regulatory elements within and outside Hox gene clusters. Proceedings of the National Academy of Sciences 103, 6994–6999 (2006).

28. E. E. Hare, B. K. Peterson, V. N. Iyer, R. Meier, M. B. Eisen, Sepsid even-skipped enhancers are functionally conserved in Drosophila despite lack of sequence conservation. PLoS Genetics 4 (2008).

29. K. M. Weiss, S. M. Fullerton, Phenogenetic drift and the evolution of genotype-phenotype relationships (2000).

30. J. R. True, E. S. Haag, Developmental system drift and flexibility in evolutionary trajectories (2001).

31. M. E. Palmer, M. W. Feldman, Dynamics of hybrid incompatibility in gene networks in a constant environment. Evolution 63, 418–431 (2009).

32. M. Pavlicev, G. P. Wagner, A model of developmental evolution: Selection, pleiotropy and compensation (2012).

33. M. Rebeiz, et al., Evolution of the tan Locus Contributed to Pigment Loss in Drosophila santomea: A Response to Matute et al. Cell 139, 1189–1196 (2009).

34. J. D. Kinsey, Studies on an embryonic lethal hybrid in Drosophila. (1967).

35. K. Sawamura, M. T. Yamamoto, Cytogenetical localization of Zygotic hybrid rescue (Zhr), a Drosophila melanogaster gene that rescues interspecific hybrids from embryonic lethality. MGG Molecular & General Genetics (1993). 10.1007/BF00276943.

36. K. Sawamura, M. T. Yamamoto, T. K. Watanabe, Hybrid lethal systems in the Drosophila melanogaster species complex. II. The Zygotic hybrid rescue (Zhr) gene of D. melanogaster. Genetics 133, 307–313 (1993).

37. K. Sawamura, T. Taira, T. K. Watanabe, Hybrid lethal systems in the Drosophila melanogaster species complex. I. The maternal hybrid rescue (mhr) gene of Drosophila simulans. Genetics 133, 299–305 (1993).

38. J. Gavin-Smyth, D. R. Matute, Embryonic lethality leads to hybrid male inviability in hybrids between Drosophila melanogaster and D. santomea. Ecology and Evolution 3, 1580–1589 (2013).

39. M. Z. Ludwig, et al., Functional evolution of a cis-regulatory module. PLoS Biology 3, 0588–0598 (2005).

40. X. Wang, R. J. Sommer, Antagonism of LIN-17/Frizzled and LIN-18/Ryk in Nematode Vulva Induction Reveals Evolutionary Alterations in Core Developmental Pathways. PLOS Biology 9, e1001110 (2011).

41. R. J. Sommer, “Evolution of Regulatory Networks: Nematode Vulva Induction as an Example of Developmental Systems Drift” in Evolutionary Systems Biology, Advances in Experimental Medicine and Biology., O. S. Soyer, Ed. (Springer, 2012), pp. 79–91.

42. C. M. S. Cauret, et al., Developmental Systems Drift and the Drivers of Sex Chromosome Evolution. Molecular Biology and Evolution 37, 799–810 (2020).

43. V. J. Lynch, Use with caution: Developmental systems divergence and potential pitfalls of animal models. Yale J Biol Med 82, 53–66 (2009).

44. D. R. Matute, I. A. Butler, D. A. Turissini, J. A. Coyne, A test of the snowball theory for the rate of evolution of hybrid incompatibilities. Science 329, 1518–1521 (2010).

45. A. Suvorov, et al., Widespread introgression across a phylogeny of 155 Drosophila genomes. bioRxiv 2020.12.14.422758 (2020). 10.1101/2020.12.14.422758.

46. K. Tamura, S. Subramanian, S. Kumar, Temporal Patterns of Fruit Fly (Drosophila) Evolution Revealed by Mutation Clocks. Molecular Biology and Evolution 21, 36–44 (2004).

47. D. R. Matute, J. Gavin-Smyth, Fine Mapping of Dominant X-Linked Incompatibility Alleles in Drosophila Hybrids. PLoS Genetics 10 (2014).

48. A. Suvorov, et al., Widespread introgression across a phylogeny of 155 Drosophila genomes. Current Biology 32, 111–123.e5 (2022).

49. D. A. Turissini, G. Liu, J. R. David, D. R. Matute, The evolution of reproductive isolation in the Drosophila yakuba complex of species. J. Evol. Biol. 28, 557–575 (2015).

50. D. A. Turissini, J. A. McGirr, S. S. Patel, J. R. David, D. R. Matute, The rate of evolution of postmating-prezygotic reproductive isolation in drosophila. Molecular Biology and Evolution 35, 312–334 (2018).

51. B. S. Cooper, P. S. Ginsberg, M. Turelli, D. R. Matute, Wolbachia in the Drosophila yakuba complex: Pervasive frequency variation and weak cytoplasmic incompatibility, but no apparent effect on reproductive isolation. Genetics 205, 333–351 (2017).

52. A. H. Sturtevant, Genetic Studies on Drosophila Simulans. I. Introduction. Hybrids With Drosophila Imelanogaster. Genetics 5, 488–500 (1920).

53. D. R. Matute, J. Gavin-Smyth, G. Liu, Variable post-zygotic isolation in Drosophila melanogaster/D. simulans hybrids. Journal of Evolutionary Biology (2014). 10.1111/jeb.12422.

54. T. F. C. MacKay, et al., The Drosophila melanogaster Genetic Reference Panel. Nature 482, 173–178 (2012).

55. W. Huang, et al., Natural variation in genome architecture among 205 Drosophila melanogaster Genetic Reference Panel lines. Genome Research 24, 1193–1208 (2014).

56. R. K. Cook, et al., A new resource for characterizing X-linked genes in Drosophila melanogaster: Systematic coverage and subdivision of the *X* chromosome with nested, Y-linked duplications. Genetics 186, 1095–1109 (2010).

57. K. J. T. Venken, et al., A molecularly defined duplication set for the X chromosome of *Drosophila melanogaster*. Genetics 186, 1111–1125 (2010).

58. G. Dos Santos, et al., FlyBase: Introduction of the Drosophila melanogaster Release 6 reference genome assembly and large-scale migration of genome annotations. Nucleic Acids Research 43, D690–D697 (2015).

59. N. Perrimon, L. Engstrom, A. P. Mahowald, Developmental Genetics of the 2e-F Region of the Drosophila X Chromosome: A Region Rich in “Developmentally Important” Genes. Genetics 108, 559–572 (1984).

60. J. P. Petschek, N. Perrimon, A. P. Mahowald, Region-specific defects in l(1)giant embryos of *Drosophila melanogaster*. Developmental Biology 119, 175–189 (1987).

61. J. A. Coyne, S. Simeonidis, P. Rooney, Relative paucity of genes causing inviability in hybrids between Drosophila melanogaster and D. simulans. Genetics 150, 1091–1103 (1998).

62. J. Kaminker, et al., The transposable elements of the *Drosophila melanogaster* euchromatin: a genomics perspective. Genome Biology 3, 1–20 (2002).

63. J. B. Brown, et al., Diversity and dynamics of the Drosophila transcriptome. Nature 512, 393–399 (2014).

64. P. Nosil, D. Schluter, The genes underlying the process of speciation (2011).

65. J. Mohler, E. D. Eldon, V. Pirrotta, A novel spatial transcription pattern associated with the segmentation gene, giant, of Drosophila. The EMBO journal 8, 1539–1548 (1989).

66. E. D. Eldon, V. Pirrotta, Interactions of the Drosophila gap gene giant with maternal and zygotic pattern-forming genes. Development (Cambridge, England) 111, 367–378 (1991).

67. R. Kraut, M. Levine, Mutually repressive interactions between the gap genes giant and Krüppel define middle body regions of the Drosophila embryo. Development (Cambridge, England) 111, 611–621 (1991).

68. M. Capovilla, E. D. Eldon, V. Pirrotta, The giant gene of Drosophila encodes a b-ZIP DNA-binding protein that regulates the expression of other segmentation gap genes. Development (Cambridge, England) 114, 99–112 (1992).

69. X. Wu, R. Vakani, S. Small, Two distinct mechanisms for differential positioning of gene expression borders involving the *Drosophila* gap protein giant. Development 125, 3765–3774 (1998).

70. G. Struhl, P. Johnston, P. A. Lawrence, Control of Drosophila body pattern by the hunchback morphogen gradient. Cell 69, 237–249 (1992).

71. J. Reinitz, M. Levine, Control of the initiation of homeotic gene expression by the gap genes giant and tailless in *Drosophila*. Developmental Biology 140, 57–72 (1990).

72. G. Brönner, et al., Sp1/egr-like zinc-finger protein required for endoderm specification and germ-layer formation in *Drosophila*. Nature 369, 664–668 (1994).

73. C. Schulz, D. Tautz, Zygotic caudal regulation by hunchback and its role in abdominal segment formation of the *Drosophila* embryo. Development (Cambridge, England) 121, 1023–1028 (1995).

74. Rivera-Pomar R, Lu X, Perrimon N, Taubert H, Jäckle H. Activation of posterior gap gene expression in the Drosophila blastoderm. Nature. 1995 Jul 20;376(6537):253–6.

75. J. Jaeger, The gap gene network (2011).

76. E. E. Hare, B. K. Peterson, M. B. Eisen, A careful look at binding site reorganization in the even-skipped enhancers of Drosophila and sepsids (2008).

77. G. Bucher, Divergent segmentation mechanism in the short germ insect Tribolium revealed by *giant* expression and function. Development 131, 1729–1740 (2004).

78. M. J. Wilson, M. Havler, P. K. Dearden, Giant, Krüppel, and caudal act as gap genes with extensive roles in patterning the honeybee embryo. Developmental Biology 339, 200–211 (2010).

79. K. R. Nitta, et al., Conservation of transcription factor binding specificities across 600 million years of bilateria evolution. eLife 4, e04837 (2015).

80. Y. Goltsev, W. Hsiong, G. Lanzaro, M. Levine, Different combinations of gap repressors for common stripes in Anopheles and Drosophila embryos. Developmental Biology 275, 435–446 (2004).

81. A. Crombach, M. A. García-Solache, J. Jaeger, Evolution of early development in dipterans: Reverse-engineering the gap gene network in the moth midge *Clogmia albipunctata* (Psychodidae). BioSystems 123, 74–85 (2014).

82. A. Crombach, K. R. Wotton, E. Jiménez-Guri, J. Jaeger, Gap Gene Regulatory Dynamics Evolve along a Genotype Network. Molecular Biology and Evolution 33, 1293–1307 (2016).

83. K. R. Wotton, et al., Quantitative system drift compensates for altered maternal inputs to the gap gene network of the scuttle fly *Megaselia abdita*. eLife 2015 (2015).

84. M. García-Solache, J. Jaeger, M. Akam, A systematic analysis of the gap gene system in the moth midge *Clogmia albipunctata*. Developmental Biology 344, 306–318 (2010).

85. H. Janssens, et al., A quantitative atlas of *even-skipped* and *Hunchback* expression in Clogmia albipunctata (Diptera: Psychodidae) blastoderm embryos. EvoDevo 5 (2014).

86. A. C. Cerny, D. Grossmann, G. Bucher, M. Klingler, The Tribolium ortholog of knirps and knirps-related is crucial for head segmentation but plays a minor role during abdominal patterning. Developmental Biology 321, 284–294 (2008).

87. J. S. Schiffman, P. L. Ralph, System drift and speciation. Evolution (2021). 10.1111/evo.14356.

88. B. Z. He, A. K. Holloway, S. J. Maerkl, M. Kreitman, Does positive selection drive transcription factor binding site turnover? a test with drosophila *cis*-regulatory modules. PLoS Genetics 7 (2011).

89. X. Ni, et al., Adaptive Evolution and the Birth of CTCF Binding Sites in the Drosophila Genome. PLoS Biology 10 (2012).

90. D. Bachtrog, K. Thornton, A. Clark, P. Andolfatto, Extensive introgression of mitochondrial DNA relative to nuclear genes in the *Drosophila yakuba* species group. Evolution 60, 292–302 (2006).

91. D. A. Turissini, D. R. Matute, Fine scale mapping of genomic introgressions within the *Drosophila yakuba* clade. PLOS Genetics 13, e1006971 (2017).

92. L. C. Moyle, T. Nakazato, Hybrid incompatibility “snowballs” between *Solanum* species. Science 329, 1521–1523 (2010).

93. R. J. Wang, M. A. White, B. A. Payseur, The Pace of hybrid incompatibility evolution in house mice. Genetics 201, 229–242 (2015).

94. A. S. Kondrashov, S. Sunyaev, F. A. Kondrashov, Dobzhansky–Muller incompatibilities in protein evolution. PNAS 99, 14878–14883 (2002).

95. N. A. Johnson, A. H. Porter, Evolution of branched regulatory genetic pathways: Directional selection on pleiotropic loci accelerates developmental system drift. Genetica 129, 57–70 (2007).

96. A. Y. Tulchinsky, N. A. Johnson, A. H. Porter, Hybrid Incompatibility Despite Pleiotropic Constraint in a Sequence-Based Bioenergetic Model of Transcription Factor Binding. Genetics 198, 1645–1654 (2014).

97. K. L. Mack, M. W. Nachman, Gene regulation and speciation. Trends in Genetics 33, 68–80 (2017).

98. A. D. Cutter, J. D. Bundus, Speciation and the developmental alarm clock. eLife 9, e56276 (2020).

99. W. Chang, D. R. Matute, M. Kreitman, Rapid evolution of the functionally conserved gap gene *giant* in *Drosophila*. bioRxiv 2021.07.08.451553 (2021). 10.1101/2021.07.08.451553.

100. G. J□rgens, E. Wieschaus, C. Nusslein-Volhard, H. Kluding, Mutations affecting the pattern of the larval cuticle inDrosophila melanogaster: II. Zygotic loci on the third chromosome. Wilhelm Roux’ Archiv 193, 283–295 (1984).

101. D. L. Lindsley, G. G. Zimm, The genome of Drosophila melanogaster (1992).

102. D. R. Matute, I. A. Butler, D. A. Turissini, J. A. Coyne, A test of the snowball theory for the rate of evolution of hybrid incompatibilities. Science 329, 1518–1521 (2010).

103. J. A. Coyne, Genetic studies of three sibling species of Drosophila with relationship to theories of speciation. Genetical Research (1985). 10.1017/S0016672300022643.

104. D. C. Presgraves, C. D. Meiklejohn, Hybrid Sterility, Genetic Conflict and Complex Speciation: Lessons From the *Drosophila simulans* Clade Species. Front. Genet. 0 (2021).

105. J. A. Coyne, S. Elwyn, S. Y. Kim, A. Llopart, Genetic studies of two sister species in the *Drosophila melanogaster* subgroup, D. yakuba and D. santomea. Genetics Research 84, 11–26 (2004).

106. A. Llopart, E. Brud, N. Pettie, J. M. Comeron, Support for the Dominance Theory in Drosophila Transcriptomes. Genetics 210, 703–718 (2018).

107. R. C. Johnson, et al., Accounting for multiple comparisons in a genome-wide association study (GWAS). BMC genomics 11, 1–6 (2010).

108. H. a Orr, Haldane’s rule has multiple genetic causes. Nature 361, 532–533 (1993).

109. M. V. Cattani, D. C. Presgraves, Incompatibility between *X* chromosome factor and pericentric heterochromatic region causes lethality in hybrids between *Drosophila melanogaster* and its sibling species. Genetics 191, 549–559 (2012).

110. O. Nagy, et al., Correlated evolution of two copulatory organs via a single cis-regulatory nucleotide change. Current Biology 28, 3450–3457 (2018).

111. C. J. J. Miller, D. R. Matute, The Effect of Temperature on *Drosophila* Hybrid Fitness. G3 Genes|Genomes|Genetics 7, 377–385 (2017).

112. R. L. Rogers, et al., Landscape of standing variation for tandem duplications in *Drosophila yakuba* and *Drosophila simulans*. Molecular Biology and Evolution 31, 1750–1766 (2014).

113. R. L. Rogers, L. Shao, K. R. Thornton, Tandem duplications lead to novel expression patterns through exon shuffling in *Drosophila yakuba*. PLoS Genetics 13 (2017).

114. D. Garrigan, et al., Genome sequencing reveals complex speciation in the *Drosophila simulans* clade. Genome Research 22, 1499–1511 (2012).

115. C. L. Brand, S. B. Kingan, L. Wu, D. Garrigan, A selective sweep across species boundaries in *Drosophila*. Molecular Biology and Evolution (2013). 10.1093/molbev/mst123.

116. D. R. Schrider, J. Ayroles, D. R. Matute, A. D. Kern, Supervised machine learning reveals introgressed loci in the genomes of *Drosophila simulans* and *D. sechellia*. PLoS genetics (2018). 10.1371/journal.pgen.1007341.

117. A. Serrato-Capuchina, et al., P-elements strengthen reproductive isolation within the *Drosophila simulans* species complex. bioRxiv 2020.08.17.254169 (2020). 10.1101/2020.08.17.254169.

118. J. E. Pool, The mosaic ancestry of the *Drosophila* genetic reference panel and the D. melanogaster reference genome reveals a network of epistatic fitness interactions. Molecular biology and evolution 32, 3236–3251 (2015).

119. J. B. Lack, J. D. Lange, A. D. Tang, R. B. Corbett-Detig, J. E. Pool, A thousand fly genomes: An expanded *Drosophila* genome nexus. Molecular Biology and Evolution 33, 3308–3313 (2016).

120. H. Li, Aligning sequence reads, clone sequences and assembly contigs with BWA-MEM. arXiv preprint arXiv 00, 3 (2013).

121. H. Li, R. Durbin, Fast and accurate short read alignment with Burrows-Wheeler transform. Bioinformatics 25, 1754–1760 (2009).

122. Drosophila 12 Genomes Consortium, et al., Evolution of genes and genomes on the Drosophila phylogeny. Nature 450, 203–218 (2007).

123. T. T. Hu, M. B. Eisen, K. R. Thornton, P. Andolfatto, A second-generation assembly of the Drosophila simulans genome provides new insights into patterns of lineage-specific divergence. Genome Research 23, 89–98 (2013).

124. H. Li, et al., The sequence alignment/map format and SAMtools. Bioinformatics 25, 2078–2079 (2009).

125. A. McKenna, et al., The Genome Analysis Toolkit: A MapReduce framework for analyzing next-generation DNA sequencing data. Genome Res. 20, 1297–1303 (2010).

126. M. A. DePristo, et al., A framework for variation discovery and genotyping using next-generation DNA sequencing data. Nature genetics 43, 491–498 (2011).

127. R. A. Hoskins, et al., The Release 6 reference sequence of the Drosophila melanogaster genome. Genome research 25, 445–458 (2015).

128. W. Maddison, D. Maddison, Mesquite 2. Manual 1–258 (2010).

129. Z. Yang, PAML: a program package for phylogenetic analysis by maximum likelihood. Bioinformatics (1997). 10.1093/bioinformatics/13.5.555.

130. Z. Yang, PAML 4: Phylogenetic analysis by maximum likelihood. Molecular Biology and Evolution (2007). 10.1093/molbev/msm088.

131. A. Venkat, M. W. Hahn, J. W. Thornton, Multinucleotide mutations cause false inferences of lineage-specific positive selection. Nature Ecology and Evolution (2018). 10.1038/s41559-018-0584-5.

## SI REFERENCES

1. D. L. Lindsley, G. G. Zimm, The genome of Drosophila melanogaster (1992).

2. 2. W. Chang, D. R. Matute, M. Kreitman, Rapid evolution of the functionally conserved gap gene giant in Drosophila. bioRxiv, 2021.07.08.451553 (2021).

3. D. G. Gibson, et al., Enzymatic assembly of DNA molecules up to several hundred kilobases. Nature methods 6, 343–345 (2009).

4. S. T. Thibault, et al., A complementary transposon tool kit for Drosophila melanogaster using P and piggyBac. Nature genetics 36, 283–287 (2004).

5. R Core Team, R Development Core Team (2016).

6. J. A. Coyne, Genetic studies of three sibling species of Drosophila with relationship to theories of speciation. Genetical Research (1985) 10.1017/S0016672300022643.

7. D. C. Presgraves, C. D. Meiklejohn, Hybrid Sterility, Genetic Conflict and Complex Speciation: Lessons From the Drosophila simulans Clade Species. Front. Genet. 0 (2021).

8. J. A. Coyne, S. Elwyn, S. Y. Kim, A. Llopart, Genetic studies of two sister species in the Drosophila melanogaster subgroup, D. yakuba and D. santomea. Genetics Research 84, 11–26 (2004).

9. D. A. Turissini, G. Liu, J. R. David, D. R. Matute, The evolution of reproductive isolation in the Drosophila yakuba complex of species. J. Evol. Biol. 28, 557–575 (2015).

10. A. Llopart, E. Brud, N. Pettie, J. M. Comeron, Support for the Dominance Theory in Drosophila Transcriptomes. Genetics 210, 703–718 (2018).

11. K. J. T. Venken, et al., A molecularly defined duplication set for the X chromosome of Drosophila melanogaster. Genetics 186, 1111–1125 (2010).

12. D. R. Matute, J. Gavin-Smyth, Fine Mapping of Dominant X-Linked Incompatibility Alleles in Drosophila Hybrids. PLoS Genetics 10 (2014).

13. H. a Orr, Haldane’s rule has multiple genetic causes. Nature 361, 532–533 (1993).

14. M. V. Cattani, D. C. Presgraves, Incompatibility between X chromosome factor and pericentric heterochromatic region causes lethality in hybrids between Drosophila melanogaster and its sibling species. Genetics 191, 549–559 (2012).

15. O. Nagy, et al., Correlated evolution of two copulatory organs via a single cis-regulatory nucleotide change. Current Biology 28, 3450–3457 (2018).

16. J. A. Coyne, S. Simeonidis, P. Rooney, Relative paucity of genes causing inviability in hybrids between Drosophila melanogaster and D. simulans. Genetics 150, 1091–1103 (1998).

17. D. C. Presgraves, A fine-scale genetic analysis of hybrid incompatibilities in $\backslash$emph{Drosophila}. Genetics 163, 955–972 (2003).

18. D. R. Matute, I. A. Butler, D. A. Turissini, J. A. Coyne, A test of the snowball theory for the rate of evolution of hybrid incompatibilities. Science 329, 1518–1521 (2010).

19. C. J. J. Miller, D. R. Matute, The Effect of Temperature on Drosophila Hybrid Fitness. G3 Genes|Genomes|Genetics 7, 377–385 (2017).

20. J. Gavin-Smyth, D. R. Matute, Embryonic lethality leads to hybrid male inviability in hybrids between Drosophila melanogaster and D. santomea. Ecology and Evolution 3, 1580–1589 (2013).

21. R. L. Rogers, et al., Landscape of standing variation for tandem duplications in drosophila yakuba and drosophila simulans. Molecular Biology and Evolution 31, 1750–1766 (2014).

22. R. L. Rogers, L. Shao, K. R. Thornton, Tandem duplications lead to novel expression patterns through exon shuffling in Drosophila yakuba. PLoS Genetics 13 (2017).

23. D. A. Turissini, D. R. Matute, Fine scale mapping of genomic introgressions within the Drosophila yakuba clade. PLOS Genetics 13, e1006971 (2017).

24. D. A. Turissini, J. A. McGirr, S. S. Patel, J. R. David, D. R. Matute, The rate of evolution of postmating-prezygotic reproductive isolation in drosophila. Molecular Biology and Evolution 35, 312–334 (2018).

25. D. Garrigan, et al., Genome sequencing reveals complex speciation in the Drosophila simulans clade. Genome Research 22, 1499–1511 (2012).

26. C. L. Brand, S. B. Kingan, L. Wu, D. Garrigan, A selective sweep across species boundaries in Drosophila. Molecular Biology and Evolution (2013) 10.1093/molbev/mst123.

27. D. R. Schrider, J. Ayroles, D. R. Matute, A. D. Kern, Supervised machine learning reveals introgressed loci in the genomes of Drosophila simulans and D. sechellia. PLoS genetics (2018) 10.1371/journal.pgen.1007341.

28. A. Serrato-Capuchina, et al., P-elements strengthen reproductive isolation within the Drosophila simulans species complex. bioRxiv, 2020.08.17.254169 (2020).

29. J. E. Pool, The mosaic ancestry of the Drosophila genetic reference panel and the D. melanogaster reference genome reveals a network of epistatic fitness interactions. Molecular biology and evolution 32, 3236–3251 (2015).

30. J. B. Lack, J. D. Lange, A. D. Tang, R. B. Corbett-Detig, J. E. Pool, A thousand fly genomes: An expanded drosophila genome nexus. Molecular Biology and Evolution 33, 3308–3313 (2016).

31. H. Li, Aligning sequence reads, clone sequences and assembly contigs with BWA-MEM. arXiv preprint arXiv 00, 3 (2013).

32. H. Li, R. Durbin, Fast and accurate short read alignment with Burrows-Wheeler transform. Bioinformatics 25, 1754–1760 (2009).

33. Drosophila 12 Genomes Consortium, et al., Evolution of genes and genomes on the Drosophila phylogeny. Nature 450, 203–218 (2007).

34. T. T. Hu, M. B. Eisen, K. R. Thornton, P. Andolfatto, A second-generation assembly of the Drosophila simulans genome provides new insights into patterns of lineage-specific divergence. Genome Research 23, 89–98 (2013).

35. H. Li, et al., The sequence alignment/map format and SAMtools. Bioinformatics 25, 2078–2079 (2009).

36. A. McKenna, et al., The Genome Analysis Toolkit: A MapReduce framework for analyzing next-generation DNA sequencing data. Genome Res. 20, 1297–1303 (2010).

37. M. A. DePristo, et al., A framework for variation discovery and genotyping using next-generation DNA sequencing data. Nature genetics 43, 491–498 (2011).

38. G. Dos Santos, et al., FlyBase: Introduction of the Drosophila melanogaster Release 6 reference genome assembly and large-scale migration of genome annotations. Nucleic Acids Research 43, D690–D697 (2015).

39. W. Maddison, D. Maddison, Mesquite 2. Manual, 1–258 (2010).

40. E. D. Eldon, V. Pirrotta, Interactions of the Drosophila gap gene giant with maternal and zygotic pattern-forming genes. Development (Cambridge, England) 111, 367–378 (1991).

41. T. R. Hartman, et al., Drosophila Boi limits hedgehog levels to suppress follicle stem cell proliferation. Journal of Cell Biology 191, 943–952 (2010).

42. J. B. Skeath, et al., The extracellular metalloprotease AdamTS-A anchors neural lineages in place within and preserves the architecture of the central nervous system. Development (2017) 10.1242/dev.145854.

43. J. M. Toivonen, et al., Technical knockout, a drosophila model of mitochondrial deafness. Genetics (2001).

44. A. Serrato-Capuchina, et al., Paternally Inherited P-Element Copy Number Affects the Magnitude of Hybrid Dysgenesis in Drosophila simulans and D. melanogaster. Genome Biol Evol 12, 808–826 (2020).

45. K. Sawamura, M. T. Yamamoto, Cytogenetical localization of Zygotic hybrid rescue (Zhr), a Drosophila melanogaster gene that rescues interspecific hybrids from embryonic lethality. MGG Molecular & General Genetics (1993) 10.1007/BF00276943.

