## Supplementary material for "Gap genes are involved in inviability in hybrids between *Drosophila melanogaster and D. santomea*": Tables S1-S14 Figures S1-S21

Daniel R. Matute

Daniel R. Matute

**This PDF file includes:**

Tables S1 to S14

Figures S1 to S21

SI References

**SUPPLEMENTARY TABLES**

**TABLE S1.** **Rates of ablation in F1 hybrids between the species of the *yakuba* species complex.** None of the nine possible F1 hybrids show abdominal ablations.

| **Cross** | **Scored embryos** | **Dead embryos** | **Ablated embryos** |
| --- | --- | --- | --- |
| **♀ *D. santomea ×* ♂ *D. santomea*** | 120 | 2 | 0 |
| **♀ *D. yakuba ×* ♂ *D. yakuba*** | 162 | 4 | 0 |
| **♀ *D. teissieri ×* ♂ *D. teissieri*** | 149 | 7 | 0 |
| **♀ *D. santomea ×* ♂ *D. yakuba*** | 102 | 16 | 0 |
| **♀ *D. santomea ×* ♂ *D. teissieri*** | 110 | 21 | 0 |
| **♀ *D. yakuba ×* ♂ *D. santomea*** | 123 | 19 | 0 |
| **♀ *D. yakuba ×* ♂ *D. teissieri*** | 98 | 16 | 0 |
| **♀ *D. teissieri ×* ♂ *D. santomea*** | 101 | 15 | 0 |
| **♀ *D. teissieri ×* ♂ *D. yakuba*** | 98 | 10 | 0 |

**TABLE S2. Hybrids between *D. melanogaster* females and *D. simulans* and *D. mauritiana* show no abdominal ablations or embryonic lethality.** The reciprocal crosses (♀*D. simulans* × ♂ *D. melanogaster;* ♀*D. mauritiana* × ♂*D. melanogaster*) show high levels of female embryonic lethality but eggs do not turn brown (i.e., the assay of brown embryos cannot detect early embryo lethality).

| **Cross** | **Hatched embryos** | **Dead brown embryos** | **Ablated embryos** |
| --- | --- | --- | --- |
| **♀ *D. melanogaster ×* ♂ *D. melanogaster*** | 110 | 6 | 0 |
| **♀ *D. simulans ×* ♂ *D. simulans*** | 105 | 5 | 0 |
| **♀ *D. sechellia ×* ♂ *D. sechellia*** | 88 | 8 | 0 |
| **♀ *D. mauritiana ×* ♂ *D. mauritiana*** | 121 | 4 | 0 |
| **♀ *D. melanogaster ×* ♂ *D. simulans*** | 98 | 5 | 0 |
| **♀ *D. melanogaster ×* ♂ *D. mauritiana*** | 74 | 5 | 0 |
| **♀ *D. melanogaster ×* ♂ *D. sechellia*** | 99 | 41 | 0 |
| **♀ *D. simulans ×* ♂ *D. melanogaster*** | 101 | 6 | 0 |
| **♀ *D. simulans ×* ♂ *D. sechellia*** | 78 | 1 | 0 |
| **♀ *D. simulans ×* ♂ *D. mauritiana*** | 95 | 2 | 0 |
| **♀ *D. sechellia ×* ♂ *D. melanogaster*** | 73 | 2 | 0 |
| **♀ *D. sechellia ×* ♂ *D. simulans*** | 77 | 4 | 0 |
| **♀ *D. sechellia ×* ♂ *D. mauritiana*** | 71 | 6 | 0 |
| **♀ *D. mauritiana ×* ♂ *D. sechellia*** | 83 | 4 | 0 |
| **♀ *D. mauritiana ×* ♂ *D. simulans*** | 99 | 2 | 0 |
| **♀ *D. mauritiana ×* ♂ *D. melanogaster*** | 68 | 4 | 0 |

**TABLE S3. Mutant stocks used in this study.**

|  | **Stock number** | **Genotype** |
| --- | --- | --- |
| **1** | 53 | gt^1^ w^a^ |
| **2** | 54 | gt^13z^/Dp(1;2;Y)w^+^/C(1)DX, y^1^ f^1^ |
| **3** | 1528 | gt^[Q292]^ rst^[6]^/FM7a |
| **4** | 1529 | y^[1]^ sc^[1]^ gt^[X11]^/FM6 |
| **5** | 1530 | y^[1]^ gt^[E6]^ rst^[6]^ |
| **6** | 35 | dor^4^/C(1)RM, y^1^ w^1^ f^1^ |
| **7** | 3721 | C(1)DX, y^1^ f^1^/FM6; ry^506^ |
| **8** | 17003 | P{EP}boiEP1385 w^1118^ |
| **9** | 13245 | y^1^ P{SUPor-P}boi^KG03233^ |
| **10** | 4283 | tko^3^/FM7a/Dp(1;2;Y)w^+^ |
| **11** | 59642 | y^1^ Mi{MIC}tko^MI15120^ w^*^/FM7h |
| **12** | 52400 | y^1^ tko^A^ w^*^ P{neoFRT}19A/FM7c, P{GAL4-Kr.C}DC1, P{UAS-GFP.S65T}DC5, sn^+^ |
| **13** | 52401 | y^1^ tkoB w* P{neoFRT}19A/FM7c, P{GAL4-Kr.C}DC1, P{UAS-GFP.S65T}DC5, sn^+^ |
| **14** | 9372 | w^*^ sd^11^/FM7i, P{ActGFP}JMR3 |
| **15** | 29799 | Dp(1;Y)BSC75, y^+^P{w[+mW.Scer\FRT.hs3]=3'.RS5+3.3'}BSC3, B^S^/Df(1)ED6630, P{w[+mW.Scer\FRT.hs3]=3'.RS5+3.3'}ED6630 w^1118^/C(1)RA, In(1)sc[J1], In(1)sc[8], l(1)1Ac^1^, sc^J1^ sc^8^ |
| **16** | 29801 | Dp(1;Y)BSC77, y^+^ P{w[+mW.Scer\FRT.hs3]=3'.RS5+3.3'}BSC3, B[S]/Df(1)ED6630, P{w[+mW.Scer\FRT.hs3]=3'.RS5+3.3'}ED6630 w^1118^/C(1)RA, In(1)sc[J1], In(1)sc^8^, l(1)1Ac^1^, sc^J1^ sc^8^ |
| **17** | 29803 | Dp(1;Y)BSC79, y^+^ P{w[+mW.Scer\FRT.hs3]=3'.RS5+3.3'}BSC3, B[S]/Df(1)ED6630, P{w[+mW.Scer\FRT.hs3]=3'.RS5+3.3'}ED6630 w^1118^/C(1)RA, In(1)sc[J1], In(1)sc[8], l(1)1Ac^1^, sc^J1^ sc^8^ |
| **18** | 29804 | Dp(1;Y)BSC80, y^+^ P{w[+mW.Scer\FRT.hs3]=3'.RS5+3.3'}BSC3, B[S]/Df(1)ED6630, P{w[+mW.Scer\FRT.hs3]=3'.RS5+3.3'}ED6630 w^1118^/C(1)RA, In(1)sc^J1^, In(1)sc8, l(1)1Ac^1^, sc^J1^ sc^8^ |
| **19** | 29805 | Dp(1;Y)BSC81, y[+] P{w[+mW.Scer\FRT.hs3]=3'.RS5+3.3'}BSC3, B[S]/Df(1)ED6630, P{w[+mW.Scer\FRT.hs3]=3'.RS5+3.3'}ED6630 w^1118^/C(1)RA, In(1)sc[J1], In(1)sc[8], l(1)1Ac^1^, sc^J1^ sc^8^ |
| **20** | 29806 | Dp(1;Y)BSC82, y[+] P{w[+mW.Scer\FRT.hs3]=3'.RS5+3.3'}BSC3, B^S^/Df(1)Exel6233, w[1118] P{w[+mC]=XP-U}Exel6233/C(1)RA, In(1)sc[J1], In(1)sc[8], l(1)1Ac^1^, sc^J1^ sc^8^ |
| **21** | 29807 | Dp(1;Y)BSC83, y[+] P{w[+mW.Scer\FRT.hs3]=3'.RS5+3.3'}BSC3, B[S]/Df(1)Exel6233, w[1118] P{w[+mC]=XP-U}Exel6233/C(1)RA, In(1)sc[J1], In(1)sc[8], l(1)1Ac^1^, sc^J1^ sc^8^ |
| **22** | 29808 | Dp(1;Y)BSC84, y[+] P{w[+mW.Scer\FRT.hs3]=3'.RS5+3.3'}BSC3, B[S]/Df(1)Exel6233, w^1118^ P{w[+mC]=XP-U}Exel6233/C(1)RA, In(1)sc[J1], In(1)sc^8^, l(1)1Ac^1^, sc^J1^ sc^8^ |
| **23** | 29809 | Dp(1;Y)BSC85, y[+] P{w[+mW.Scer\FRT.hs3]=3'.RS5+3.3'}BSC3, B[S]/Df(1)Exel6233, w^1118^ P{w[+mC]=XP-U}Exel6233/C(1)RA, In(1)sc[J1], In(1)sc[8], l(1)1Ac^1^, sc^J1^ sc^8^ |
| **24** | 29811 | Dp(1;Y)BSC87, y^+^ P{w[+mW.Scer\FRT.hs3]=3'.RS5+3.3'}BSC3, B[S]/Df(1)Exel6233, w^1118^ P{w[+mC]=XP-U}Exel6233/C(1)RA, In(1)sc[J1], In(1)sc^8^, l(1)1Ac^1^, sc^J1^ sc^8^ |
| **25** | 29812 | Dp(1;Y)BSC88, y^+^ P{w[+mW.Scer\FRT.hs3]=3'.RS5+3.3'}BSC3, B^S^/Df(1)Exel6233, w^1118^ P{w[+mC]=XP-U}Exel6233/C(1)RA, In(1)sc^J1^, In(1)sc^8^, l(1)1Ac^1^, sc^J1^ sc^8^ |
| **26** | 29814 | Dp(1;Y)BSC90, y^+^P{w[+mW.Scer\FRT.hs3]=3'.RS5+3.3'}BSC3, B^S^/winscy/C(1)RA, In(1)sc^J1^, In(1)sc^8^, l(1)1Ac^1^, sc^J1^ sc^8^ |
| **27** | 32295 | w^1118^; Dp(1;3)DC454, PBac{y[+mDint2] w[+mC]=DC454}VK00033/TM6C, Sb^1^ |
| **28** | 30233 | w^1118^; Dp(1;3)DC045, PBac{y[+mDint2] w[+mC]=DC045}VK00033 |
| **29** | 30751 | w^1118^; Dp(1;3)DC107, PBac{y[+mDint2] w[+mC]=DC107}VK00033 |
| **30** | 38476 | w^1118^; Dp(1;3)RC013, PBac{y[+mDint2] w[+mC]=RC013}VK00033/TM6C, Sb^1^ |
| **31** | 31455 | w^1118^; Dp(1;3)DC405, PBac{y[+mDint2] w[+mC]=DC405}VK00033 |
| **32** | 32294 | w^1118^; Dp(1;3)DC453, PBac{y[+mDint2] w[+mC]=DC453}VK00033/TM6C, Sb^1^ |
| **33** | 8950 | Df(1)ED409, P{w[+mW.Scer\FRT.hs3]=3'.RS5+3.3'}ED409 w^1118^/FM7h |
| **34** | 9345 | Df(1)ED11354, P{w[+mW.Scer\FRT.hs3]=3'.RS5+3.3'}ED11354 w^1118^/FM7h |
| **35** | 9054 | Df(1)ED11354, P{w[+mW.Scer\FRT.hs3]=3'.RS5+3.3'}ED11354 w^1118^/FM7h |
| **36** | 8031 | Df(1)ED411, P{w[+mW.Scer\FRT.hs3]=3'.RS5+3.3'}ED411 w^1118^/FM7j, B^1^ |
| **37** | 26569 | Df(1)BSC717, P+PBac{w[+mC]=XP3.RB5}BSC717 w^1118^/FM7h/Dp(2;Y)G, P{w[+mC]=hs-hid}Y |
| **38** | 7705 | Df(1)Exel6230, P{XP-U}Exel6230 w^1118^/FM7c |
| **39** | 7706 | Df(1)Exel6231, P{XP-U}Exel6231 w^1118^/FM7c |

**TABLE S4.** **Isofemale lines from four species used to assess whether HI in *mel/san* hybrids was a species-specific phenomenon or a line specific phenomenon.** All lines were collected by D. R. Matute with the exception of the three *D. teissieri* lines marked with an asterisk. Those three lines were donated by J.R. David.

MISSING THE MELANOGASTER LINES (ALSO SIM>????)

| **Species** | **Line** | **Notation in Figures S4 and S18** | **Location** | **Year** |
| --- | --- | --- | --- | --- |
| *D. santomea* | Thena5 | S1 | São Tomé | 2005 |
| *D. santomea* | sanCAR1490 | S2 | São Tomé | 2005 |
| *D. santomea* | sanCOST1250.5 | S3 | São Tomé | 2009 |
| *D. santomea* | sanCOST1270.1 | S4 | São Tomé | 2009 |
| *D. santomea* | sanOBAT1200 | S5 | São Tomé | 2005 |
| *D. santomea* | sanOBAT1200.2 | S6 | São Tomé | 2009 |
| *D. santomea* | sanRain39 | S7 | São Tomé | 2009 |
| *D. santomea* | sanCAR1600.3 | S8 | São Tomé | 2009 |
| *D. santomea* | Carv2015.1 | S9 | São Tomé | 2015 |
| *D. santomea* | Carv2015.5 | S10 | São Tomé | 2015 |
| *D. santomea* | Carv2015.11 | S11 | São Tomé | 2015 |
| *D. santomea* | Carv2015.16 | S12 | São Tomé | 2015 |
| *D. santomea* | Pico1680.1 | S13 | São Tomé | 2015 |
| *D. santomea* | Pico1659.2 | S14 | São Tomé | 2015 |
| *D. santomea* | Pico1659.3 | S15 | São Tomé | 2015 |
| *D. santomea* | Amelia2015.1 | S16 | São Tomé | 2015 |
| *D. santomea* | Amelia2015.6 | S17 | São Tomé | 2015 |
| *D. santomea* | Amelia2015.12 | S18 | São Tomé | 2015 |
| *D. santomea* | A1200.7 | S19 | São Tomé | 2009 |
| *D. santomea* | Rain42 | S20 | São Tomé | 2009 |
| *D. teissieri* | Balancha_1 | T1 | Bioko, Equatorial Guinea | 2013 |
| *D. teissieri* | Balancha_2 | T2 | Bioko, Equatorial Guinea | 2013 |
| *D. teissieri* | Balancha_3 | T3 | Bioko, Equatorial Guinea | 2013 |
| *D. teissieri* | House_Bioko_0 | T4 | Bioko, Equatorial Guinea | 2013 |
| *D. teissieri* | House_Bioko_1 | T5 | Bioko, Equatorial Guinea | 2013 |
| *D. teissieri* | House_Bioko_2 | T6 | Bioko, Equatorial Guinea | 2013 |
| *D. teissieri* | La_Lope_Gabon* | T7 | Gabón | ~1975 |
| *D. teissieri* | Selinda* | T8 | Gabón | ~1975 |
| *D. teissieri* | Zimbabwe* | T9 | Gabón | ~1975 |
| *D. teissieri* | cascade_2_1 | T10 | Bioko, Equatorial Guinea | 2013 |
| *D. teissieri* | cascade_2_2 | T11 | Bioko, Equatorial Guinea | 2013 |
| *D. teissieri* | cascade_2_4 | T12 | Bioko, Equatorial Guinea | 2013 |
| *D. teissieri* | cascade_4_1 | T13 | Bioko, Equatorial Guinea | 2013 |
| *D. teissieri* | cascade_4_2 | T14 | Bioko, Equatorial Guinea | 2013 |
| *D. teissieri* | cascade_4_3 | T15 | Bioko, Equatorial Guinea | 2013 |
| *D. teissieri* | cascade_4_4 | T16 | Bioko, Equatorial Guinea | 2013 |
| *D. teissieri* | cascade_4_5 | T17 | Bioko, Equatorial Guinea | 2013 |
| *D. teissieri* | cascade_4_6 | T18 | Bioko, Equatorial Guinea | 2013 |
| *D. teissieri* | Bata_2 | T19 | Bata, Equatorial Guinea | 2009 |
| *D. teissieri* | Bata_8 | T20 | Bata, Equatorial Guinea | 2009 |
| *D. simulans* | MD06 | NA | Madagascar | NA |
| *D. simulans* | MD105 | NA | Madagascar | NA |
| *D. simulans* | MD106 | NA | Madagascar | NA |
| *D. simulans* | MD15 | NA | Madagascar | NA |
| *D. simulans* | MD199 | NA | Madagascar | NA |
| *D. simulans* | MD221 | NA | Madagascar | NA |
| *D. simulans* | MD233 | NA | Madagascar | NA |
| *D. simulans* | MD251 | NA | Madagascar | NA |
| *D. simulans* | MD63 | NA | Madagascar | NA |
| *D. simulans* | MD73 | NA | Madagascar | NA |
| *D. simulans* | NS05 | NA | Nairobi, Kenya | NA |
| *D. simulans* | NS113 | NA | Nairobi, Kenya | NA |
| *D. simulans* | NS137 | NA | Nairobi, Kenya | NA |
| *D. simulans* | NS33 | NA | Nairobi, Kenya | NA |
| *D. simulans* | NS39 | NA | Nairobi, Kenya | NA |
| *D. simulans* | NS40 | NA | Nairobi, Kenya | NA |
| *D. simulans* | NS50 | NA | Nairobi, Kenya | NA |
| *D. simulans* | NS67 | NA | Nairobi, Kenya | NA |
| *D. simulans* | NS78 | NA | Nairobi, Kenya | NA |
| *D. simulans* | NS79 | NA | Nairobi, Kenya | NA |

**TABLE S5. *C(1)DX, Dp(1;3)/TM3*** **lay few eggs than *C(1)DX, Dp(1;Y)* females in conspecific *mel* matings.** Each column shows the median number of eggs produced by a singly-mated female when mated to a *w^1118^* male over 4 days (*N* = 15 females).

| **Duplication** | **Stock** | ***C(1)DX, Dp(1;3)/Dp(1;3)*** | ***C(1)DX, Dp(1;3)/TM3,Sb*** | ***C(1)DX, Dp(1;Y)*** |
| --- | --- | --- | --- | --- |
| ***Dp(1;3)DC405*** | 31455 | 47 | 7 | NA |
| ***Dp(1;3)DC301*** | 31452 | 51 | 24 | NA |
| ***Dp(1;3)DC112*** | 31445 | 43 | 15 | NA |
| ***Dp(1;3)DC272*** | 30389 | 24 | 20 | NA |
| ***Dp(1;Y)BSC87*** | 29811 | NA | NA | 56 |
| ***Dp(1;Y)*** ***BSC170*** | 32117 | NA | NA | 68 |
| ***Dp(1;Y)BSC58*** | 29782 | NA | NA | 53 |
| ***Dp(1;Y)BSC14*** | 29782 | NA | NA | 64 |

**TABLE S6. *Y*-linked duplications of the *X*-chromosome rescue a *gt_mel_* null allele more effectively than duplications on the third chromosome in a pure *mel* background.** All crosses involved *gt_mel_^X11^/FM7* females and hemizygote males for two different types of duplications **(***Dp(1;Y)* or *Dp(1;3)*). Cells show the total progeny from 15 crosses. In the case of *Dp(1;Y)* duplications, the father had a *X_mel_/Y_mel_Dp(1;Y)* genotype. In the case of *Dp(1;3*) duplication, the father had a *X_mel_/Y_mel_*, *Dp(1,3)/ Dp(1,3)* genotype. I only scored *gt^X11^*-carriers.

| Genotype | Stock | *Dp(1;Y)* or *Dp(1;3)*? | F1 male genotypes | | Rescue Rate | *X^2^* | P |
| --- | --- | --- | --- | --- | --- | --- | --- |
|  |  |  | *gt^X11^*/ Dp | *FM7*/Dp |  |  |  |
| *Dp(1;Y)BSC74* | 29798 | *Dp(1;Y)* | 81 | 73 | 0.526 | 0.117 | 0.732 |
| *Dp(1;Y)BSC78* | 29802 | *Dp(1;Y)* | 53 | 48 | 0.525 | 0.045 | 0.833 |
| *Dp(1;3)DC405* | 31455 | *Dp(1;3)* | 41 | 75 | 0.353 | 4.511 | 0.034 |
| *Dp(1;3)RC013* | 38476 | *Dp(1;3)* | 16 | 37 | 0.302 | 3.545 | 0.060 |

**TABLE S7**. **Complementation mapping using loss-of function and hypomoprhic alleles show that *gt* mutants are the only alleles in cytological region 3A3 that lead to an increase of female viability in *mel/san* hybrid.** *mutant/san* refers to the number of F1 hybrids carrying the mutant allele. *gt^X11^* is a loss of function (amorphic) allele (2, 40). Both *boi^d04197^* ^and^ *boi^e01708^* shows significantly lower RNA production than wild-type flies (41). *trol^G0271^*is a hypomorph (42). *tko^3^* is a null allele. *tko^3^*/*tko^3^* homozygote females and *tko^3^* hemizygote males die as larvae but are able to complete embryogenesis (43).

| **Gene** | **Allele** | ***FM7::GFP/san*** | ***mutant/san*** | ***Ratio***  ***(F1 mutant/san)/Total*** |
| --- | --- | --- | --- | --- |
| *Giant* | *gt^X11^* | 256 | 610 | 0.704 |
| *Boi* | P{XP}*boi*^d04197^ | 440 | 478 | 0.521 |
| *boi* | PBac{RB}*boi*^e01708^ | 561 | 610 | 0.521 |
| *troll* | *trol^G0271^* | 510 | 464 | 0.476 |
| *tko* | *tko^3^* | 501 | 457 | 0.477 |

**TABLE S8. Haplotypes associated with HI in *mel/san* hybrids. This table only shows autosomal haplotypes besides the one that contains *tll^mel^* (.xls file).**

**TABLE S9.** ***tll^mel^* exacerbates the defects caused by *gt^mel^* in *mel/san* hybrids as revealed by a second *tll* mutant**. The analysis is similar to that shown in Table 1. **A.** Female viability **B.** Frequency of abdominal ablations in hybrid males. **C.** Frequency of abdominal ablations in hybrid females.

| A. **Female hybrid viability** | | | | |
| --- | --- | --- | --- | --- |
|  | *FM7; sqh::mCherry/X^san^; TM3, Act::GFP, Ser/3^san^* | *FM7; sqh::mCherry/X^san^; tll^1^/3^san^* | *gt^X11^; /X^san^; TM3, Act::GFP, Ser/3^san^* | *gt^X11^; /X^san^; tll^1^/3^san^* |
|  | 18 | 22 | 34 | 54 |
|  | χ^2^_1_=0.40, P=0.527 | | χ^2^_1_=4.55, P=0.033 | |
| B. **Proportion of male embryos showing abdominal ablations (100 embryos each)** | | | | |
|  | *FM7; sqh::mCherry/Y^san^; TM3, Act::GFP, Ser/3^san^* | *FM7; sqh::mCherry/Y^san^; tll^1^/3^san^* | *gt^X11^; /Y^san^; TM3, Act::GFP, Ser/3^san^* | *gt^X11^; /Y^san^; tll^1^/3^san^* |
|  | 94 | 88 | 23 | 9 |
|  | χ^2^_1_= 1.526, P= 0.217 | | χ^2^_1_= 6.287, P = 0.012 | |
| C. **Proportion of female embryos showing abdominal ablations (100 embryos each)** | | | | |
|  | *FM7; sqh::mCherry/X^san^; TM3, Act::GFP, Ser/3^san^* | *FM7; sqh::mCherry/X^san^; tll^1^/3^san^* | *gt^X11^; /X^san^; TM3, Act::GFP, Ser/3^san^* | *gt^X11^; /X^san^; tll^1^/3^san^* |
|  | 48 | 35 | 16 | 6 |
|  | χ^2^_1_= 2.966, P= 0.08505 | | χ^2^_1_= 4.137, P= 0.0420 | |

**TABLE S10**. **Double mutant analysis of survival in hybrids between *D. melanogaster* females and three males from four different species (*D. melanogaster*, *D. simulans*, *D. mauritiana*, and *D. teissieri*).** Individuals that are hemizygote for both alleles do not show different viability from individuals that are hemizygote for one allele and heterozygote for the other one, or heterozygote for both alleles.

| *tll^1^* | | | | | | |
| --- | --- | --- | --- | --- | --- | --- |
|  | *FM7; sqh::mCherry/X^san^; TM3, Act::GFP, Ser/3^san^* | *FM7; sqh::mCherry/X^san^; tll^1^/3^san^* | *gt^X11^; /X^san^; TM3, Act::GFP, Ser/3^san^* | *gt^X11^; /X^san^; tll^1^/3^san^* | χ^2^, df =3 | P-value |
| *D. melanogaster* | 95 | 104 | 121 | 98 | 1.8695 | 0.5999 |
| *D. simulans* | 34 | 31 | 28 | 29 | 0.33864 | 0.9526 |
| *D. mauritiana* | 52 | 41 | 39 | 41 | 1.1436 | 0.7666 |
| *D. teissieri* | 19 | 22 | 15 | 20 | 0.71574 | 0.8695 |
| *tll*^Δ^*^GFP^* | | | | | | |
|  | *FM7; sqh::mCherry/X^san^; TM3, Act::GFP, Ser/3^san^* | *FM7; sqh::mCherry/X^san^; tll^1^/3^san^* | *gt^X11^; /X^san^; TM3, Act::GFP, Ser/3^san^* | *gt^X11^; /X^san^; tll^1^/3^san^* | χ^2^ |  |
| *D. melanogaster* | 104 | 87 | 99 | 78 | 2.1428 | 0.5433 |
| *D. simulans* | 42 | 36 | 39 | 44 | 0.46075 | 0.9274 |
| *D. mauritiana* | 51 | 38 | 49 | 43 | 1.1815 | 0.7575 |
| *D. teissieri* | 20 | 25 | 21 | 26 | 0.56722 | 0.9039 |

**TABLE S11**. **A *tll_san_^ΔdsRed^* has no effect on HI in crosses between male-carriers and *mel* females from four different backgrounds.**

|  | ***tll^ΔdsRed^/3_mel_*** | ***3_san_/3_mel_*** | **χ^2^** | **P-value** |
| --- | --- | --- | --- | --- |
| *mel AkLa* | 31 | 36 | 0.067 | 0.795 |
| *mel Zs2* | 62 | 74 | 1.600 | 0.206 |
| *mel Senegal* | 49 | 41 | 0.200 | 0.654 |
| *mel NC103* | 62 | 71 | 0.184 | 0.668 |

**TABLE S12.** **Abrogating the *tll_san_* allele has no viability effect in *gt_mel_^X11^ mel*/san hybrids.**

| A. **Female hybrid viability** | | | | |
| --- | --- | --- | --- | --- |
|  | *FM7; Act::GFP/X_san_; 3_mel_, tll_san_^dsRed^* | *FM7; Act::GFP/X_san_; 3_mel_, 3_san_* | *gt_mel_^X11^/X_san_; 3_mel_, tll_san_^dsRed^* | *gt_mel_^X11^/X_san_; 3_mel_, 3_san_* |
|  | 12 | 20 | 41 | 44 |
|  | χ^2^_1_=0.571, P=0.450 | | χ^2^_1_=0.006, P=0.939 | |

**TABLE S13. *gt* and *tll* are slowly evolving genes as measured by the rate of aminoacid substitutions.** I show K_A_/K_S_ values for two species comparisons (*D. melanogaster* vs. *D. santomea*, and *D. melanogaster* vs. *D. yakuba*) and the quantiles of the K_A_/K_S_ values compared to the rest of the genome. I used four different parametrizations in PAML (listed in the first column). *D. melanogaster*: *mel*, *D. santomea*: *san*; *D. yakuba*: *yak*.

| *Giant* |  |  |  |  |  |  |  |  |  |
| --- | --- | --- | --- | --- | --- | --- | --- | --- | --- |
| Parameter model | CG | K_A_  *mel* vs.*san* | K_S_  *mel* vs. *san* | K_A_/K_s_  *mel* vs. *san* | Quantile  K_A_ *mel* vs. *san* | K_A_  *mel* vs. *yak*_ | K_S_  *mel* vs. *yak* | K_A_/K_s_  *mel* vs. *yak* | Quantile  K_A_ *mel* vs. *yak* |
| 2_ratios | CG7952 | 0.0062 | 0.2192 | 0.0283 | 0.2227 | 0.0060 | 0.2074 | 0.0289 | 0.2281 |
| 3_ratios | CG7952 | 0.0062 | 0.2189 | 0.0283 | 0.2427 | 0.0060 | 0.2071 | 0.0290 | 0.2499 |
| basic_model | CG7952 | 0.0055 | 0.2218 | 0.0248 | 0.1869 | 0.0052 | 0.2102 | 0.0247 | 0.1862 |
| free_ratios | CG7952 | 0.0055 | 0.2222 | 0.0248 | 0.2278 | 0.0044 | 0.2139 | 0.0206 | 0.1960 |
| *Tailless* |  |  |  |  |  |  |  |  |  |
| Parameter model | CG | K_A_  *mel* vs.*san* | K_S_  *mel* vs. *san* | K_A_/K_s_  *mel* vs. *san* | Quantile  K_A_ *mel* vs. *san* | K_A_  *mel* vs. *yak*_ | K_S_  *mel* vs. *yak* | K_A_/K_s_  *mel* vs. *yak* | Quantile  K_A_ *mel* vs. *yak* |
| 2_ratios | CG1378 | 0.0045 | 0.384 | 0.012 | 0.095 | 0.0045 | 0.376 | 0.012 | 0.0977 |
| 3_ratios | CG1378 | 0.0034 | 0.395 | 0.009 | 0.086 | 0.0034 | 0.383 | 0.009 | 0.0872 |
| basic_model | CG1378 | 0.0037 | 0.389 | 0.010 | 0.075 | 0.0037 | 0.380 | 0.010 | 0.076 |
| free_ratios | CG1378 | 0.0028 | 0.389 | 0.007 | 0.087 | 0.0028 | 0.381 | 0.007 | 0.091 |

**TABLE S14**. **Sequencing details and coverage for all the lines included in this study.**

| **Species** | **Line** | **Read type** | **Average coverage** | **Source** | **SRA** |
| --- | --- | --- | --- | --- | --- |
| *D. mauritiana* | mau12w | pe | 153.67 |  | SRR1555246,SRR1560430, SRR1560444, SRR483621 |
| *D. mauritiana* | MauKiti | se | 13.86 |  |  |
| *D. mauritiana* | mauST | se | 3.11 |  |  |
| *D. mauritiana* | MS17 | se,pe | 60.56 |  | SRR556195, SRR556206, SRR556199, SRR556196 |
| *D. mauritiana* | R23 | pe | 115.98 |  | SRR1560090, SRR1560089, SRR1560087 |
| *D. mauritiana* | R31 | pe | 99.96 |  | SRR1560098, SRR1560097, SRR1560095 |
| *D. mauritiana* | R32 | pe | 120.55 |  | SRR1560102, SRR1560100, SRR1560103 |
| *D. mauritiana* | R39 | pe | 116.33 |  | SRR1560110, SRR1560109, SRR1560108 |
| *D. mauritiana* | R41 | pe | 145.38 |  | SRR1560130, SRR1560132, SRR1560131 |
| *D. mauritiana* | R44 | pe | 122.59 |  | SRR1560147, SRR1560146, SRR1560133 |
| *D. mauritiana* | R56 | pe | 121.66 |  | SRR1560150, SRR1560149, SRR1560148 |
| *D. mauritiana* | R61 | pe | 140.32 |  | SRR1560268, SRR1560267, SRR1560269 |
| *D. mauritiana* | R8 | pe | 90.27 |  | SRR1560276, SRR1560275 |
| *D. santomea* | Qiuja630.39 | se | 24.16 | (23, 24) | SRX3029341 |
| *D. santomea* | Quija37 | se | 11.74 | (23, 24) | SRX3029336 |
| *D. santomea* | sanC1350.14 | se | 18.62 | (23, 24) | SRX3029340 |
| *D. santomea* | sanCAR1490.5 | se | 15.77 | (23, 24) | SRX3029339 |
| *D. santomea* | sanCOST1250.5 | se | 13.27 | (23, 24) | SRX3029322 |
| *D. santomea* | sanCOST1270.6 | se | 14.76 | (23, 24) | SRX3029337 |
| *D. santomea* | sanOBAT1200.13 | se | 14.47 | (23, 24) | SRX3029334 |
| *D. santomea* | sanOBAT1200.5 | se | 16.82 | (23, 24) | SRX3029332 |
| *D. santomea* | sanRain39 | se | 15.81 | (23, 24) | SRX3029333: |
| *D. santomea* | sanSTO7 | se | 15.29 | (23, 24) | SRX3029335 |
| *D. santomea* | sanThena5 | se | 12.98 | (23, 24) | SRX3029338 |
| *D. sechellia* | Anro_B1 | pe | 36.85 | (24, 27) | SRX3029286 |
| *D. sechellia* | Anro_B2 | pe | 34.56 | (24, 27) | SRX3029285 |
| *D. sechellia* | Anro_B3 | pe | 38.75 | (24, 27) | SRX3029281 |
| *D. sechellia* | Anro_B5 | pe | 38.36 | (24, 27) | SRX3029270 |
| *D. sechellia* | Anro_B6 | pe | 33.48 | (24, 27) | SRX3029283 |
| *D. sechellia* | Anro_B7 | pe | 39.25 | (24, 27) | SRX3029301 |
| *D. sechellia* | Anro_B8 | pe | 34.84 | (24, 27) | SRX3029373 |
| *D. sechellia* | Denis124 | se | 24.73 | (24, 27) | SRX3029315 |
| *D. sechellia* | Denis135 | se | 32.4 | (24, 27) | SRX3029275 |
| *D. sechellia* | Denis7_2 | se | 28 | (24, 27) | SRX3029314 |
| *D. sechellia* | Denis7_8 | se | 28 | (24, 27) | SRX3029277 |
| *D. sechellia* | DenisAMT | se | 14.44 | (24, 27) | SRX3029303 |
| *D. sechellia* | DenisAT3 | se | 28.03 | (24, 27) | SRX3029317 |
| *D. sechellia* | DenisDNJ6 | se | 22.63 | (24, 27) | SRX3029319 |
| *D. sechellia* | DenisJT1 | se | 23.99 | (24, 27) | SRX3029316 |
| *D. sechellia* | DenisMCL | se | 46.04 | (24, 27) | SRX3029276 |
| *D. sechellia* | DenisNF100 | se | 10.12 | (24, 27) | SRX3029307 |
| *D. sechellia* | DenisNF123 | se | 13.07 | (24, 27) | SRX3029306 |
| *D. sechellia* | DenisNF13 | se | 24.12 | (24, 27) | SRX3029313 |
| *D. sechellia* | DenisNF134 | se | 14.45 | (24, 27) | SRX3029302 |
| *D. sechellia* | DenisNF155 | se | 15.84 | (24, 27) | SRX3029321 |
| *D. sechellia* | DenisNF66 | se | 27.16 | (24, 27) | SRX3029312 |
| *D. sechellia* | DenisNoni10 | se | 14.63 | (24, 27) | SRX3029320 |
| *D. sechellia* | DenisNoni101 | se | 25.07 | (24, 27) | SRX3029278 |
| *D. sechellia* | DenisNoni60 | se | 19.35 | (24, 27) | SRX3029318 |
| *D. sechellia* | LD11_sech | pe | 44.37 | (24, 27) | SRX3029282 |
| *D. sechellia* | LD12 | pe | 37.29 | (24, 27) | SRX3029284 |
| *D. sechellia* | LD13 | pe | 45.52 | (24, 27) | SRX3029289 |
| *D. sechellia* | LD14 | pe | 34.93 | (24, 27) | SRX3029288 |
| *D. sechellia* | LD15 | pe | 49.06 | (24, 27) | SRX3029362 |
| *D. sechellia* | LD16 | pe | 40.63 | (24, 27) | SRX3029290 |
| *D. sechellia* | LD8 | pe | 41.54 | (24, 27) | SRX3029364 |
| *D. sechellia* | mariane_1 | pe | 49.51 | (24, 27) | SRX3029287 |
| *D. sechellia* | maria_3 | pe | 39.82 | (24, 27) | SRX3029376 |
| *D. sechellia* | PNF10 | pe | 34.48 | (24, 27) | SRX3029280 |
| *D. sechellia* | PNF11 | pe | 39.12 | (24, 27) | SRX3029279 |
| *D. sechellia* | PNF3 | pe | 46.24 | (24, 27) | SRX3029272 |
| *D. sechellia* | PNF4 | pe | 32.94 | (24, 27) | SRX3029363 |
| *D. sechellia* | PNF5 | pe | 41.99 | (24, 27) | SRX3029273 |
| *D. sechellia* | PNF7 | pe | 28.88 | (24, 27) | SRX3029271 |
| *D. sechellia* | PNF8 | pe | 55.93 | (24, 27) | SRX3029274 |
| *D. simulans* | Bioko_cascade_1 | pe | 38.1 | (28, 44) | SRX7116491 |
| *D. simulans* | Bioko_H1 | pe | 38.4 | (28, 44) | SRX7116492 |
| *D. simulans* | Bioko_LB1 | pe | 38.26 | (28, 44) | SRX7116493 |
| *D. simulans* | Bioko_Riaba_9 | pe | 32.7 | (28, 44) | SRX7116489 |
| *D. simulans* | Bioko_Riaba_mixed | pe | 32.53 | (28, 44) | SRX7116490 |
| *D. simulans* | Kib32 | se,pe | 52.05 | (21, 22) | SRR580348, SRR580347, SRR580350, SRR580349 |
| *D. simulans* | MD06 | pe | 128.34 | (21, 22) | SRX6458044 |
| *D. simulans* | MD105 | pe | 101.25 | (21, 22) | SRX6458047 |
| *D. simulans* | MD106 | pe | 104.11 | (21, 22) | SRX6458049 |
| *D. simulans* | MD15 | pe | 115.94 | (21, 22) | SRX6458052 |
| *D. simulans* | MD199 | pe | 128.89 | (21, 22) | SRX8034374 |
| *D. simulans* | MD221 | pe | 115.61 | (21, 22) | SRX6458058 |
| *D. simulans* | MD233 | pe | 139.25 | (21, 22) | SRX6458061 |
| *D. simulans* | MD251 | pe | 133.51 | (21, 22) | SRX6458064 |
| *D. simulans* | MD63 | pe | 64.75 | (21, 22) | SRX6458067 |
| *D. simulans* | MD73 | pe | 130.06 | (21, 22) | SRX6458070 |
| *D. simulans* | NS05 | pe | 135.43 | (21, 22) | SRX6458073 |
| *D. simulans* | NS113 | pe | 125.58 | (21, 22) | SRX6458076 |
| *D. simulans* | NS137 | pe | 111.69 | (21, 22) | SRX6458078 |
| *D. simulans* | NS33 | pe | 125 | (21, 22) | SRX6458081 |
| *D. simulans* | NS39 | pe | 136.06 | (21, 22) | SRX6458084 |
| *D. simulans* | NS40 | pe | 136.32 | (21, 22) | SRX6458087 |
| *D. simulans* | NS50 | pe | 131.23 | (21, 22) | SRX6458090 |
| *D. simulans* | NS67 | pe | 139.1 | (21, 22) | SRX6458093 |
| *D. simulans* | NS78 | pe | 136.03 | (21, 22) | SRX6458096 |
| *D. simulans* | NS79 | pe | 135.12 | (21, 22) | SRX6458099 |
| *D. simulans* | tsimbazazaa | pe | 38.11 |  | SRR869580, SRR869579 |
| *D. simulans* | w501 | pe | 26.35 | (34) | SRR520350 |
| *D. teissieri* | Balancha_1 | pe | 30.37 | (23) | SRX3029331 |
| *D. teissieri* | Bata2 | se | 20.7 | (23) | SRX3029370 |
| *D. teissieri* | Bata8 | se | 18.56 | (23) | SRX3029369 |
| *D. teissieri* | cascade_2_1 | pe | 29.2 | (23) | SRX3029323 |
| *D. teissieri* | cascade_2_2 | pe | 33.88 | (23) | SRX3029330 |
| *D. teissieri* | cascade_2_4 | pe | 26.91 | (23) | SRX3029324 |
| *D. teissieri* | cascade_4_1 | pe | 27.07 | (23) | SRX3029328 |
| *D. teissieri* | cascade_4_2 | pe | 39.54 | (23) | SRX3029374 |
| *D. teissieri* | cascade_4_3 | pe | 23.26 | (23) | SRX3029329 |
| *D. teissieri* | House_Bioko | pe | 35.7 | (23) | SRX3029325 |
| *D. teissieri* | La_Lope_Gabon | pe | 36.6 | (23) | SRX3029375 |
| *D. teissieri* | Selinda | pe | 27.74 | (23) | SRX3029326 |
| *D. yakuba* | 1_19 | se | 18.51 | (23) | SRX3029345 |
| *D. yakuba* | 1_5 | se | 19.27 | (23) | SRX3518253 |
| *D. yakuba* | 1_6 | se | 20.16 | (23) | SRX3029348 |
| *D. yakuba* | 1_7 | se | 22.01 | (23) | SRX3029297 |
| *D. yakuba* | 2_11 | se | 19.51 | (23) | SRX3029350 |
| *D. yakuba* | 2_14 | se | 19.15 | (23) | SRX3029347 |
| *D. yakuba* | 2_6 | se | 23.43 | (23) | SRX3029353 |
| *D. yakuba* | 2_8 | se | 20.38 | (23) | SRX3029343 |
| *D. yakuba* | 3_16 | se | 19.82 | (23) | SRX3029344 |
| *D. yakuba* | 3_2 | se | 21.89 | (23) | SRX3029356 |
| *D. yakuba* | 3_23 | se | 22.11 | (23) | SRX3029291 |
| *D. yakuba* | 4_21 | se | 22.44 | (23) | SRX3029299 |
| *D. yakuba* | Abidjan_12 | se | 23.79 | (23) | SRX3029360 |
| *D. yakuba* | Airport_16_5 | se | 20.11 | (23) | SRX3029355 |
| *D. yakuba* | Anton_1_Principe | se | 19.54 | (23) | SRX3029295 |
| *D. yakuba* | Anton_2_Principe | se | 21.38 | (23) | SRX3029361 |
| *D. yakuba* | BAR_1000_2 | se | 21.23 | (23) | SRX3029367 |
| *D. yakuba* | BIOKO_NE_4_6 | se | 17.17 | (23) | SRX3029351 |
| *D. yakuba* | Bosu_1235_14 | se | 17.22 | (23) | SRX3029300 |
| *D. yakuba* | Cascade_18 | se | 23.96 | (23) | SRX3029354 |
| *D. yakuba* | Cascade_19_16 | se | 16.5 | (23) | SRX3029342 |
| *D. yakuba* | Cascade_21 | se | 20.59 | (23) | SRX3029368 |
| *D. yakuba* | Cascade_SN6_1 | se | 18.85 | (23) | SRX3029358 |
| *D. yakuba* | COST_1235_2 | se | 17.69 | (23) | SRX3029296 |
| *D. yakuba* | COST_1235_3 | se | 15.42 | (23) | SRX3029349 |
| *D. yakuba* | CY01A | pe | 196.72 | (21, 22) | SRX6457979, SRX6457980 , SRX6457981, SRX6457982 |
| *D. yakuba* | CY02B5 | pe | 69.98 | (21, 22) | SRX6457983, SRX6457984, SRX6457985 |
| *D. yakuba* | CY04B | pe | 157.94 | (21, 22) | SRX6457986, SRX6457987, SRX6457988, SRX6457989 |
| *D. yakuba* | CY08A | pe | 75.04 | (21, 22) | SRX6457990, SRX6457991, SRX6457992 |
| *D. yakuba* | CY13A | pe | 72.72 | (21, 22) | SRX6457990, SRX6457991, SRX6457992 |
| *D. yakuba* | CY17C | pe | 193.88 | (21, 22) | SRX6457996, SRX6457997, SRX6457998, SRX6457999 |
| *D. yakuba* | CY20A | pe | 183.65 | (21, 22) | SRX6458000, SRX6458001, SRX6458002 |
| *D. yakuba* | CY21B3 | pe | 173.17 | (21, 22) | SRX6458003, SRX6458004, SRX6458005 |
| *D. yakuba* | CY22B | pe | 69.84 | (21, 22) | SRX6458006, SRX6458007, SRX6458008 |
| *D. yakuba* | CY28 | pe | 110.16 | (21, 22) | SRX6458009, SRX6458010, SRX6458011 |
| *D. yakuba* | Montecafe_17_17 | se | 19.97 | (23) | SRX3029359 |
| *D. yakuba* | NY141 | pe | 143.54 | (23) | SRX6458037, SRX6458038, SRX6458039 |
| *D. yakuba* | NY42 | pe | 118.02 | (23) | SRX6458040, SRX6458041 |
| *D. yakuba* | NY48 | pe | 84.99 | (21, 22) | SRX6458013, SRX6458014, SRX6458015 |
| *D. yakuba* | NY56 | pe | 88.65 | (21, 22) | SRX6458016, SRX6458017, SRX6458018 |
| *D. yakuba* | NY62 | pe | 94.51 | (21, 22) | SRX6458019, SRX6458020, SRX6458021 |
| *D. yakuba* | NY65 | pe | 91.46 | (21, 22) | SRX6458022, SRX6458023, SRX6458024 |
| *D. yakuba* | NY66 | pe | 148.65 | (21, 22) | SRX6458025, SRX6458026, SRX6458027 |
| *D. yakuba* | NY73 | pe | 92.08 | (21, 22) | SRX6458028, SRX6458029, SRX6458030 |
| *D. yakuba* | NY81 | pe | 148.42 | (21, 22) | SRX6458031, SRX6458032, SRX6458033 |
| *D. yakuba* | NY85 | pe | 99.03 | (21, 22) | SRX6458034, SRX6458035, SRX6458036 |
| *D. yakuba* | OBAT_1200_5 | se | 22.7 | (23) | SRX3029357 |
| *D. yakuba* | SanTome_city_14_26 | se | 22.75 | (23) | SRX3029372 |
| *D. yakuba* | SA_3 | se | 18.64 | (23) | SRX3029294 |
| *D. yakuba* | SJ14 | se | 15.77 | (23) | SRX3029293 |
| *D. yakuba* | SJ4 | se | 25.82 | (23) | SRX3029292 |
| *D. yakuba* | SJ7 | se | 19.51 | (23) | SRX3029352 |
| *D. yakuba* | SJ_1 | se | 21.35 | (23) | SRX3029371 |
| *D. yakuba* | SN7 | se | 23.66 | (23) | SRX3029366 |
| *D. yakuba* | SN_Cascade_22 | se | 21.78 | (23) | SRX3029365 |
| *D. yakuba* | Tai_18 | se | 22.17 | (23) | SRX3029298 |

**SUPPLEMENTARY FIGURES**

**FIGURE S1. *X_sim_* and *X_mau_* cause abdominal ablations in hybrid males with *D. santomea*.** Hybrid males from the **♀***sim ×* **♂***san* and **♀***mau ×* **♂** *san* crosses show high frequency of abdominal ablations similar to those observed in **♀***mel ×* **♂** *san* hybrids (Figures 1 and 2C). Hybrid females from the same crosses show a lower frequency of ablations. The nature of the defect in these hybrid males is identical to that seen in *mel/san* hybrid males, a characteristic ablation of abdominal segments (Figure 1C).
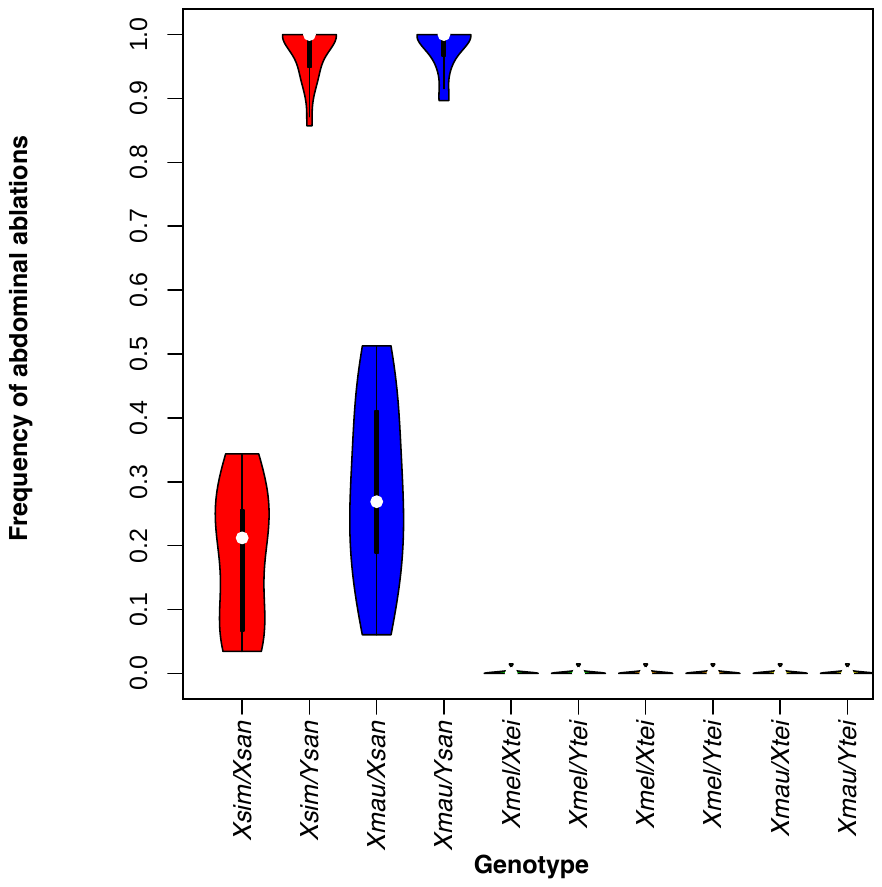

**FIGURE S2. Crossing design to assess whether of *Y*-linked pieces of the *X_mel_* chromosome cause hybrid inviability and abdominal ablations.** Blue bars represent *D. santomea* chromosomes; yellow bars represent *D. melanogaster* chromosomes. Solid colors: sex chromosomes, stripped bars: autosomes. This approach is a modified version of (12, 14, 45).

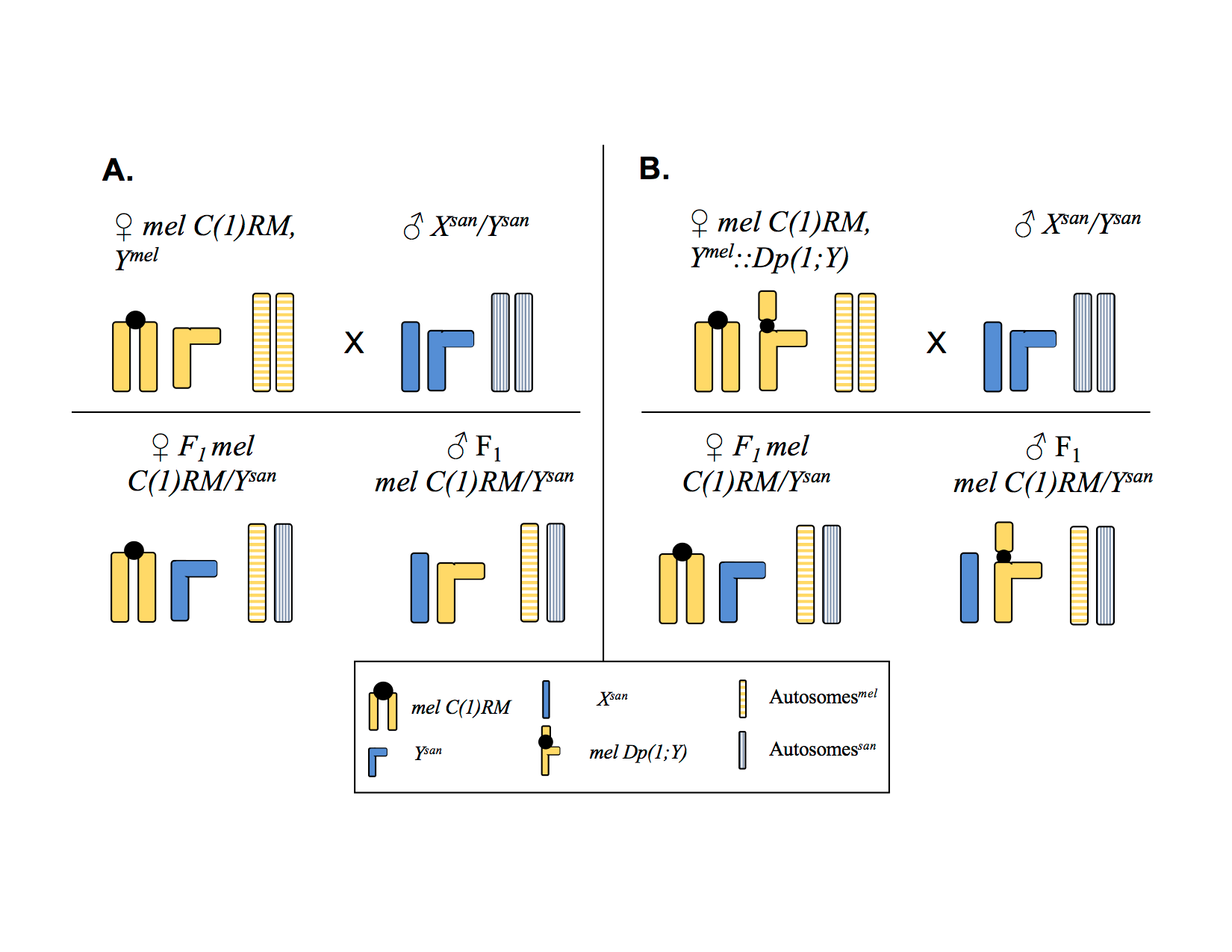

**FIGURE S3.** **Frequency of abdominal ablations caused by the *X_mel_* translocations shown in Figure 2 in *X_san_/Y_mel_* hybrid males.** Each Bloomington stock number is shown within the bar. The precise genotype of each stock is shown in Table S3.

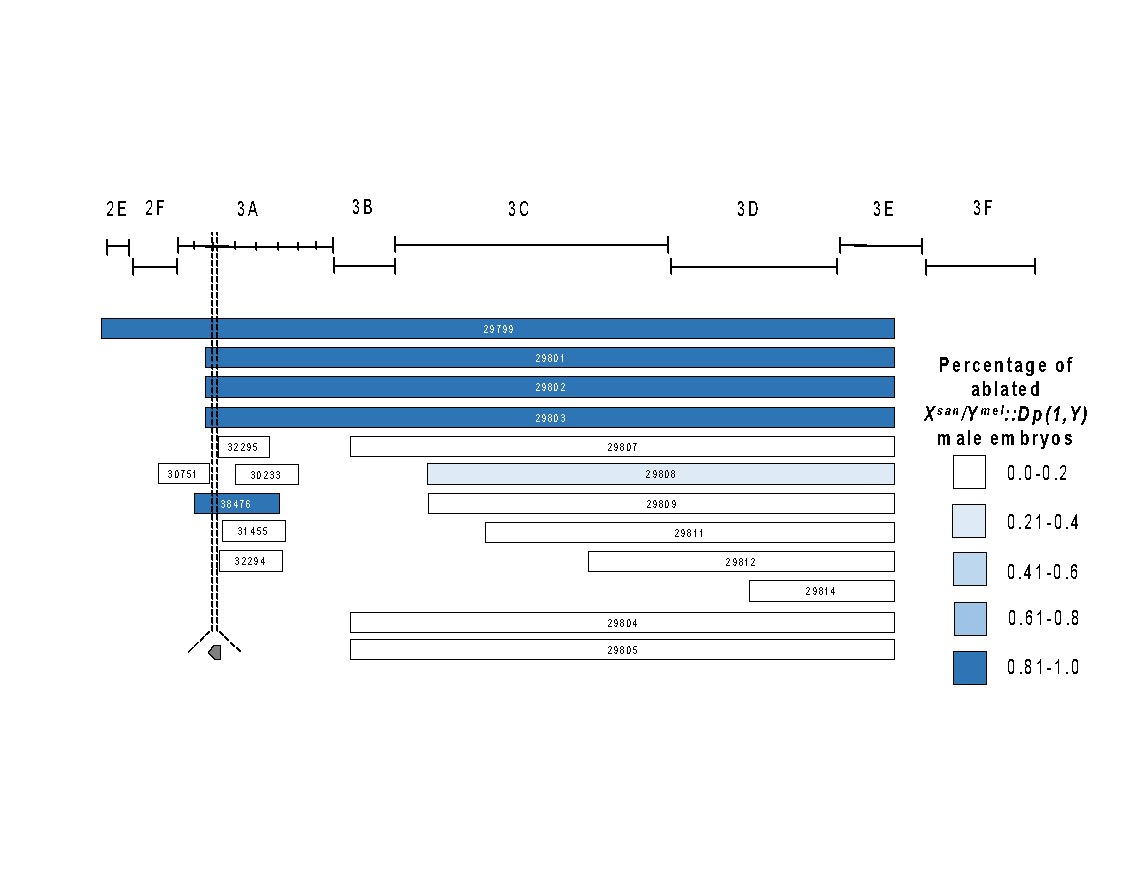

**FIGURE S4. The presence of *gt_mel_* causes HI in *mel/san* hybrids produced with all *D. santomea* lines but the magnitude of the inviability varies.** I measured frequency of abdominal ablations in hybrid *X_mel_/Y_san_* males (**A**). The magnitude of the frequency of abdominal ablations and of hybrid female inviability are correlated among lines (panel **B**; ρ = 0.1734, P = 0.0141). Boxes in the boxplot are ordinated from the lower median (left) to the highest (right). S1: Thena5; S2: sanCAR1490; S3: sanCOST1250.5; S4: sanCOST1270.1; S5: sanOBAT1200; S6: sanOBAT1200.2; S7: sanRain39; S8: sanCAR1600.3; S9: Carv2015.1; S10: Carv2015.5; S11: Carv2015.11; S12: Carv2015.16; S13: Pico1680.1; S14: Pico1659.2; S15: Pico1659.3; S16: Amelia2015.1; S17: Amelia2015.6; S18: Amelia2015.12; S19: A1200.7; S20: Rain42.

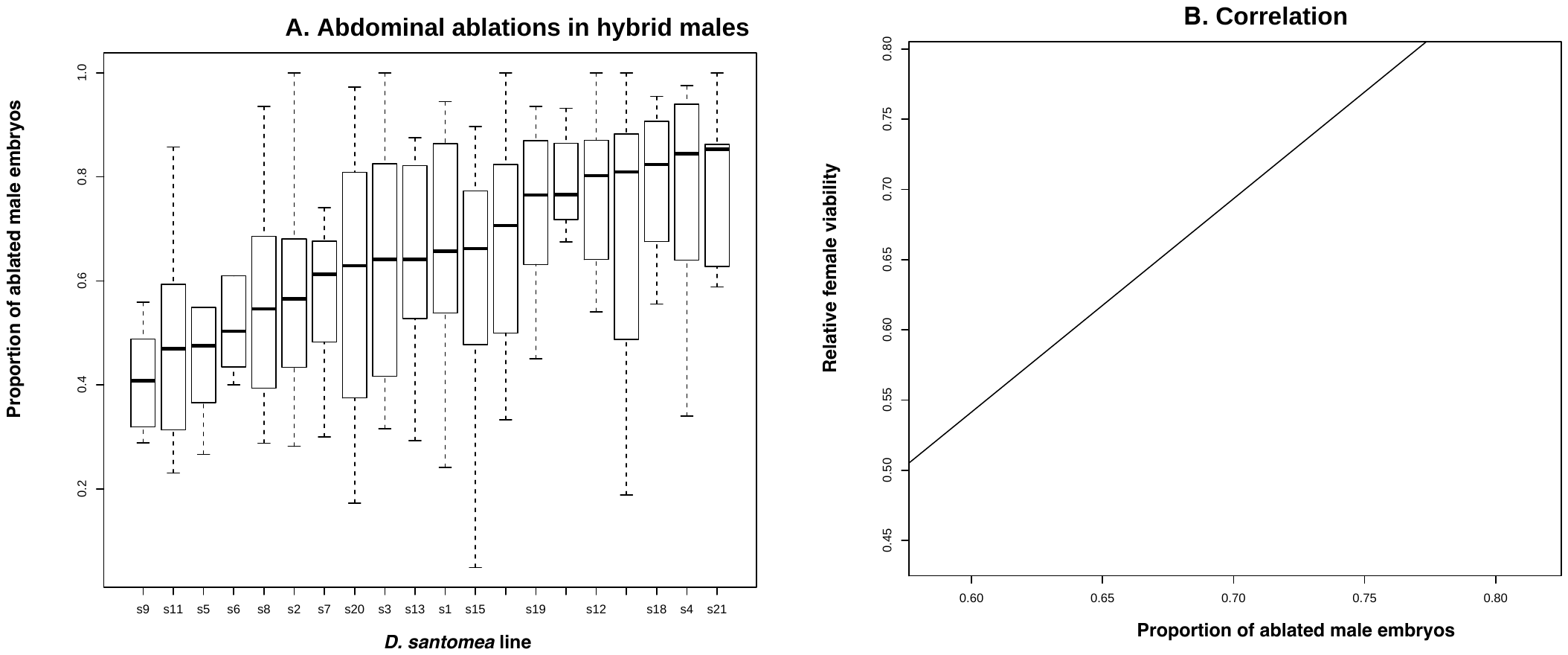

**FIGURE S5. Crosses between *D. melanogaster* females harboring *Dp(1;Y)* duplications and males from the *simulans* species group (*D. mauritiana*, and *D. simulans*) show no evidence of male embryo lethality.** *Dp(1:Y)* duplications containing *gt_mel_* cause no embryonic defects and do not cause heightened hybrid inviability. The color code is the same as used in Figure 2C. The lack of gray bars indicates that none of the duplications caused hybrid inviability in any of the crosses.

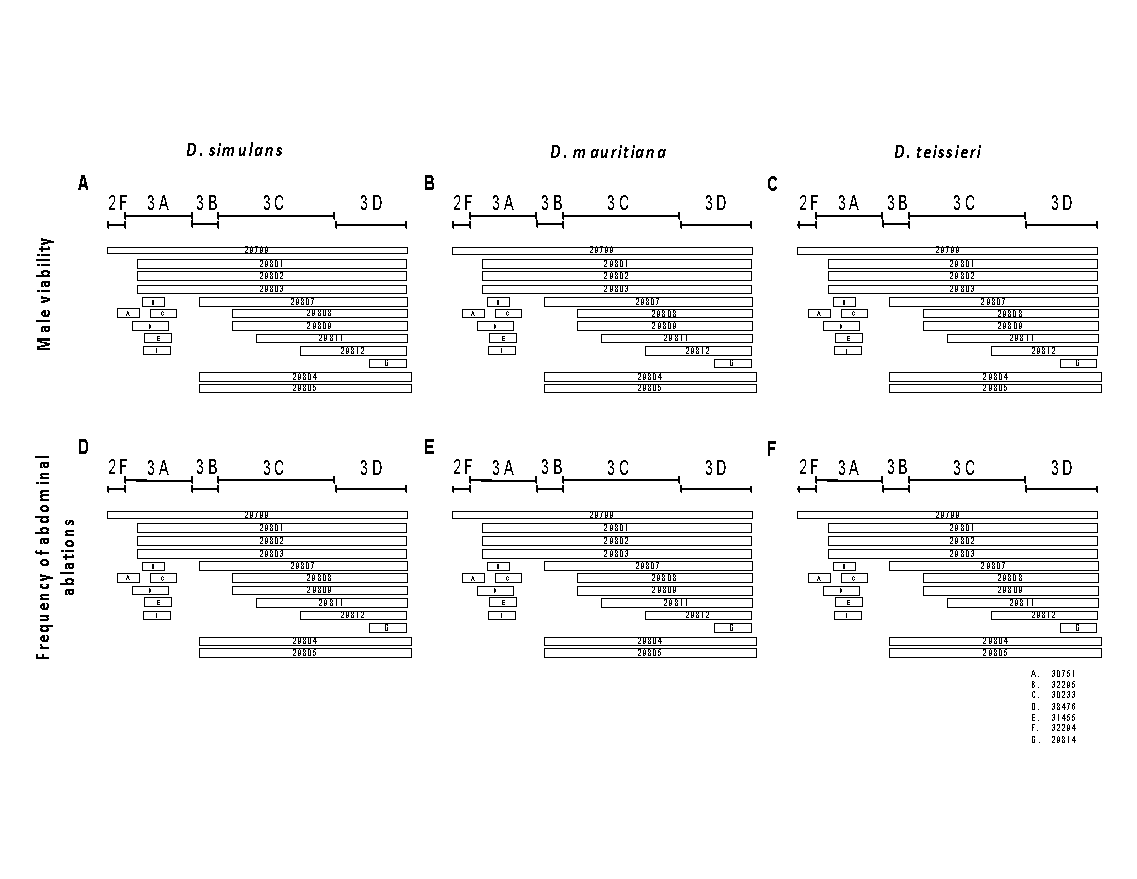

**FIGURE S6. Crosses between *D. melanogaster* females harboring *Dp(1;Y)* duplications and males from different lines of *D. melanogaster* show no evidence of male embryo lethality or abdominal ablations.** *Dp(1:Y)* duplications containing *gt^mel^* cause no embryonic defects and do not cause heightened HI. The color code is the same as used in Figure 2C. The lack of gray bars indicates that none of the duplications caused hybrid inviability in any of the crosses.

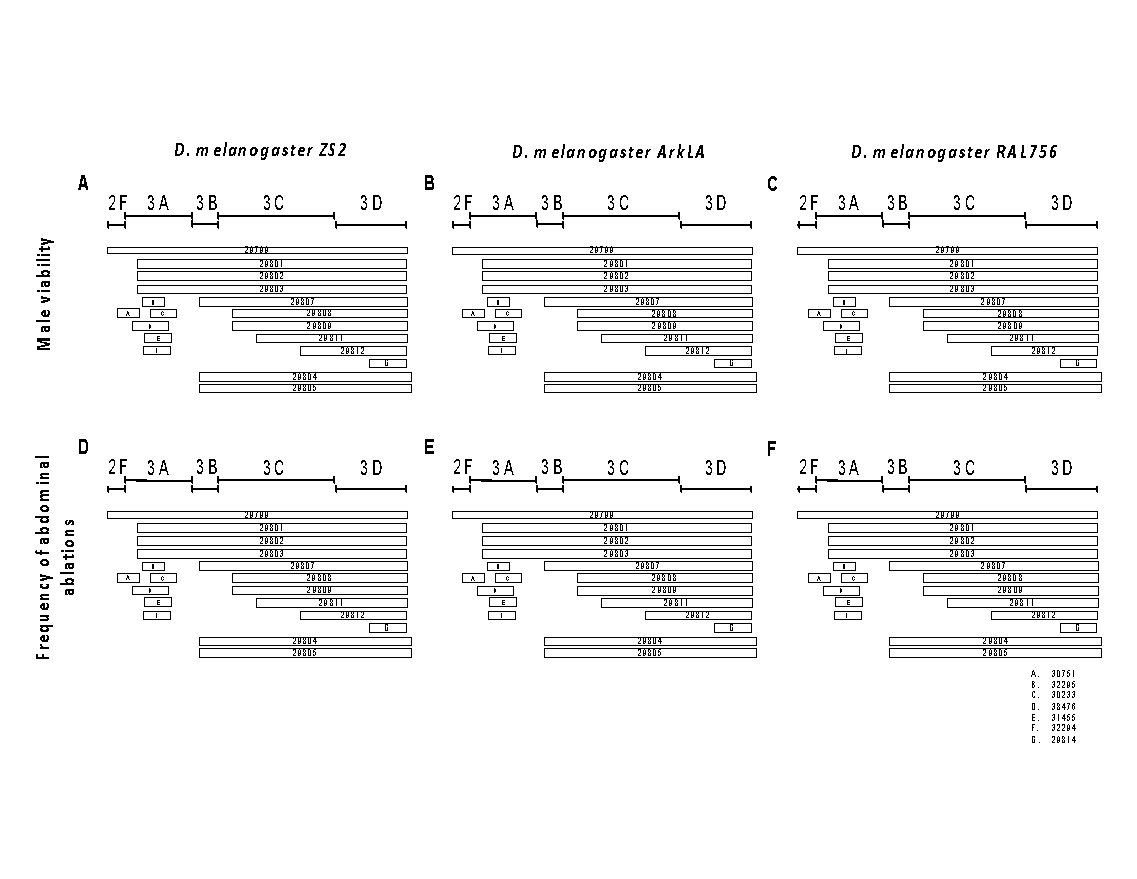

**FIGURE S7. *mel/san* hybrid females carrying a *D. melanogaster* either a balancer chromosome or a mutant (hypomorphic or null) alleles for *boi*, *trol*, or *tko* show similar levels of hybrid viability.** This is in contrast to males also carrying a *D. melanogaster* chromosome but a *gt* null allele (*gt_mel_^X11^*) which show a significant reduction in abdominal ablations (shown in Figure 2). These three genes have no effect in hybrid female viability between *D. melanogaster* and three more species (*D. teissieri* [B], *D. simulans* [C], and *D. mauritiana* [D]).

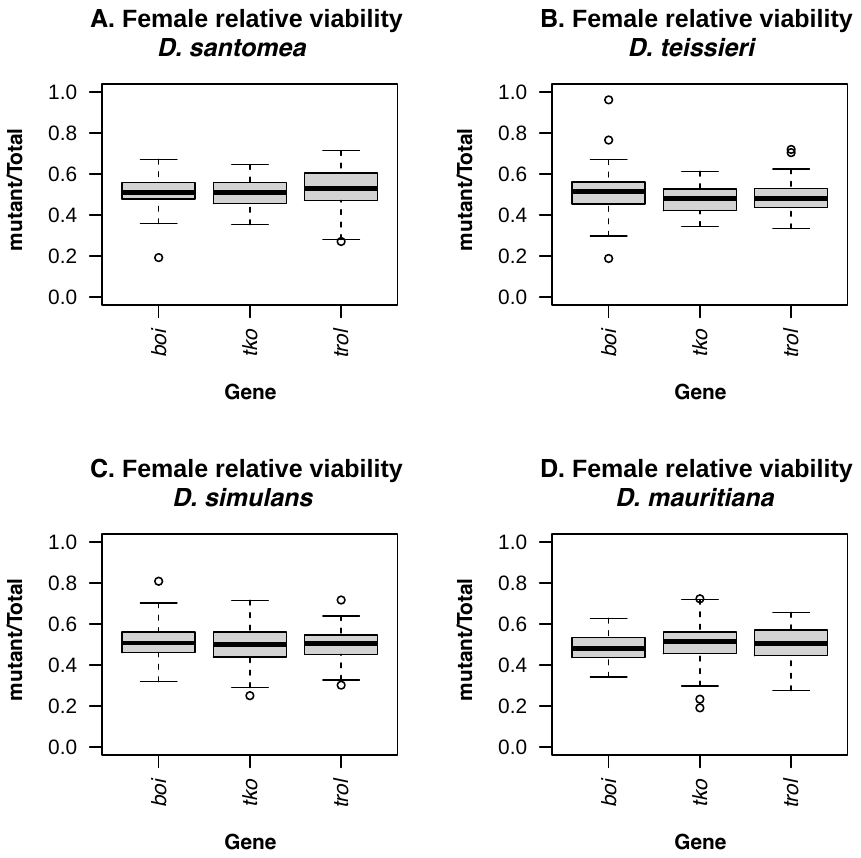

**FIGURE S8. The *X*-chromosome balancer identity has no effect on the quantification of hybrid inviability in *mel/san* hybrid females.** I found no heterogeneity in the relative viability of two different *gt* null mutants (A. *gt^X11^;* B. *Df(1)Exel6231*) when different balancer chromosomes are used (One-way ANOVA, F_6,109_= 0.694, P= 0.655). I used seven different *X*-chromosomes balancers and none of them had a major effect on the quantification of the relative frequency of viability in any of the hybrid crosses.

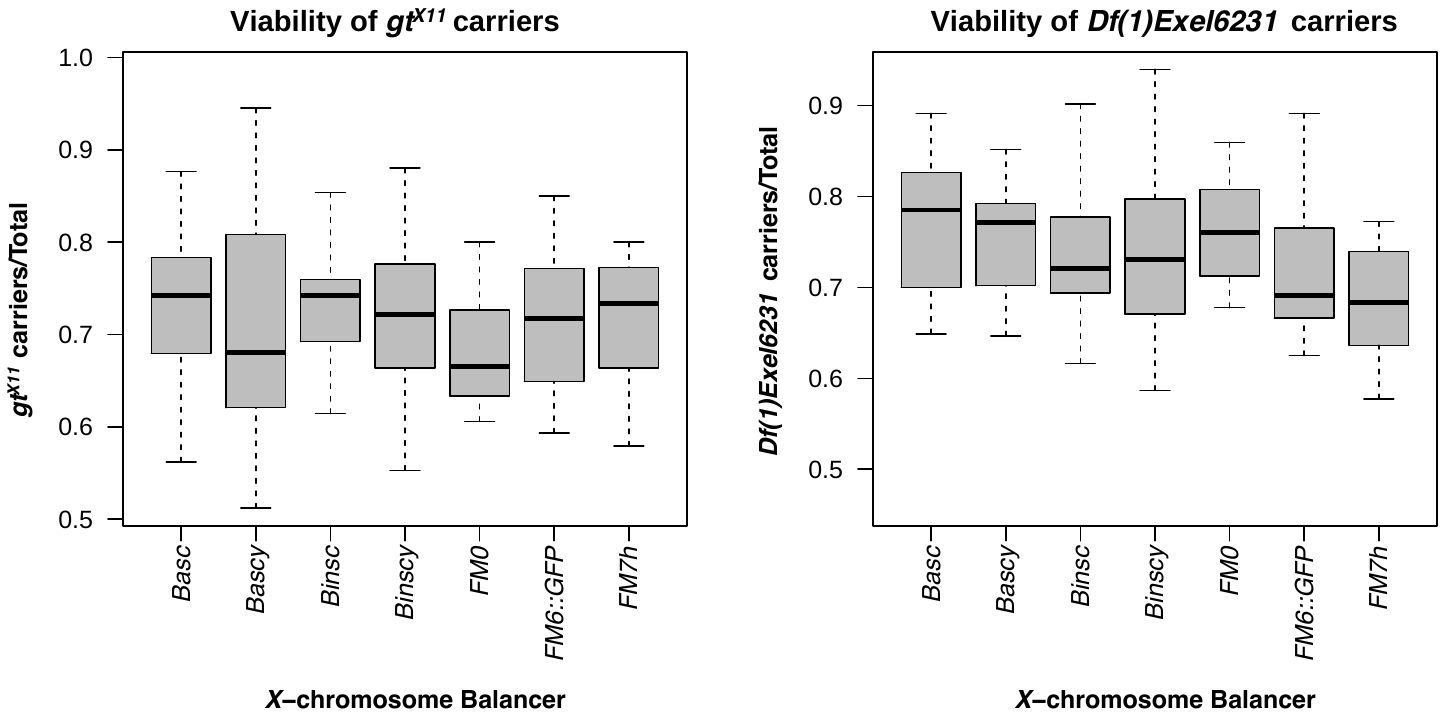

**FIGURE S9.** **Frequency of abdominal ablations in each deficiency cross shown in Figure 3A.** Deficiency chromosomes that contain a functional *gt_mel_* (e.g., 8031, 9054, and 8950) were more prone to show abdominal ablations than deficiencies that harbored other genes. Conversely, deficiency chromosomes with no functional copy of *gt_mel_* show reduced rates of abdominal ablations. The mean proportion of abdominal ablation in *X_mel_/X_san_* hybrid females is 0.420.

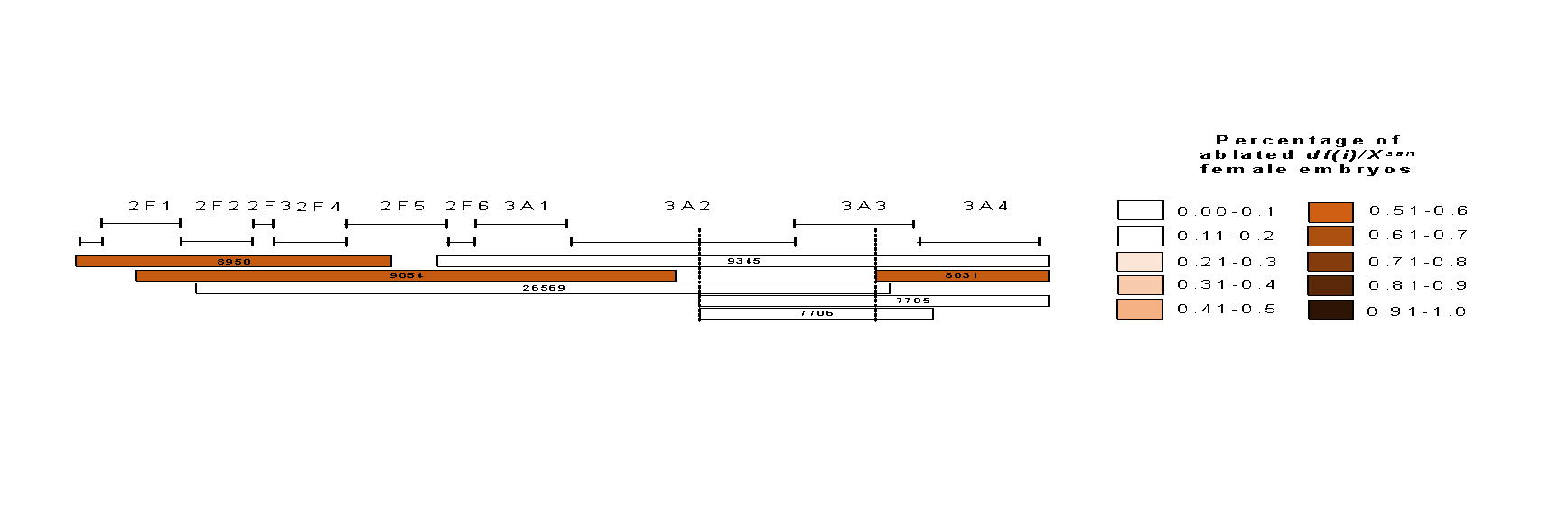

**FIGURE S10.** ***gt^mel^* has no effect on the viability of *mel/tei*, *mel/mau*, and *mel/sim* hybrid females.** I used the same deficiency chromosomes reported to detect the effect of *gt_mel_* in *mel/san* hybrid females to detect a potential effect of *gt_mel_* in hybrids between *D. melanogaster* and other species (Figure 3A). In all three cases, *df/FM7::GFP* crossed to males of each of the species led to 1:1 ratios in female progeny. The color code is the same as in Figure 3A but since no deficiency departed from the 1:1 expected (i.e., no *gt_mel_* effect on hybrid viability), there are no gray bars.

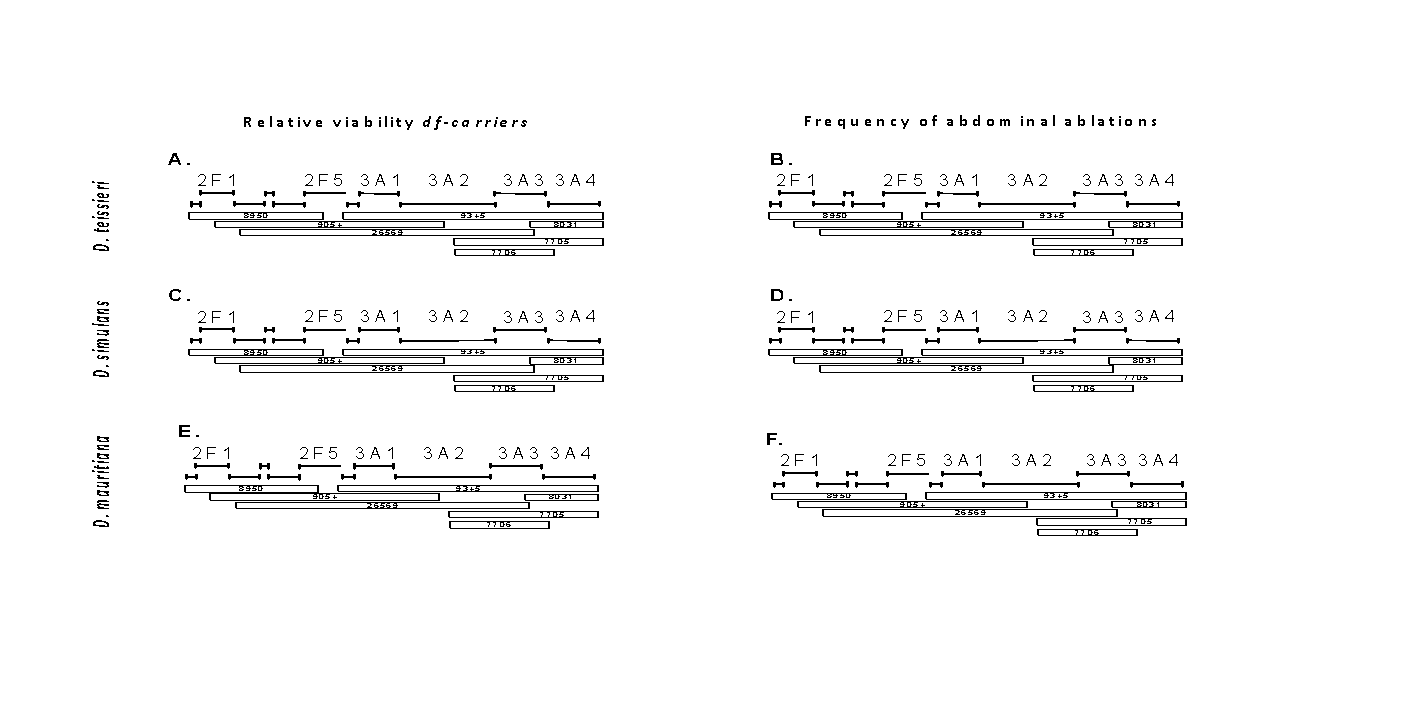

**FIGURE S11.** ***gt_mel_* has no effect on the viability of pure-species *D. melanogaster* F1 females.** I used the same deficiency chromosomes reported to detect the effect of *gt_mel_* in *mel/san* hybrid females (Figure 3A). In all three types of crosses (three isofemale lines), *df/FM7::GFP* crossed to males of each of the species led to 1:1 ratios in female progeny. The color code is the same as in Figure 3A but since no deficiency departed from the 1:1 expected (i.e., no *gt_mel_* effect on hybrid viability), there are no gray bars. No cross showed any abdominal ablation.

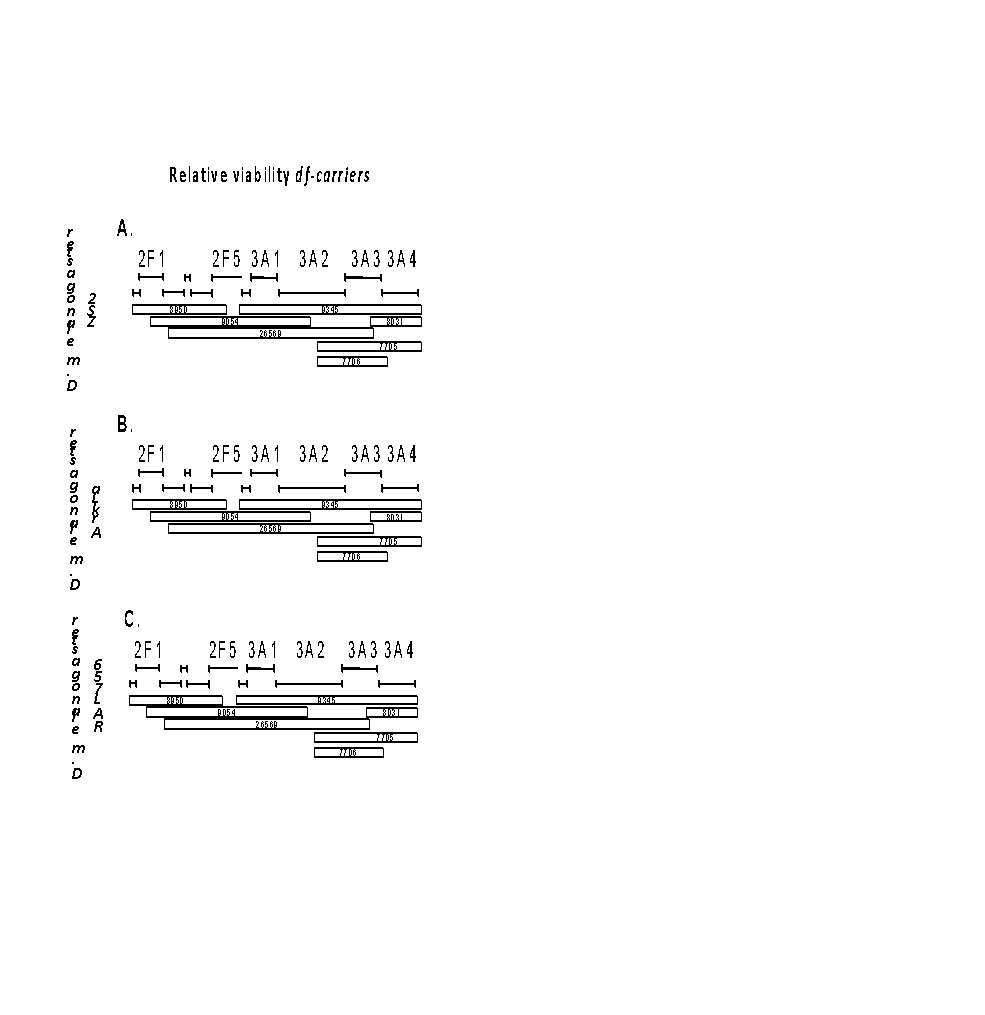

**FIGURE S12**. **Hybrid male embryos carrying a *gt_mel_^X11^* *D. melanogaster* allele show a variety of developmental defects.** *gt_mel_^X11^/Y_san_ males* are inviable and show a variety of developmental defects (A-C). A small proportion of individuals also show abdominal ablations (D).

**FIGURE S13. Introgression of the *FM7::GFP* and *gt^X11^* alleles into 200 lines of the DGRP lines.** Each bar represents a chromosome. Short bars are the *X*-chromosome while longer bars represent the autosomes (only one set of autosomes shown). The bar with dashed lines represents the *FM7::GFP* balancer. Solid yellow represent the *FM7::GFP* background. Red bars represent each of the DGRP genetic backgrounds. Dashed blue lines represent the the *gt^X11^* chromosome, while solid blue lines represent the autosomal background of the *gt^X11^* stock. **A.** The first step of the experimental design involves introgressing the *FM7::GFP* balancer into each of the 200 DGRP backgrounds. After ten generations of repeating backcrossing, I obtained both females and males that carried the balancer and the DGRP background. **B.** Males from the cross shown in **A** (carriers of the *FM7::GFP* balancer) were crossed the *gt^X11^/FM7::GFP* females. This cross produces females that carry the *gt^X11^* chromosome, the *FM7::GFP* balancer and a mixed genetic background. These females were crossed to males that had the DGRP autosomal background and a *FM7::GFP* balancer (**C**). I repeated this backcrossing approach for ten generations. After, 11 generations I obtained *gt^X11^/FM7::GFP* females with DGRP autosomal backgrounds. These females were then crossed to *D. santomea* and the percentage of ablated progeny were scored.

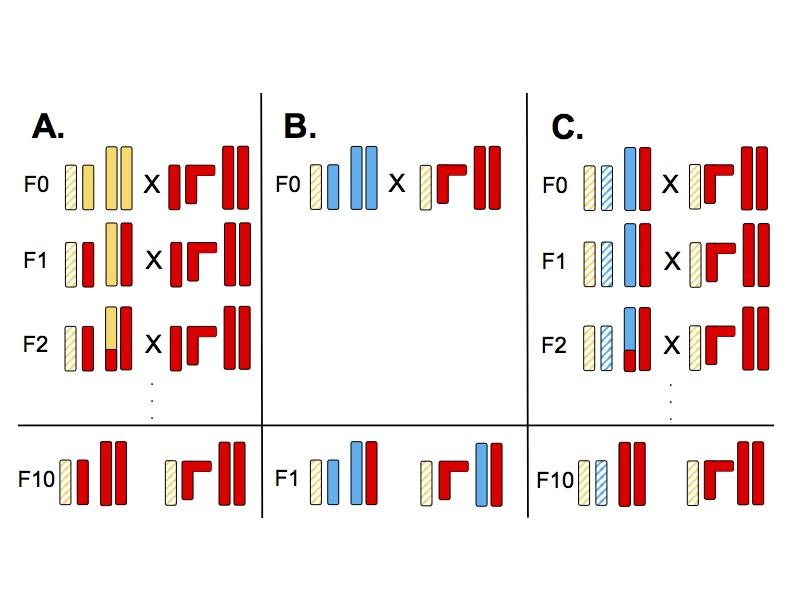

**FIGURE S14**. ***gt* alleles from six different species in the *melanogaster* species complex**. I found no major differences at the coding portion of *gt* between the *melanogaster* species supercomplex (*D. melanogaster*, *D. simulans*, and *D. sechellia*) and the *D. santomea/D. yakuba* species pair is the structure of poly-glutamine repeats. melgt: *D. melanogaster*, simgt: *D. simulans*, sechgt: *D. sechellia*, sangt: *D. santomea*, eregt: *D. erecta*. Asterisks show residues that are conserved across the whole group.

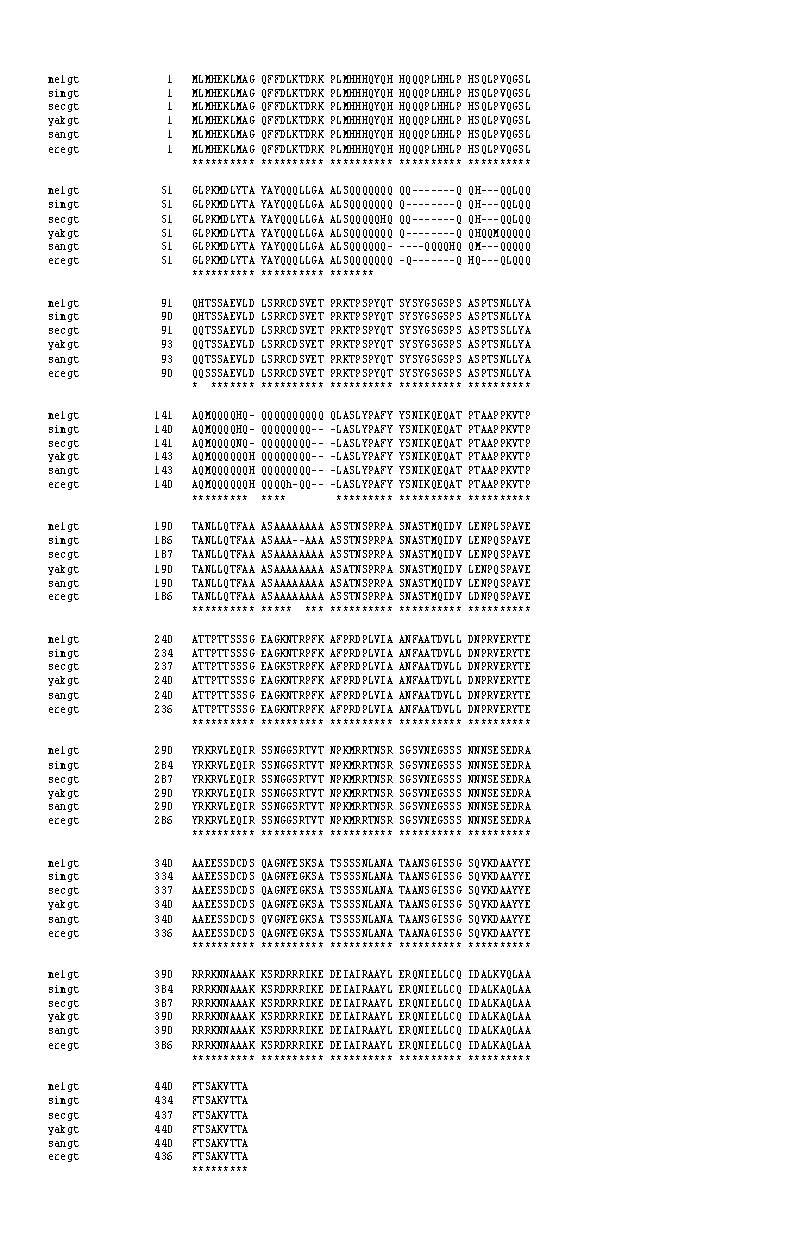

**FIGURE S15. Maximum-likelihood ancestral sequence reconstruction of GT protein, excluding polyQ.** All ancestral sites could be reconstructed with high confidence (posterior probability > 0.98), except for the two polyQ tracks. Aminoacid positions are all based on the *D. melanogaster GT* protein.

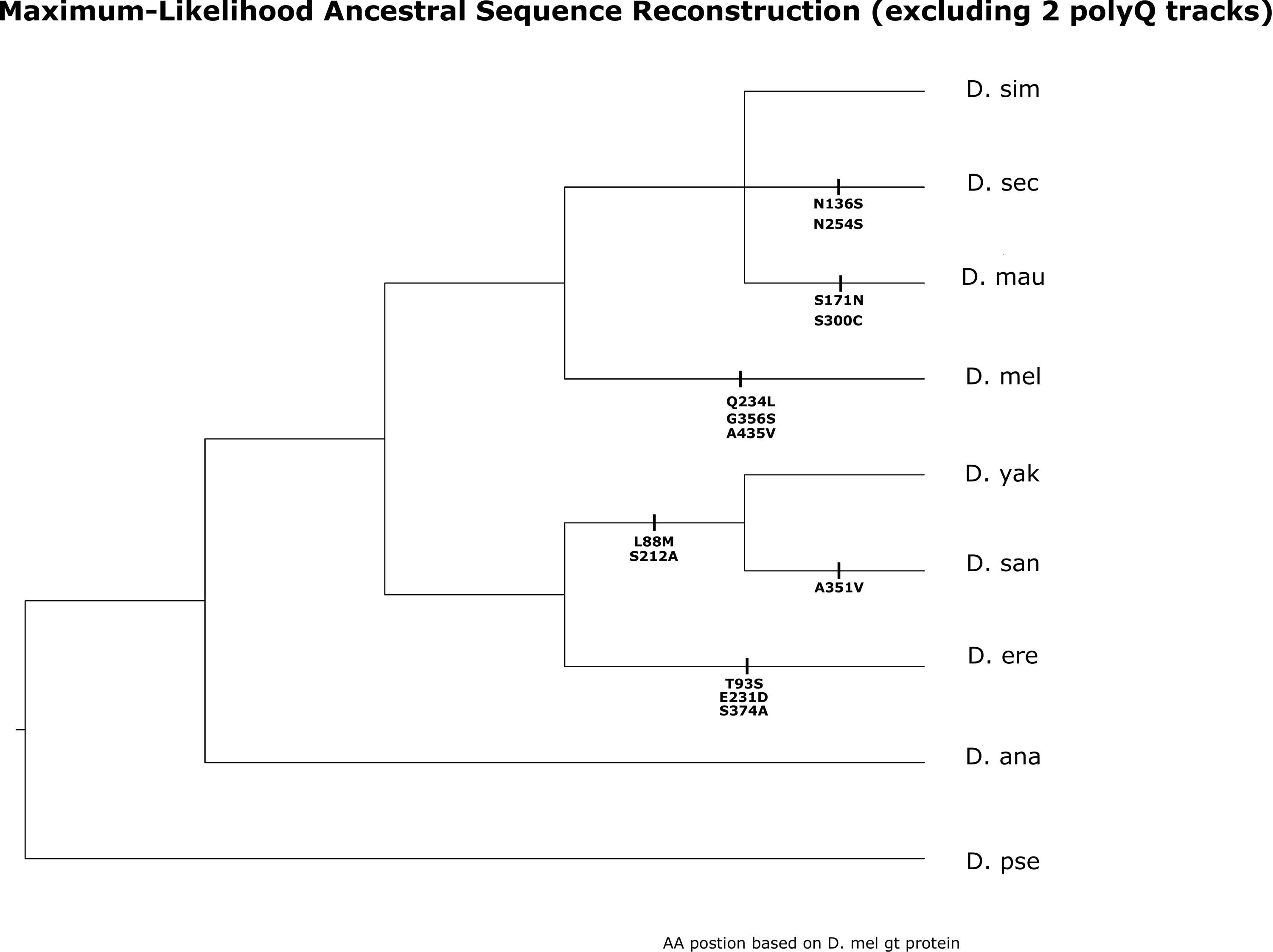

**FIGURE S16.** **The polyglutamine repeats (polyQ) show differences among the *melanogaster* subspecies complex species.** Maximum-parsimony ancestral sequence reconstruction of 2 polyQ tracks of gt protein. The conserved H within polyQ tracks helps to delineate both polyQ tracks to two parts. Due to the lack of proper substitution models, this maximum-parsimony based ancestral reconstruction for polyQ tracks may subject to bias due to alignment error, and arbitrary choice of polyQ unit.

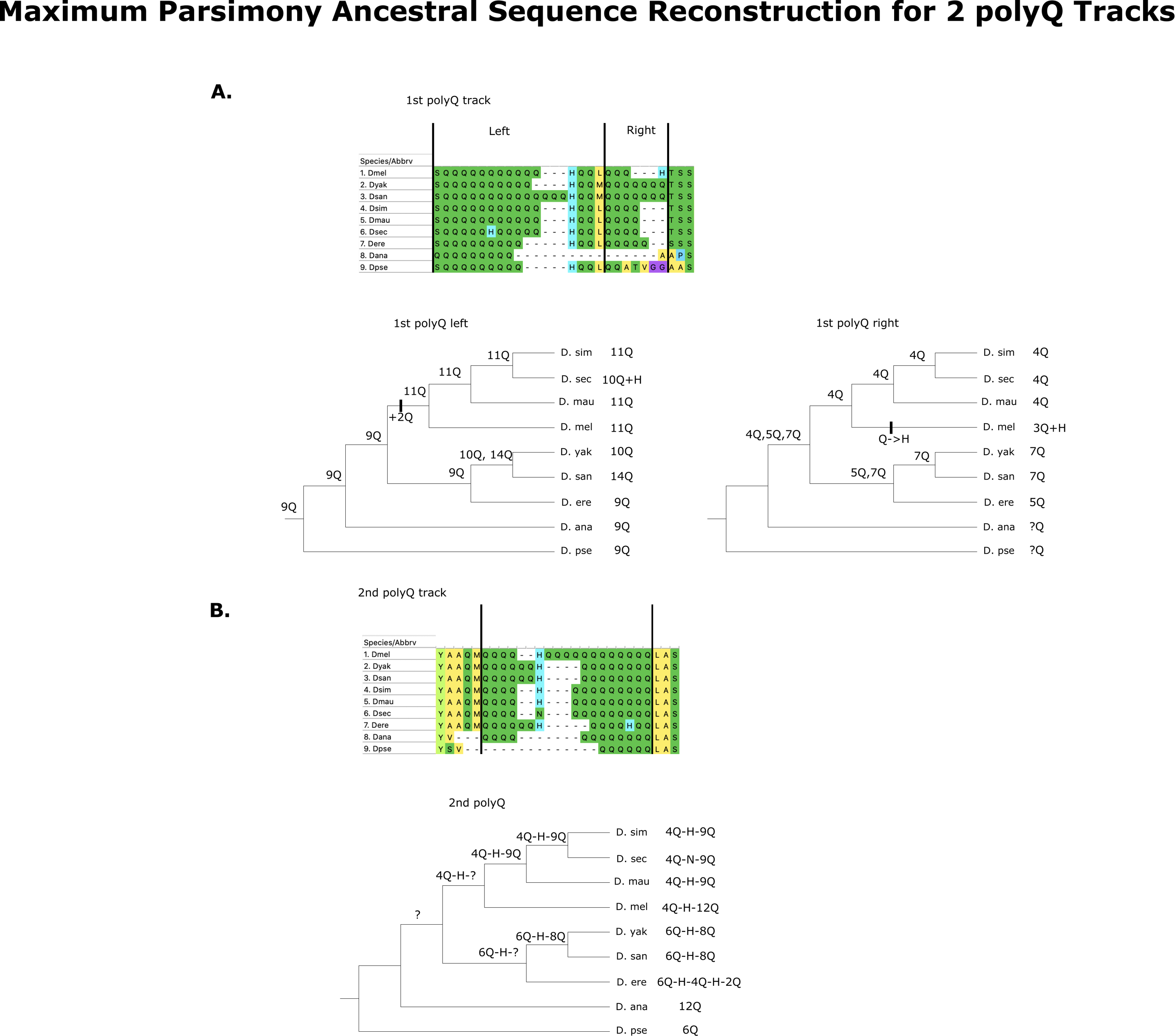

**FIGURE S17. *tll* alleles from six different species in the *melanogaster* species complex**. I found no major differences at the coding portion of *tll* between the *melanogaster* species supercomplex (*D. melanogaster*, *D. simulans*, and *D. sechellia*) and the *D. santomea/D. yakuba* species pair. tll_mel: *D. melanogaster*, tll_sim: *D. simulans*, tll_mau: *D. mauritiana*, tll_ere: *D. erecta*, tll_tei: *D. teissieri,* tll_san: *D. santomea*, tll_yak: *D. yakuba*. Asterisks show residues that are conserved across the whole group.

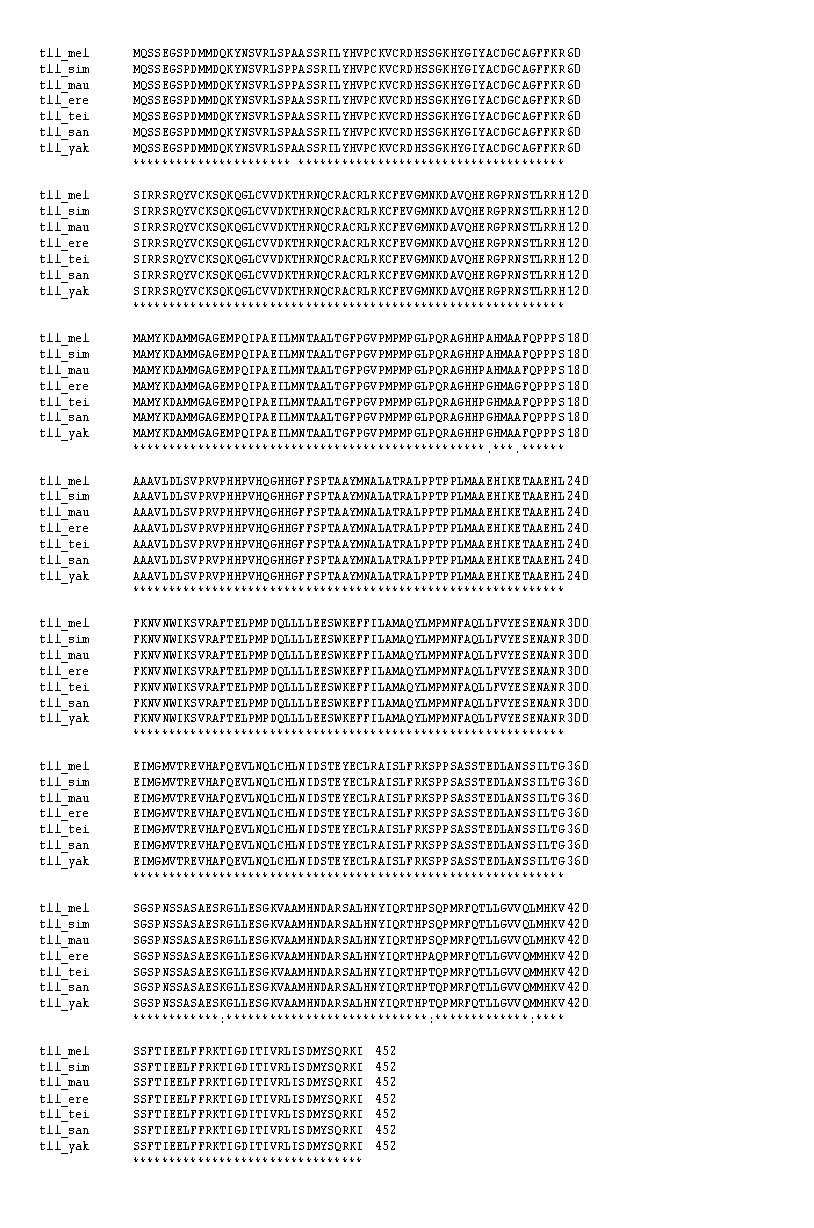

**FIGURE S18.** **Embryonic hybrid inviability does not occur in *mel/tei* hybrids.** No line of *D. teissieri* showed either embryonic inviability or abdominal ablations when crossed with *D. melanogaster*, *D. simulans*, or *D. mauritiana* females. The vast majority of assays revealed no dead embryos. **A.** *mel/tei* male hybrids. **B.** *sim/tei* male hybrids. **C.** *mau/tei* male hybrids. **D.** *mel/tei* female hybrids. **E.** *sim/tei* female hybrids. **F.** *mau/tei* female hybrids. tei1: Balancha_1; tei2: Balancha_2, tei3: Balancha_3, tei4: House_Bioko_0, tei5: House_Bioko_1, tei6: House_Bioko_2, tei7: La_Lope_Gabon, tei8: Selinda, tei9: Zimbabwe, tei10: cascade_2_1, tei11: cascade_2_2; tei12: cascade_2_4, tei13: cascade_4_1, tei14: cascade_4_2, tei15: cascade_4_3, tei16: cascade_4_4, tei17: cascade_4_5, tei18: cascade_4_6, tei19: Bata_2, tei20: Bata_8.

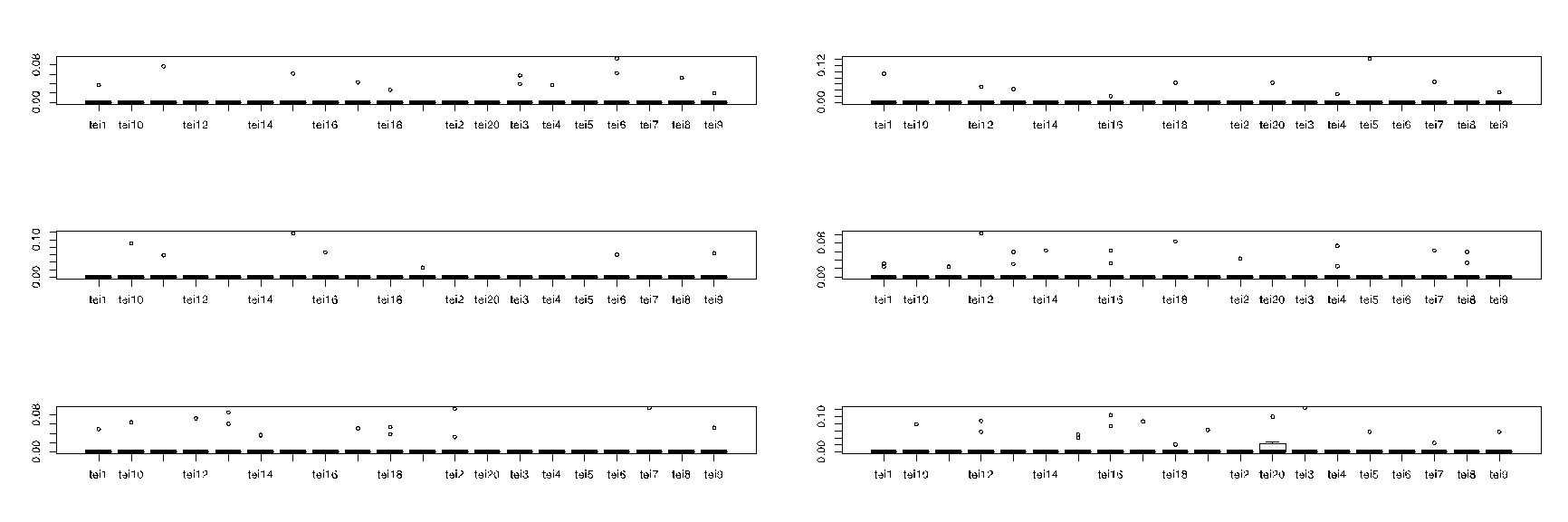

**FIGURE S19**. **Divergence in *giant* in the *melanogaster* species subgroup.** **A**. The evolutionary timing of *gt^mel^* and its interactor leading to HI. The *gt* allele responsible for HI in hybrids with *D. santomea* evolved before *mel, sim, and mau* had a common ancestor (Blue branch). At least one of the interactors of *gt^mel^* is not shared with *D.* *teissieri* which indicates that such element must have evolved after the last common ancestor of *D. santomea* and *D. teissieri* speciated (Red branch).

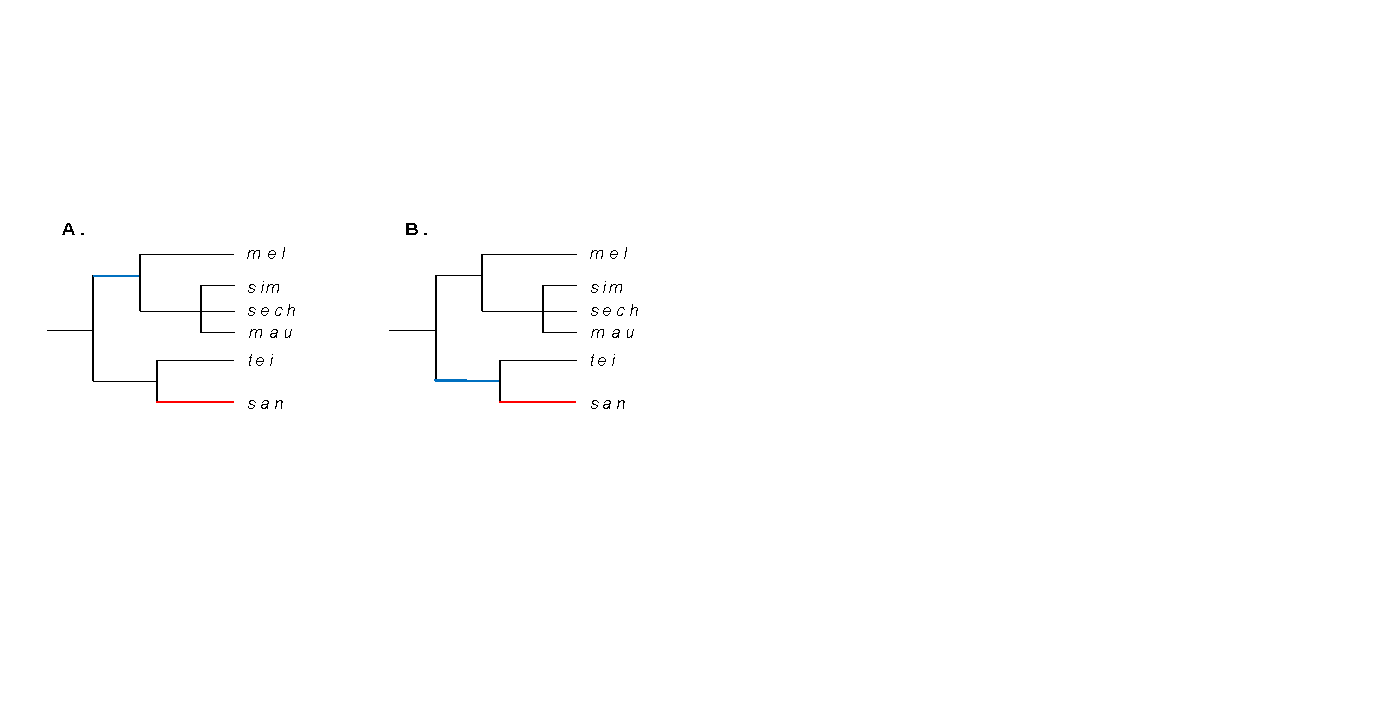

**FIGURE S20. Experimental design to generated a GFP-mediated disruption of *tll_mel_*.**

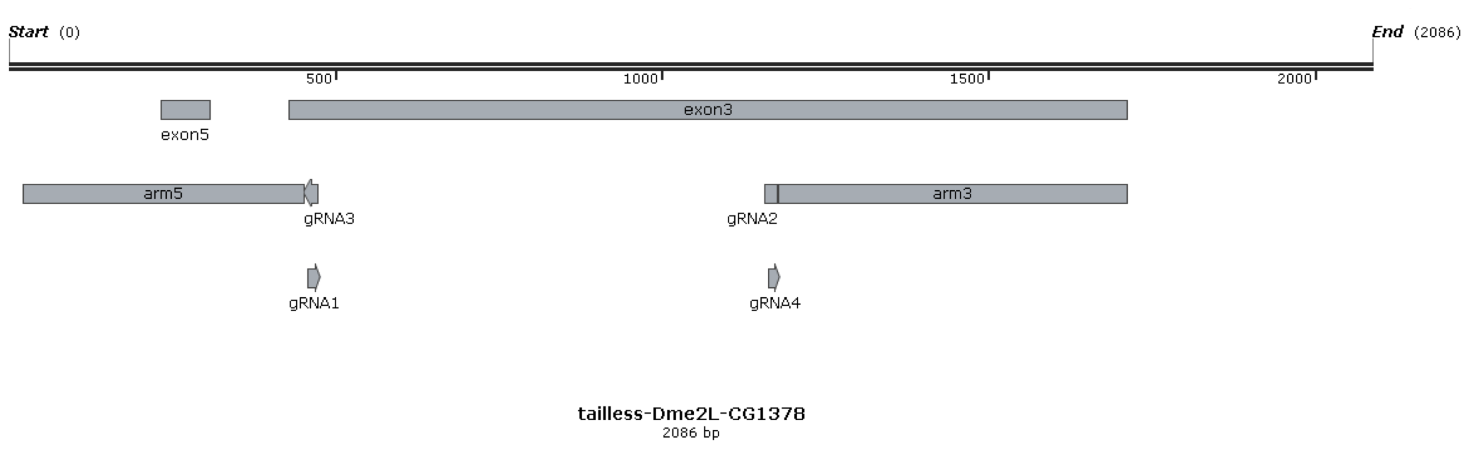

**SI REFERENCES**

1.	D. L. Lindsley, G. G. Zimm, *The genome of Drosophila melanogaster* (1992).

2.	W. Chang, D. R. Matute, M. Kreitman, Rapid evolution of the functionally conserved gap gene giant in Drosophila. *bioRxiv*, 2021.07.08.451553 (2021).

3.	D. G. Gibson, *et al.*, Enzymatic assembly of DNA molecules up to several hundred kilobases. *Nature methods* 6, 343–345 (2009).

4.	S. T. Thibault, *et al.*, A complementary transposon tool kit for Drosophila melanogaster using P and piggyBac. *Nature genetics* 36, 283–287 (2004).

5.	R Core Team, *R Development Core Team* (2016).

6.	J. A. Coyne, Genetic studies of three sibling species of Drosophila with relationship to theories of speciation. *Genetical Research* (1985) https:/doi.org/10.1017/S0016672300022643.

27.	D. R. Schrider, J. Ayroles, D. R. Matute, A. D. Kern, Supervised machine learning reveals introgressed loci in the genomes of Drosophila simulans and D. sechellia. *PLoS genetics* (2018) https:/doi.org/10.1371/journal.pgen.1007341.
